# Studies on the selectivity of the SARS-CoV-2 papain-like protease reveal the importance of the P2′ proline of the viral polyprotein

**DOI:** 10.1101/2023.07.11.548309

**Authors:** H. T. Henry Chan, Lennart Brewitz, Petra Lukacik, Claire Strain-Damerell, Martin A. Walsh, Christopher J. Schofield, Fernanda Duarte

## Abstract

The SARS-CoV-2 papain-like protease (PL^pro^) is an antiviral drug target that catalyzes the hydrolysis of the viral polyproteins pp1a/1ab, releasing the non-structural proteins (nsps) 1-3 that are essential for the coronavirus lifecycle. The LXGG↓X motif found in pp1a/1ab is crucial for recognition and cleavage by PL^pro^. We describe molecular dynamics, docking, and quantum mechanics/molecular mechanics (QM/MM) calculations to investigate how oligopeptide substrates derived from the viral polyprotein bind to PL^pro^. The results reveal how the substrate sequence affects the efficiency of PL^pro^-catalyzed hydrolysis. In particular, a proline at the P2′ position promotes catalysis, as validated by residue substitutions and mass spectrometry-based analyses. Analysis of PL^pro^ catalyzed hydrolysis of LXGG motif-containing oligopeptides derived from human proteins suggests that factors beyond the LXGG motif and the presence of a proline residue at P2′ contribute to catalytic efficiency, possibly reflecting the promiscuity of PL^pro^. The results will help in identifying PL^pro^ substrates and guiding inhibitor design.

## Introduction

The SARS-CoV-2 polyproteins ORF1a and ORF1ab are precursors of the 16 non-structural proteins (nsp1-16) that are essential for virus maturation and replication.^1^ SARS-CoV-2 employs two nucleophilic cysteine proteases to release the nsps: the nsp5 main protease (M^pro^) and the papain-like protease (PL^pro^), the latter of which is a domain within nsp3.^2^ Inhibition of M^pro^ and/or PL^pro^ hinders nsp release, leading to the stalling of viral maturation and replication.^3^ The M^pro^ inhibitors PF-07321332 (nirmatrelvir), the active pharmaceutical ingredient (API) in Paxlovid, and ensitrelvir, the API in Xocova, are approved for treating COVID-19 infections.^4–6^ However, clinical use of PL^pro^ inhibitors has not yet been approved.^3, 7, 8^

Although M^pro^ and PL^pro^ are both nucleophilic cysteine proteases, they differ in their structures, catalytic mechanisms, and substrate selectivities.^7^ M^pro^ is predominantly homodimeric and employs an active site Cys-His dyad to catalyze hydrolysis of the SARS-CoV-2 polyproteins at 11 sites, which have [P4:A/V/P/T]-[P3:X]-[P2:L/F/V]-[P1:Q]**|**[P1′:S/A/N] motifs, where “**|**” denotes the scissile amide bond and X corresponds to any proteinogenic amino acid.^3, 9, 10^ By contrast, PL^pro^ is likely monomeric and employs an active site Cys-His-Asp triad to catalyze the hydrolysis of the polyprotein at the nsp1/2, nsp2/3, and nsp3/4 sites (**Figure 1**).^11^ PL^pro^ catalyzes the hydrolysis of the polyprotein C-terminal to [P4:L]-[P3:X]-[P2:G]-[P1:G] sequences; the requirements for the P′ positions of PL^pro^ substrates have not yet been identified ([P1′:X]).^12–15^

**Figure 1:**
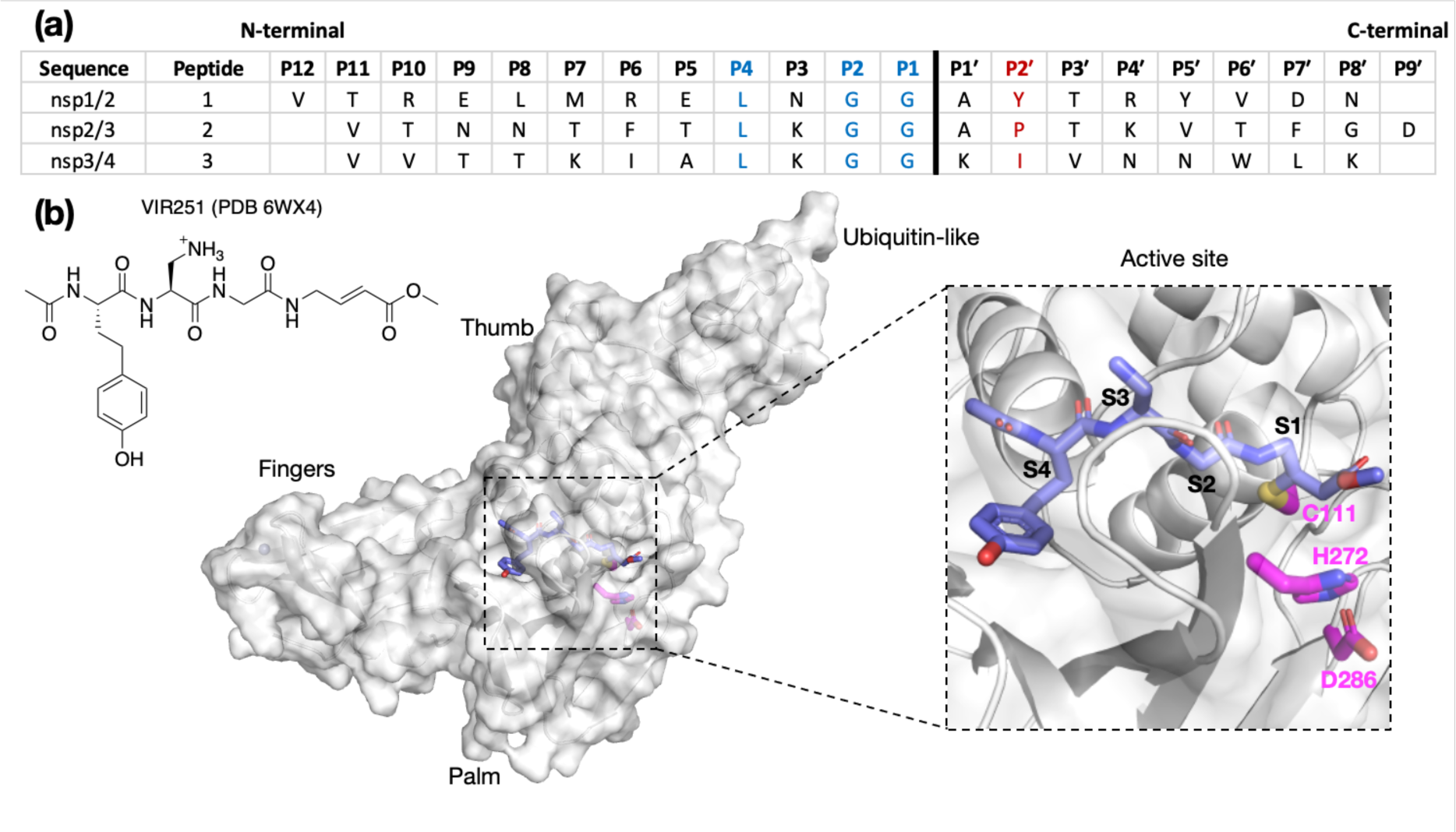
**(a)** Sequences of the reported oligopeptides used in SARS-CoV-2 PL^pro^ mass spectrometry-based assays.^22^ The consensus LXGG sequence residues are in blue; the P2′ residues are in red. **(b)** View from a reported PL^pro^ structure (PDB 6WX4)^13^ in complex with the covalently bound inhibitor VIR251 (carbon atoms in violet), which binds to the S1-S4 subsites; the catalytic triad residues are shown (carbon atoms in magenta). Non-carbon atoms are colored: N in blue; O in red; S in yellow; Zn in gray. Hydrogens are omitted for clarity.

The apparent lack of selectivity for residues binding to the S′ subsites of PL^pro^ may contribute to its reported promiscuity. PL^pro^ catalyzes the hydrolysis of peptide bonds C-terminal to LXGG motifs in various host proteins,^15, 16^ including human interferon regulatory factor 3 (IRF3),^1^ protein S (PROS1),^16^ and the serine/threonine unc-51-like kinase (ULK1).^17^ PL^pro^ also hydrolyzes isopeptide bonds C-terminal to LXGG motifs in proteins that have been post-translationally modified by conjugation of ubiquitin or ubiquitin-like modifiers.^1, 18–21^

Assays using isolated recombinant PL^pro^ have been developed for activity and inhibition studies. Most of these are fluorescence-based and monitor the release of a C-terminal linked fluorescent group.^11, 20, 21, 23–28^ We have previously reported mass spectrometry (MS)-based assays for PL^pro^ and M^pro^ utilizing solid phase extraction coupled to MS (SPE-MS).^22, 29^ The PL^pro^ SPE-MS assay employs a 20-mer peptide, containing the nsp2/3 cleavage site (V808-G818/A819-D827; **Figure 1a**) as a substrate. Surprisingly, little to no hydrolysis was observed for peptides corresponding to the nsp1/2 (V169-G180/A181-N188) and nsp3/4 (V2753-G2763/K2764-K2771) cleavage sites. Similarly, cell-based assays employing oligopeptide linkers with P5-P3′ cleavage site sequences, and liquid chromatography (LC)-based assays employing shorter polyprotein fragments, indicate relatively slow PL^pro^ catalyzed hydrolysis at the nsp1/2 and nsp3/4 sites.^30, 31^ Preferential hydrolysis of the nsp2/3 site has also been observed for SARS-CoV PL^pro^ using three analogous SARS-CoV polyprotein-derived peptide substrates as measured by LCMS assays.^32^ Collectively, these results indicate that the presence of the consensus LXGG P4-P1 residue motif is an insufficient criterion for, at least efficient, PL^pro^ catalysis. Comparison of the P4 to P1′ residues of the three SARS-CoV-2 substrates shows that P3 is the first non-conserved substrate position N-terminal to the scissile amide bond (**Figure 1a**). nsp1/2 has an Asn at P3. While both nsp2/3 and nsp3/4 have a Lys residue at P3, only the former exhibited considerable activity. C-Terminal to the scissile amide, both nsp1/2 and 2/3 have an Ala at P1′, but hydrolysis of nsp1/2 was not observed by SPE-MS. Therefore, the observed difference in activities is likely a result of differences in binding affinity rather than catalytic mechanism.

The binding modes of oligopeptide substrates or peptidomimetic inhibitors in the S4-S1 subsites of PL^pro^ have been characterized crystallographically (*e.g.*, **Figure 1b**).^12, 13, 18, 23, 33^ However, unlike M^pro^, where structures with polyprotein-derived oligopeptides provide insights into S and S′ subsite binding,^34, 35^ there is limited information on binding to the S′ subsites in PL^pro^. Such information may help understanding of the selectivity of PL^pro^ catalysis and aid in inhibitor design. We thus constructed and assessed computational models of SARS-CoV-2 PL^pro^ complexed with its nsp1/2, nsp2/3, and nsp3/4 oligopeptide substrates. Our models presented here suggest that the P2′ proline in the nsp2/3 sequence plays a key role in efficient PL^pro^ catalysis, a proposal validated by residue substitutions and SPE-MS analyses (*vide infra*). The difference in hydrolytic activities between the tested oligopeptides and full-length (poly)proteins suggests that the overall polypeptide conformation is important for substrate recognition. This finding is relevant for future mechanistic and inhibition studies.

## Results and Discussion

### Predicted conformations of the PL^pro^-bound nsp1-3 peptides

To investigate how the SARS-CoV-2 polyprotein nsp1/2, 2/3, and 3/4 cleavage sites bind to PL^pro^, AutoDock CrankPep (ADCP)^36^ was used to dock the peptides previously used in the SPE-MS assay,^22^ *i.e.* **1**, **2**, and **3**, with PL^pro^, using a crystal structure of PL^pro^ covalently linked with the peptidomimetic inhibitor VIR251 (PDB 6WX4, 1.66 Å resolution; see **Methods** for details).^13^ The top 100 ranked peptide poses were analyzed and denoted by “dX_YY”, where “X”=1-3 represents the peptide number according to **Figure 1a** and “YY” indicates the ranking, with dX_01 corresponding to the pose predicted to have the highest binding affinity. In all the highest ranked solutions (d1_01, d2_01, and d3_01), the S2-S1 channel was observed to be occupied (**Supplementary Figure S1**), suggesting it may be a binding hotspot. However, only with d2_01 was the S2-S1 channel observed to be occupied by the P2-P1 Gly-Gly residues in an apparently catalytically relevant manner. Thus, consideration of only the highest ranked solutions was considered to be insufficient for generating catalytically relevant protein-substrate complexes.

Further analysis of the top 100 poses involved filtering based on the position of the P4-P1 residues relative to the binding mode of the inhibitor, VIR251.^13^ A pose was deemed to pass the filter if all four Cα atoms of P4-P1 were within 2 Å of the corresponding VIR251 Cα atoms (**Table S1**). For peptide **1**, only one docked solution passed the filter (d1_16). For peptide **2**, 10 solutions fulfilled the criteria, including the top ranked pose (d2_01, 02, 06, 09, 38, 62, 66, 69, 75, 97); these solutions had a similar conformation to that of d2_01, except for d2_09, which adopts an extended conformation (**Figure S2**). Thus, both d2_01 and d2_09 were selected as representative modes for further modeling. Finally, none of the 100 poses passed the filter for peptide **3**. However, closer inspection of these results revealed multiple solutions in which the P2-P1 Gly-Gly residues were placed in the S2-S1 channel, but in the reverse direction, with P2-Gly in S1 and the P1-Gly in S2. In these conformations, the P1′-Lys was in S3 and the P2′-Ile in S4 (*e.g.*, d3_07; see **Figures 2 and S3**). Although no docked solutions passed the filter for **3**, it does not rule out the possibility of **3** forming a pre-reaction complex with PL^pro^. A model of such a pre-reaction complex was later constructed based on one of the poses that passed the filter for peptides **1** and **2** (d1_16, d2_01, d2_09) (**Figure 2**).

**Figure 2:**
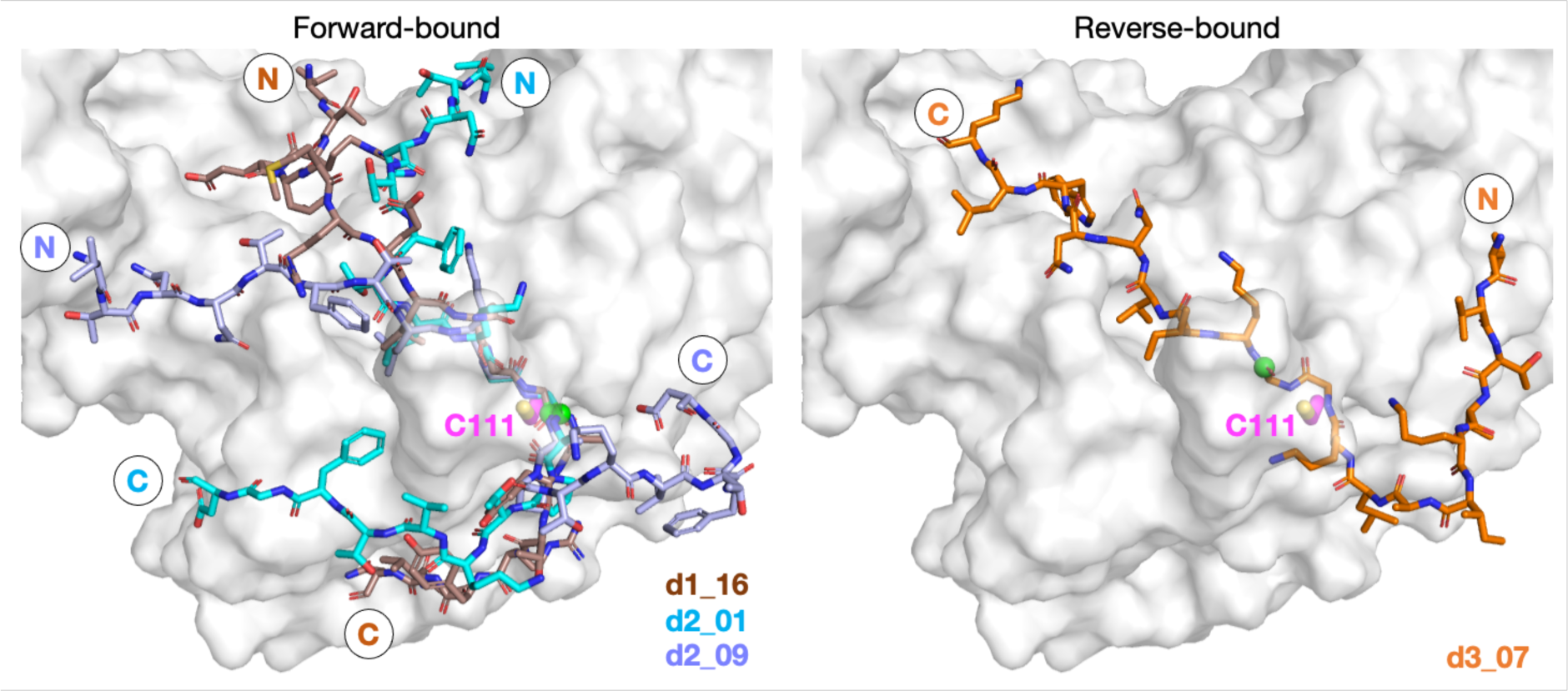
Predicted bound conformations of linear oligopeptides in complex with PL^pro^. AutoDock CrankPep (ADCP)^36^-predicted peptide conformations when non-covalently bound with PL^pro^ (white surface), with PL^pro^ Cys111 residue (magenta sticks) and the peptide N- and C-terminals labeled. The P1 scissile amide carbonyl carbon is shown as a green sphere. The poses are denoted by “dX_YY”, where “X” is the peptide number **1**, **2**, or **3** for the nsp1/2, 2/3, or 3/4 cleavage sites respectively, and “YY” the ranking out of 100 docked poses.

To assess the relative stability of the selected representative peptide poses for **1** and **2**, namely d1_16, d2_01, and d2_09, 3 × 100 ns molecular dynamics (MD) simulations were performed for each system using the GROMACS package with the AMBER99SB-ILDN forcefield and explicit (TIP3P) water solvation,^37–39^ as reported in our previous study on M^pro^-peptide complexes (see **Methods** for details).^40^ Since protonation states are fixed in conventional MD, both the neutral (“N”) and zwitterionic (“Z”) states of the active site Cys111-His272 pair were simulated.

Simulating peptides **1** and **2** in either N- or Z-states of the Cys111-His272 dyad revealed significant deviation in the peptide backbone root mean square deviation (RMSD) and root mean square fluctuation (RMSF) at the N- and C-terminal regions. Notably, the core P5-P1′ region around the consensus LXGG sequence exhibited a more rigid structure (**Figures S4-S5**). Analysis of non-hydrogen atoms RMSF (**Figure S6**) indicates stable binding of the P4-Leu residue sidechain compared to its neighbor residues, reflecting its conserved nature in PL^pro^ substrates (**Figure 1**). Among the three poses studied, d2_01 was the most stable (**Figures 2 and S5**), with relative rigidity of the N-terminal region in the Z-state and of the C-terminal region in the N-state. The d2_01 pose prior to MD was used for comparative modeling of **1** and **3** to explore if all three peptides could bind similarly. However, these MD simulations did not inform on whether the N- or Z-state was preferred in PL^pro^-substrate complexes.

### QM/MM-US calculations indicate a preference for the zwitterionic Cys-His pair

To further explore the preferred ionic state of the Cys111-His272-Asp286 catalytic triad, quantum mechanics/molecular mechanics (QM/MM) umbrella sampling (US) calculations were performed using the Amber18 and Gaussian16 packages (see **Methods**).^41, 42^ Given that the thermodynamic preference may be affected by the presence of substrate, as reported for other cysteine proteases,^43, 44^ both apo PL^pro^ and PL^pro^ non-covalently complexed with **2** were considered (**Figures S7-S11**). In both cases, the results reveal a preference for the Z-state of Cys111-His272 by >5.0 kcal/mol (**Figure S10**). Subsequent proton transfer from His272 to Asp286 leads to a state that is >1.8 kcal/mol higher in energy than when His272 is positively charged and Asp286 is deprotonated (**Figure S11**). Hence, the most thermodynamically favorable state for PL^pro^ in both the apo and substrate-bound cases, at least with peptide **2** in the d2_01 conformation, involves a negatively charged Cys111, a positively charged His272, and a negatively charged Asp286. The thermally accessible activation barriers of the proton transfer steps, which range between 2.6 and 11.9 kcal/mol relative to the Z-state, suggest fast interconversion between N- and Z-states under solution conditions (**Figure S10-S11**). These findings align with reported QM/MM calculations for the papain-like protease cathepsin B^45^ and from constant pH MD simulations of apo coronavirus PL^pro^ enzymes.^46^ However, they differ from a QM/MM study of SARS-CoV-2 PL^pro^ complexed with VIR251,^47^ which suggests that the N-state is thermodynamically favored over the Z-state. Differences in the substrate and the inclusion of Asp286 in our QM region may account for this apparent discrepancy. The preference of the catalytic triad for different ionic states likely depends on the surrounding environment, which is affected by the conformation of the bound peptide and the interactions it establishes with the protein residues, especially those involving nearby peptide P′ residues immediately C-terminal to the scissile amide. While subsequent discussions primarily focus on simulations conducted in the more favored Z-state, the N-state is discussed where necessary, particularly in relation to the P′ interactions.

### Hydrogen bond and dispersion interactions are present both in the core P5-P1′ region and around the peptide N- and C-termini

To investigate if peptides **1** and **3** can bind stably in the d2_01 conformation of **2**, models were constructed (**Figure S12**) considering both the Z- and N-states. For each setup, 3 × 200 ns MD simulations were performed (**Figures S13-S15**). Over the course of MD simulations, the P5-P1′ residues remain stable as indicated by RMSF trends (**Figures S14-S15**). Binding is conferred by a series of hydrogen bond (HB) interactions (**Figure 3a**; **Table S2**). However, these interactions are more poorly maintained around P5-P3 for **3** compared to **1** and **2**. Towards the N-terminus, peptide **3** shows higher flexibility than **1** and **2**, where residues are held more rigidly by interactions with PL^pro^ residues located between the thumb and fingers domains (**Figure S16**).

**Figure 3:**
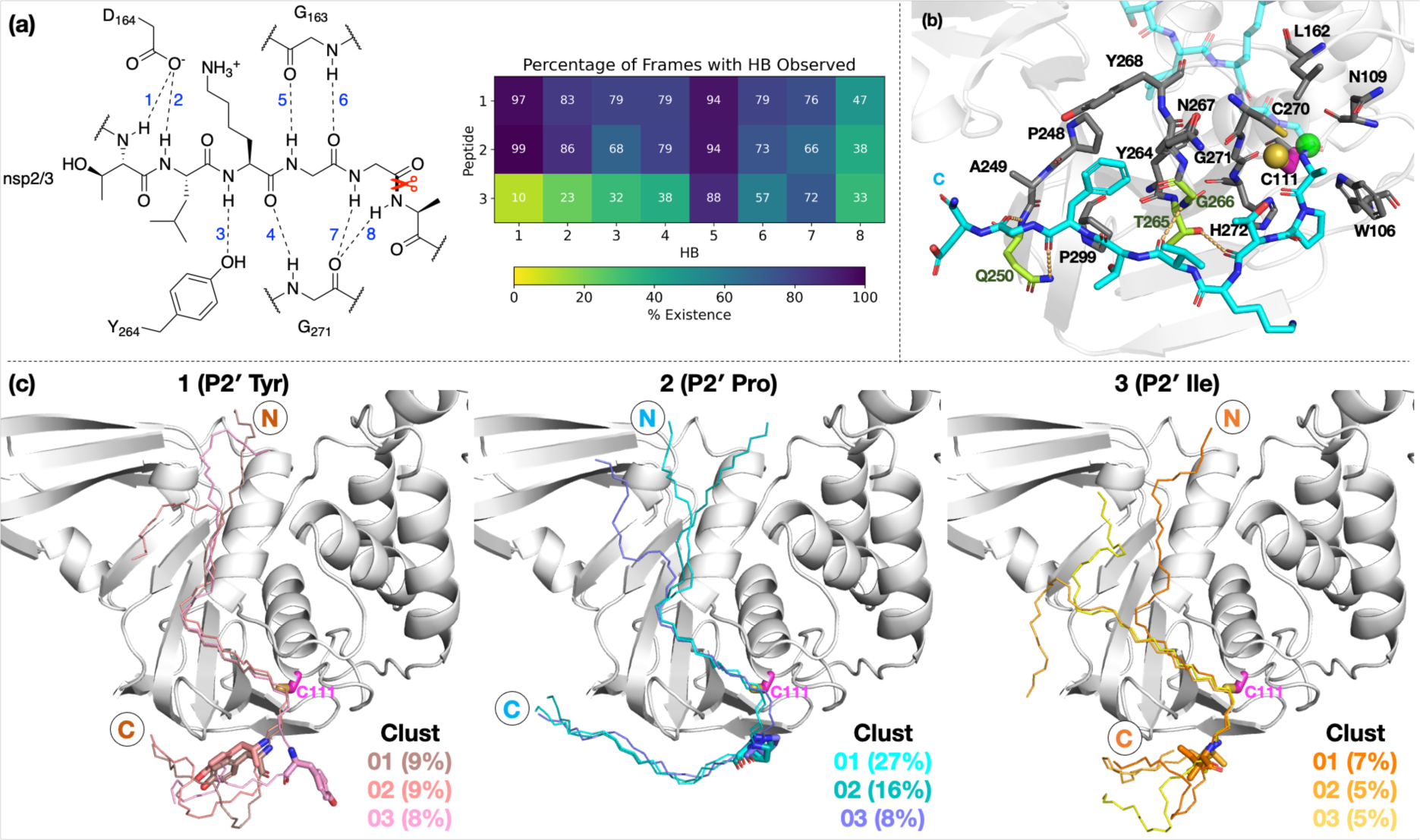
Interactions of PL^pro^ with the oligopeptides 1, 2, and 3. (a) Conserved HBs (occurrence ≥ 25% for at least two peptides) in the P5-P1′ region, observed in 3 × 200 ns MD. (b) Representative structure of N-state PL^pro^ complexed with **2** (cyan; P1 scissile amide carbonyl carbon as a green sphere) from cluster analysis (peptide backbone RMSD cut-off of 3 Å) highlighting interactions in the P′ region. PL^pro^ residues involved in HBs are shown in lime. Residues within 4 Å of P′ residues are in grey, except for Cys111 which is in magenta (its sulfur as a yellow sphere). (c) View of representative structures of the three most populated clusters, from the combined 3 × 200 ns MD of each of the three PL^pro^-peptide (N state) complexes. The peptide backbone is shown as thin sticks and the P2′ residue as thick sticks.

Towards the C-terminus of all three peptides, the P′ residues appear to be more flexible in the Z-state compared to the N-state. The backbone RMSF increases with distance from the active site and reaches values >5 Å after P3′-P5′, except for **2** in the N-state where the RMSF is >5 Å only at P8′ (**Figure S14**). Clustering analysis was performed in the N-state to visualize interactions which might confer stability, using a 3 Å RMSD cut-off on the peptide backbone (**Figure 3b**). For **2** in the N-state, the three most populated clusters (27%, 16%, 8%) have the P′ residues adopting an extended conformation adjacent to the PL^pro^ palm domain (**Figure 3c**). These clusters reveal the presence of four significant HBs, including HBs with Gln250, Thr265, and Gly266, and dispersion interactions with Trp106, Asn109, Cys111, Leu162, Pro248-Gln250, Tyr264-Tyr268, Cys270-His272, and Pro299 (**Figure 3b-c; Table S2**). Notably, the P2′ proline residue in **2** emerges as a crucial residue for inducing and stabilizing a bend centered at the P1′ residue adjacent to the P2′ proline (**Figure S18**). The P2′ proline thus influences the overall peptide conformation and facilitates binding of residues in the P′ region (**Figures 3c and S19-S20**). In **1** and **3**, such the bend centered at P1′ is not as well preserved, and residues C-terminal of P2′ explore wider conformational space in terms of their backbone *ϕ* and *ψ* dihedrals compared to **2** (**Figures S18-S20**).

### A reverse non-reactive binding pose is possible

Our initial docking studies suggested a possible reverse binding mode for **3**, wherein the P2-Gly is positioned in the PL^pro^ S1 subsite and the P1-Gly is positioned in the S2 subsite (**Figures 2 and S3**). To evaluate this binding pose for **3** and investigate if such a reverse binding conformation is possible for **1** and **2**, the d3_07 pose was subjected to initial MD simulations (**Figures S21-S22**). A stable peptide conformation was obtained from these simulations (**Figures S23-S26**) and was used to analogously generate models with **1** and **2** (**Figure S27**). Each system was subjected to 3 × 200 ns MD, with Cys111-His272 in the Z-state.

All of **1-3** bind stably in the reverse mode in the core region of PL^pro^ as indicated by peptide RMSF values (**Figures S28-S30**). Similar to the forward-bound peptides (**Figure 3a**), several HBs are observed between the peptide and PL^pro^ residues, notably, Gly271 and Gly163 in the S3-S1 subsites (**Figure 4**; **Table S3**). However, considering the distance between His272 and the leaving group scissile amide derived nitrogen, it is unlikely that the amide bond at S1 is in a catalytically productive position (**Figure S31**). In the hydrophobic S4 pocket, the original P2′ residues (Tyr, Pro, Ile) of **1**, **2**, and **3**, respectively, are stable according to RMSF analysis (**Figure S30**). Outside of the core S4-S1 region, the peptides manifest high flexibility, except for **3** where the N-terminal residues are stably bound in the PL^pro^ S′ region. Closer inspection reveals that the N-terminal residues of **3** are positioned to form well-maintained HBs and dispersion interactions with PL^pro^ adjacent to the PL^pro^ palm domain (**Figure S32; Table S3**). The simulations suggest the possibility of a reverse, non-productive binding mode for the nsp peptides **1-3.** Note, however, that the biological relevance of this observation for protein substrates is unclear and **3** does not inhibit the PL^pro^-catalyzed hydrolysis of **2**.^22^ Previous studies have shown that non-productive binding of peptides, including D-enantiomers of substrates, with protease substrate binding clefts are possible.^48^ This is perhaps unsurprising given the nature of the interactions between proteases and their substrates, which often involve multiple hydrogen bonds forming β-sheet type structures. In an *in vivo* context, most proteases are likely to encounter several potential substrates, including themselves; hence further work on understanding how selectivity is achieved via non-productive binding, as we observe with PL^pro^, is of interest.

**Figure 4:**
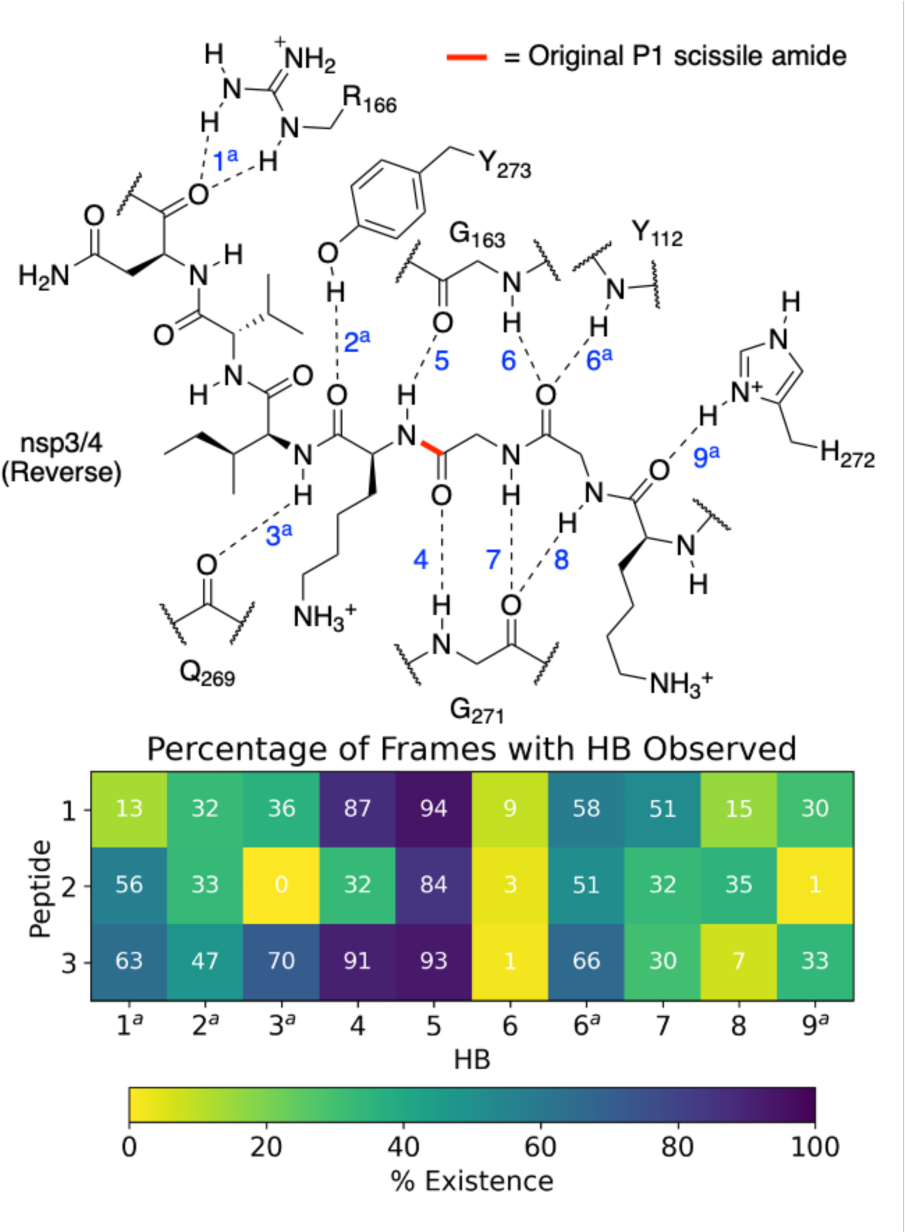
Possibility of reverse binding conformations for the oligopeptides 1, 2, and 3. Occurrence of conserved HBs during 3 × 200 ns MD of PL^pro^ complexed with the three nsp oligopeptides **1**, **2**, and **3** in the reverse binding conformations. The HBs are numbered according to the equivalent peptide position in Figure 3a, with additional HBs carrying the “a” superscript.

### Dynamics of the loop around Tyr268 distinguishes apo and substrate-bound PL^pro^

The dynamics of the PL^pro^ loop around Tyr268 (Gly266-Gly271), also known as the blocking loop 2 (BL2) and which is reported to regulate active site access, ^12, 21, 23, 46, 49, 50^ was analyzed for the apo and PL^pro^ complexes with forward- and reverse-bound peptides in each set of combined 3 × 200 ns MD. This analysis considers the observed proximity and interactions of BL2 residues with forward-bound peptide P′ residues (**Figure 3b**) and reverse-bound peptide P residues (**Figure S32**). The opening and closing states of BL2 were quantified using the backbone RMSF values of BL2 residues and the Pro248–Tyr268 Cα–Cα distance (**Figure 5a**).

**Figure 5:**
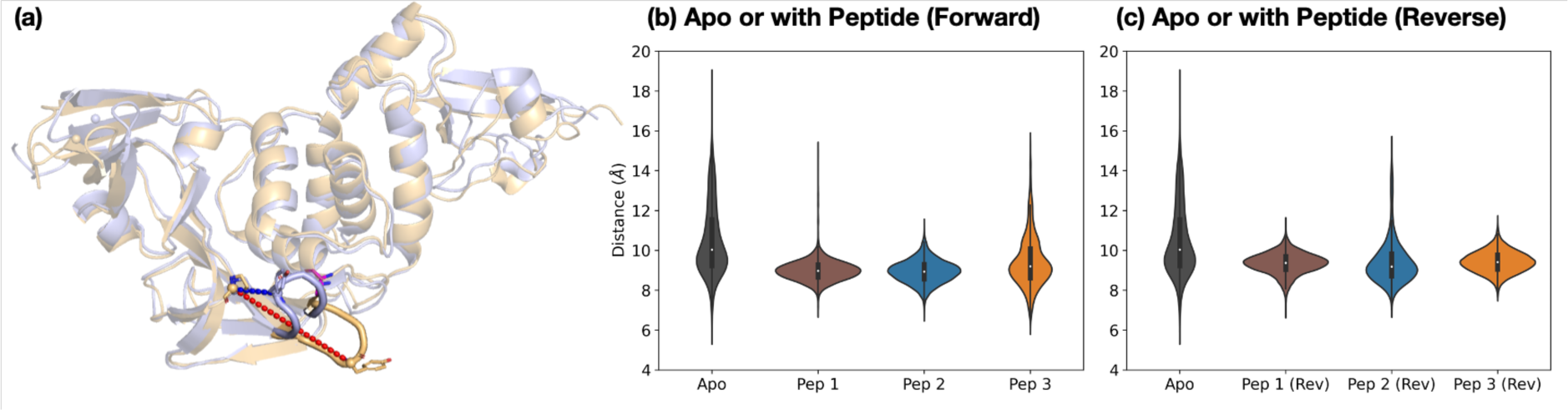
Dynamics of the Tyr268-containing loop PL^pro^ BL2. (a) Views of MD-observed structures of apo PL^pro^ when BL2 is closed (light blue) and open (light orange), which can be quantified by the Pro248–Tyr268 Cα–Cα distance (blue and red dotted lines respectively). The distribution of this distance over combined 3 × 200 ns MD is plotted for PL^pro^ in the apo state and (b) when complexed with the nsp peptides in the forward binding mode, and (c) when complexed with the nsp peptides in the reverse binding mode.

Backbone RMSF analysis indicates that BL2 is more flexible in the apo state compared to the peptide-bound state (**Figure S33-S34**). In the apo PL^pro^, the RMSF of the most flexible residue in BL2, Tyr268, is 2.5 ± 0.1 Å, whereas in the substrate peptide **2**-bound PL^pro^ it is 0.98 ± 0.03 Å (**Figure S35**). For the apo system, multiple BL2 opening and closing events were observed to occur, an observation which may be related to substrate capture processes (**Figures 5 and S36**).

In the forward-bound peptide-PL^pro^ complexes, BL2 becomes more rigid compared to the apo-form, although BL2 opening still occurred when bound to peptide **3**, and to a lesser extent when bound to peptide **1** (**Figures 5b and S36**). Such opening events with **1** or **3** bound may reflect the tendency of PL^pro^ to revert to its apo form, indicating binding instability and a propensity towards peptide dissociation. By contrast, complexation with peptide **2** results in a rigidly closed BL2 throughout the simulations, possibly as a result of the stronger interactions between the P′ residues of **2** and PL^pro^ residues close to BL2 (**Figure 3b**).

The reverse-bound complexes show the opposite trend to the forward-bound complexes (**Figures 5c and S34-S36**). BL2 remains rigidly closed over the course of MD with peptides **1** and **3**, but opens multiple times with **2**. Previous studies have reported that ligand occupation at the S4 and S3 subsites induces BL2 closure.^20, 23, 24, 49^ However, here both **1** and **2** have a P1′ Ala residue which binds in S3, and yet BL2 is more flexible when bound with **2** than with **1**. The difference in BL2 flexibility probably arises from the different residues binding in the S4 pocket. In **1** and **3**, the S4 pocket is occupied by the hydrophobic sidechains of P2′ Tyr and Ile respectively, whereas it remains mostly vacant in **2** where the P2′ Pro lacks an extended hydrophobic sidechain (**Figure S37**).

The combined evidence from our modeling studies suggests that the more efficient reaction of PL^pro^ with peptide **2** compared to peptides **1** or **3** can be attributed to the presence of a Pro residue at the P2′ position. This results in a bound pre-reaction complex conformation where the P′ residues favorably interact with PL^pro^ and promote the closure of BL2, thus reducing the probability of peptide dissociation. Additionally, compared to **1** or **3**, peptide **2** shows a greater preference for the productive forward binding mode relative to the competing non-productive reverse binding mode.

### A proline residue at the substrate P2′ position enhances PL^pro^-catalyzed hydrolysis of peptides based on the viral polyprotein

We next investigated the proposed role of the P2′ proline residue of the nsp2/3 (**2**) oligopeptide for SARS-CoV-2 PL^pro^ catalysis. Using solid phase peptide synthesis (SPPS), we synthesized nsp1/2 (**1**)- and nsp3/4 (**3**)-based peptides where the wildtype (wt) P2′ residue of the Wuhan-Hu-1 strain of SARS-CoV-2^51^ was substituted for a proline, as in the wt nsp2/3 peptide sequence (**2**), *i.e.* the nsp1/2(Y182P) (**4**) and nsp3/4(I2765P) (**5**) variant peptides (**Figure 6a**).

**Figure 6.**
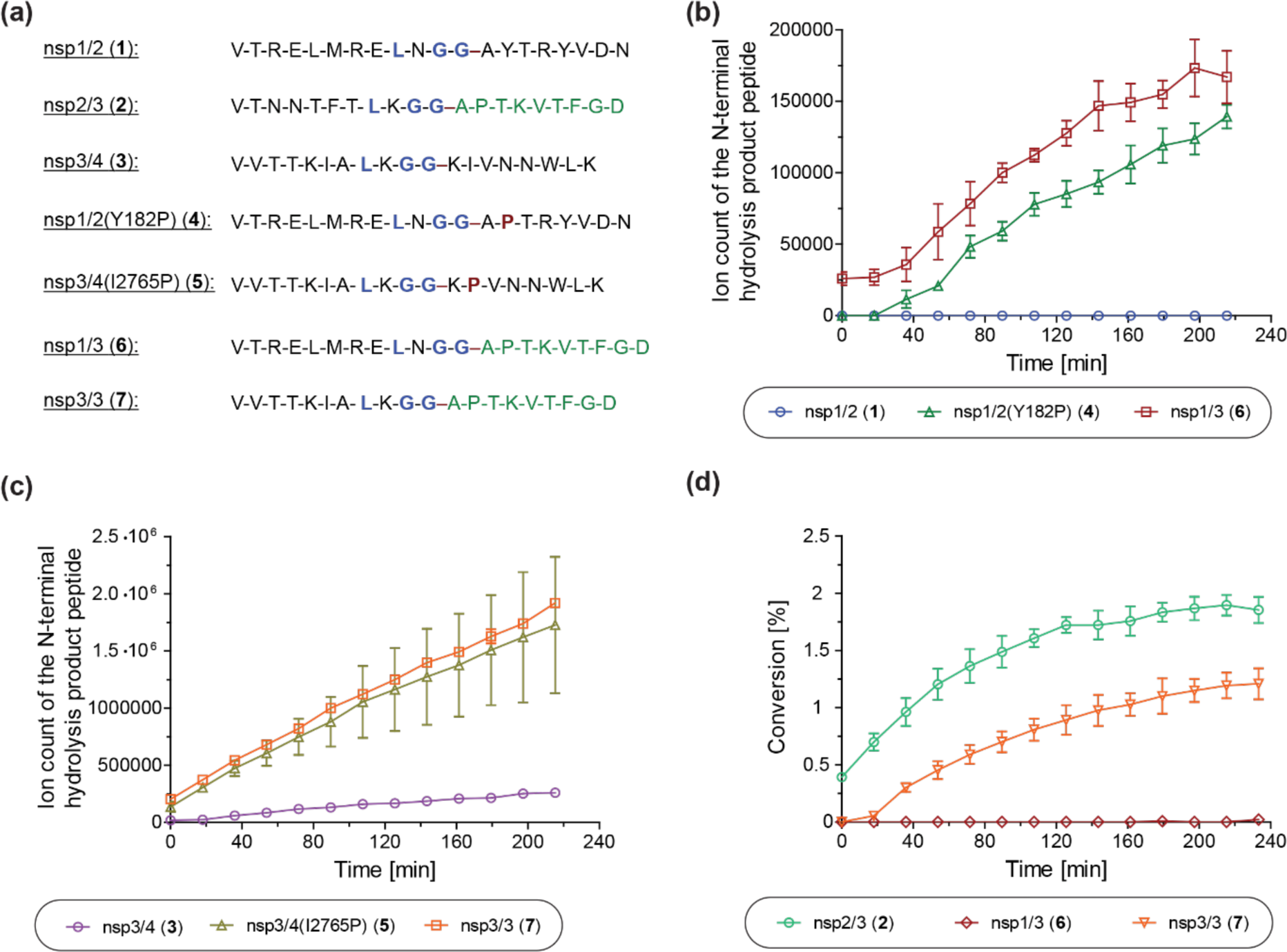
A proline at the substrate P2′ position enhances PL^pro^ catalysis. (**a**) Sequences of the employed SARS-CoV-2 polyprotein-derived oligopeptides used; (**b**) ion counts of the N-terminal hydrolysis product peptides of wt nsp1/2 (**1**; blue circles), nsp1/2(Y182P) (**4**; green triangles), and nsp1/3 (**6**; red boxes) incubated with SARS-CoV-2 PL^pro^; (**c**) ion counts of the N-terminal hydrolysis product peptides of wt nsp3/4 (**3**; lavender circles), nsp3/4(I2765P) (**5**; ochre triangles), and nsp3/3 (**7**; orange boxes) incubated with SARS-CoV-2 PL^pro^; (**d**) conversion of the PL^pro^-catalyzed hydrolysis of wt nsp2/3 (**2**; light green circles), nsp1/3 (**6**; red diamonds), and nsp3/3 (**7**; orange inverse triangles), determined using the reported N-terminal acetylated C-terminal hydrolysis product peptide of **2**^22^ as an internal standard (0.2 µM). Reactions were performed using peptide (2.0 µM) and SARS-CoV-2 PL^pro^ (0.2 µM) in buffer (50 mM Tris, pH 8.0, 20 °C) and analyzed using SPE-MS under reported conditions.^22^ Results are means of three independent runs (n = 3; mean ± standard deviation, SD).

The nsp1/2(Y182P) (**4**) and nsp3/4(I2765P) (**5**) variant peptides were incubated with isolated recombinant SARS-CoV-2 PL^pro^ under reported conditions (50 mM Tris, pH 8.0, 20 °C)^22^ and peptide hydrolysis was monitored by SPE-MS. The results reveal that PL^pro^-catalyzed hydrolysis of **4** and **5** is substantially more efficient compared to wt nsp1/2 and nsp3/4 peptides **1** and **3**, respectively, which were used as controls. PL^pro^ catalyzed the hydrolysis of **4**, albeit at low levels, whereas it did not catalyze the hydrolysis of the wt nsp1/2 peptide (**1**) under the tested conditions (**Figure 6b**). The PL^pro^-catalyzed hydrolysis of **5** was ∼10-fold more efficient than that of wt **3** (**Figure 6c**). Note that the ion counts of the N-terminal product peptides were used for comparison of the hydrolysis efficiencies of the wt and variant peptides.

To investigate whether residues C-terminal to the LXGG motif of the nsp2/3 peptide, other than the P2′ proline residue, contribute to enhanced PL^pro^ catalysis, chimeric oligopeptides of nsp1/2 (**1**) and nsp3/4 (**3**) containing the entire fragment C-terminal to the LXGG motif of the nsp2/3 peptide (**2**) were synthesized, *i.e.* the nsp1/3 (V169-G180/A819-D827) (**6**) and nsp3/3 (V2753-G2763/A819-D827) (**7**) hybrid peptides (**Figure 6a**). These chimeric peptides were incubated with isolated recombinant SARS-CoV-2 PL^pro^ and hydrolysis was monitored using SPE-MS.^22^ The results revealed that the efficiency of the PL^pro^-catalyzed hydrolysis of the hybrid peptides **6** and **7** was similar, within experimental error, to those of the nsp1/2(Y182P) and nsp3/4(I2765P) variant peptides **4** and **5**, respectively, indicating that residues at the P′ positions other than the P2′ proline did not have pronounced effects on PL^pro^ catalysis, within the context of the tested peptides (**Figures 6b and 6c**).

Reactions with the nsp1/3 and nsp3/3 hybrid peptides **6** and **7** were performed in the presence of the reported N-terminal acetylated C-terminal hydrolysis product peptide of the wt nsp2/3 peptide (**2**),^22^ which was used as an internal standard to quantify peptide hydrolysis. By comparing the PL^pro^-catalyzed hydrolysis of the wt nsp2/3 peptide (**2**) with peptides **6** and **7**, it became evident that **2** was a substantially more efficient substrate of PL^pro^ than both **6** and **7**, as well as, by implication, than the nsp1/2(Y182P) and nsp3/4(I2765P) variant peptides **4** and **5**. The PL^pro^-catalyzed hydrolysis of **2** was ∼2-fold more efficient than that of **7**, while the hydrolysis levels of **6** were marginal (**Figure 6d**). This observation suggests that substrate residues which bind to the S subsites are, in general, more important for efficient PL^pro^ catalysis than those binding to the S′ subsites, in accord with previous studies on SARS-CoV and SARS-CoV-2 PL^pro^, which have shown a preference for lysine over asparagine residues at the P3 position,^13, 31, 32, 52^ as well as with other cysteine proteases, such as SARS-CoV-2 M^pro^.^40^ Note that the low absolute levels of hydrolysis of the wt nsp2/3 peptide (**2**) may be a result of performing SPE-MS assays at 20 °C rather than at 37 °C and, potentially, of using relatively short oligopeptides instead of fully folded polyproteins as substrates, which are likely unable to bind to PL^pro^ at allosteric positions; substrate binding to pockets distal to the PL^pro^ active site has been reported to be essential for efficient catalysis.^53^

### The primary sequence of LXGG-motif containing human proteins affects their ability to be substrates of SARS-CoV-2 PL^pro^

Human proteins containing an LXGG motif in their sequence have been identified as (potential) substrates of SARS-CoV-2 PL^pro^. This conclusion is based on *in vitro* and cell-based studies,^15^ including with interferon regulatory factor 3 (IRF3),^1^ protein S (PROS1),^16^ and serine/threonine-protein kinase ULK1.^17^ To investigate these potential substrates further, we synthesized SPE-MS compatible peptides based on the sequences of IRF3 (**8**), PROS1 (**9**), ULK1 (**10**), and the ubiquitin-like modifier-activating enzyme ATG7 (ATG7, **11**). ATG7 was also investigated as a potential human PL^pro^ substrate, because, like ULK1, it is involved in autophagy induction and bears a LGGG motif; in addition, it has a Pro residue at the P2′ corresponding position.^54, 55^

The peptides were incubated with isolated recombinant SARS-CoV-2 PL^pro^ under standard conditions (50 mM Tris, pH 8.0), though at 37 °C, because of the low levels of PL^pro^ catalyzed hydrolysis initially observed at 20 °C. Analysis of the peptide mixtures after 12 h of incubation with PL^pro^ revealed that PL^pro^ catalyzes hydrolysis of PROS1 (**9**) and ULK1 (**10**) more efficiently than IRF3 (**8**). However, PL^pro^ did not catalyze the hydrolysis of ATG7 (**11**) under the tested conditions (**Table 1**). PL^pro^-catalyzed hydrolysis of PROS1 (**9**) and ULK1 (**10**) was, however, substantially less efficient than for nsp2/3 (**2**) (**Table 1**, entry ii), though was more efficient than for nsp1/2 (**1**) and nsp3/4 (**3**). This suggests that PL^pro^ may catalyze the hydrolysis of certain human proteins more efficiently than of, at least some, of its natural viral substrates. Note that conversions were estimated by comparing starting material depletion with no-enzyme controls; the identity of the N- and C-terminal product peptides was confirmed by MS.

**Table 1.**
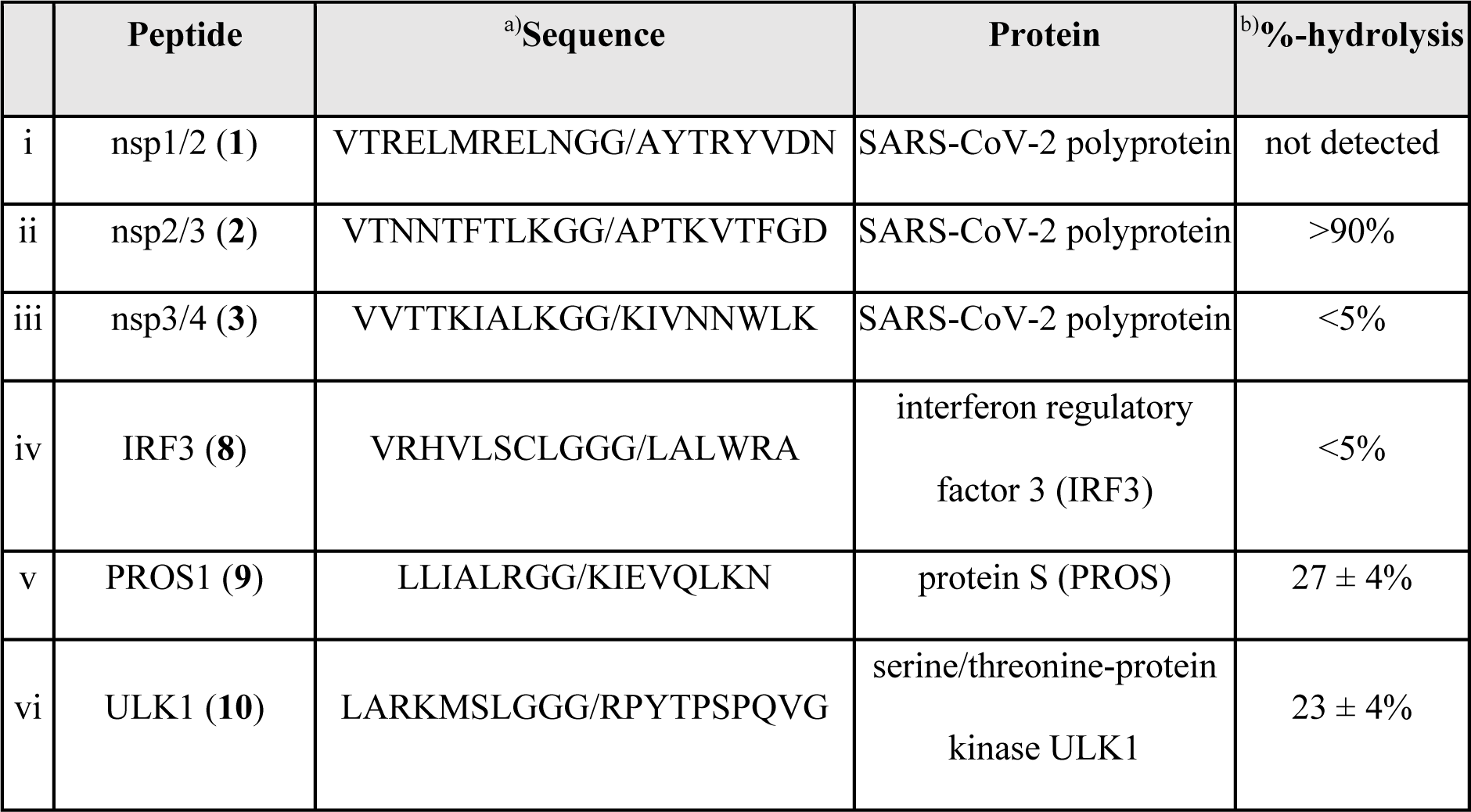
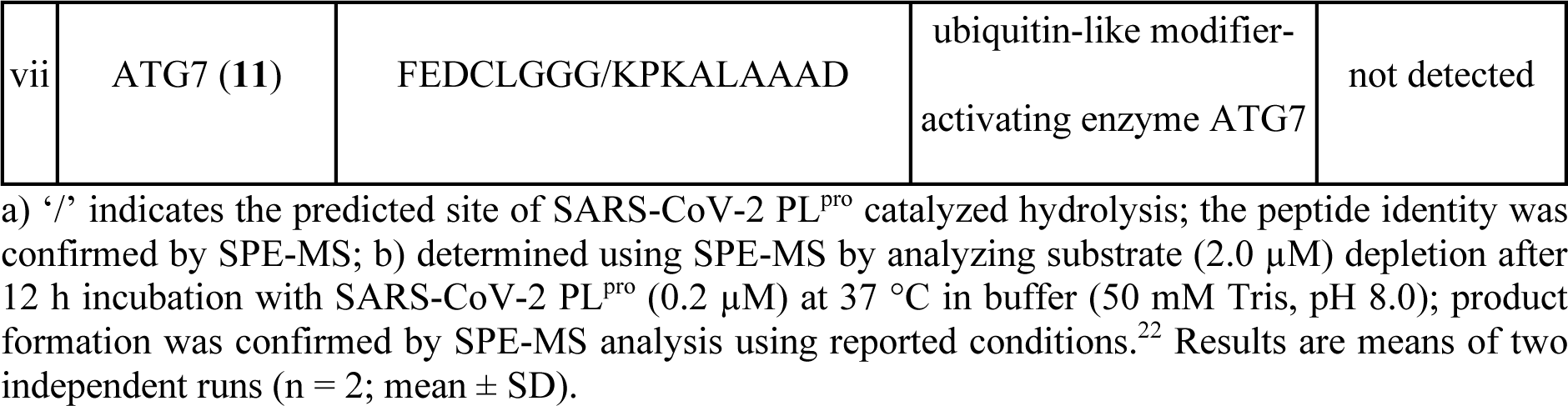
Comparison of the SARS-CoV-2 PL^pro^-catalyzed hydrolysis of LXGG-motif containing peptides derived from the primary sequence of viral and human proteins as estimated by SPE-MS analysis.

Interestingly, the results indicate that a proline residue at the substrate P2′ position does not necessarily result in improved PL^pro^ catalysis. For example, the ULK1-derived peptide **10**, which has a P2′ proline residue, shows ∼23% hydrolysis after 12 h, whereas hydrolysis products were not observed for ATG7 peptide (**11**), which also contains a proline residue at P2′ (**Table 1**, entries vi and vii). By contrast, the PL^pro^-catalyzed hydrolysis of PROS1 (**9**) proceeded with similar efficiency as that of ULK1 (**10**), *i.e.* ∼27% hydrolysis after 12 h, though it does not bear a proline residue at P2′ (**Table 1**, entry v). These combined results suggest that factors other than the presence of an LXGG motif and a proline residue at P2′ determine the efficiency of PL^pro^ catalysis, potentially including the protein/peptide fold. In addition, other, yet unidentified, interactions of primary sequence residues with PL^pro^ which bind near the active site may be of importance. For instance, it appears that efficient substrates bear an alcohol-containing side chain at P5 (Ser/Thr).

### Predicted binding modes of human protein-derived peptides substrates to PL^pro^

To investigate how peptides **8**, **9**, **10**, and **11**, derived respectively from IRF3, PROS1, ULK1, and ATG7, may bind to SARS-CoV-2 PL^pro^, the same docking procedures used for the nsp peptides **1-3** were applied, with assessment of the poses relative to the Cα atoms of the peptidomimetic inhibitor VIR251 in complex with PL^pro^ as observed by crystallography (PDB 6WX4)^13^ (**Tables S4-S5**). Among these peptides, only PROS1 (**9**) returned conformations that passed the filter (**Figure S38**). Despite the lack of a P2′ Pro, the PROS1 P′ residues are predicted to adopt a similar conformation as d2_01 for **2**. In the PROS1 poses, the hydrophobic sidechains of P2′-Ile, P4′-Val, and P6′-Leu point towards BL2, while the hydrophilic P1′-Lys, P3′-Glu, P5′-Gln, and P7′-Lys sidechains point in the opposite direction and/or outwards into solvent. These interactions in the P′ region with PL^pro^ might contribute to the formation of a productive pre-reaction complex as investigated with **2**. For ULK1 (**10**), despite the observation of efficient hydrolysis and the presence of P2′-Pro, none of the 100 predicted conformations passed the filter. Multiple poses nearly passed the filter, placing the P3-P1 residues in the corresponding S3-S1 subsites but not the P4-Leu in S4, and the P′ residues interacting with PL^pro^ underneath BL2 (**Table S4** and **Figure S39**). For IRF3 (**8**), for which efficient hydrolysis was not observed, none of the 100 docked conformations passed the filter. Several of the IRF3 poses were observed to bind in reverse, with the P1′-Leu in S4, P1-Gly in S3, P2-Gly in S2, and P3-Gly in S1, which might compete with productive binding (**Figure S40**). For ATG7 (**11**), hydrolysis of which was not detected by SPE-MS, none of the 100 docked conformations passed the filter. The only poses binding within the S2-S1 channel place the consecutive P6′-P8′ Ala-Ala-Ala in the S2-S4 subsites respectively, with the peptide bound in the reverse direction (**Figure S41**). Overall, these results suggest that whilst crystallography-based docking predictions can be helpful in suggesting possible substrate binding poses, they are unlikely to predict whether a peptide is hydrolyzed efficiently by PL^pro^, which is regulated by a complex mixture of factors, reflecting the relatively high substrate promiscuity of PL^pro^, including its ability to accept both peptide and isopeptide substrates.

### Conclusions

Several studies have provided evidence that PL^pro^ catalysis of nsp2/3 cleavage is more efficient than that of nsp1/2 or nsp3/4, indicating that the presence of an LXGG motif alone is insufficient to predict catalysis efficiency.^22, 30–32, 56^ The interaction of viral proteases with the (human) host proteome is well-documented,^57, 58^ including for SARS-CoV-2 PL^pro^, which is reported to possess deubiquitinase activity and catalyze the removal of (poly)ubiquitin and ubiquitin-like modifiers bearing a LXGG motif from post-translationally modified human proteins.^18, 21^ However, factors that govern the ability of SARS-CoV-2 PL^pro^ to catalyze peptide bond hydrolysis C-terminal to LXGG motifs in human proteins are not well understood.

Our combined modeling and experimental results reveal that the primary sequence of potential substrate proteins apart from LXGG motifs affect the efficiency of PL^pro^-catalyzed substrate hydrolysis. Studies on the bound conformations and the substrate-PL^pro^ dynamics suggest that having a proline at the P2′ position can, at least in some cases, promote catalysis of isolated recombinant PL^pro^. This proposal is validated by residue substitutions and *in vitro* turnover assays. The results suggest that the activity enhancement with a proline at the P2′ position is likely due to interactions between PL^pro^ and the P′ residues, which are enhanced by the P2′ proline induced bend in peptide fold, and, possibly, unfavorable binding of a proline in the hydrophobic S4 pocket which may limit unreactive reverse direction binding modes in the active site.

Our results also suggest that whilst docking predictions can inform potentially productive binding modes, they do not confidently predict whether a potential sequence will be efficiently hydrolyzed by PL^pro^. Studies on the PL^pro^-catalyzed hydrolysis of LXGG motif-containing oligopeptides derived from human proteins*, i.e*., IRF3, PROS1, ULK1, and ATG7, suggest that, in addition to the presence of an LXGG motif and a proline residue at P2′, other factors are important in regulating the efficiency of PL^pro^ catalysis; efficient PL^pro^-catalyzed hydrolysis can occur without a P2′ proline in the substrate.

Our data shows that primary sequences that may *per se* appear unfavorable for PL^pro^ catalysis, such as those lacking a proline at P2′, can still be efficient substrates provided that *e.g.* the protein fold favors binding to PL^pro^, as is likely the case for the viral polyprotein nsp1/2 cleavage site. Conversely, primary sequences that *per se* appear to favor PL^pro^ catalysis might in fact not be efficient PL^pro^ substrates in cells. Thus, care needs to be taken when investigating human proteins for substrate activity with PL^pro^; studies aimed at identifying novel human SARS-CoV-2 PL^pro^ substrates, and by extension substrates of other viral proteases, should involve work with folded proteins, ideally combined with proteomic studies of infected cells to correlate *in vitro* and *in silico* results to real biology.

The importance of non-covalent interactions at the P′ substrate regions in enhancing PL^pro^ proteolytic activity likely extends well beyond the consensus LXGG motif, as demonstrated with other cysteine proteases such as hepacivirus NS2-3 proteases.^59–63^ From a basic enzymology perspective, it is of interest to explore whether the different substrate binding mechanisms used by different proteases reflect their biological roles, including hydrolysis efficiency and selectivity. PL^pro^ and M^pro^ are excellent models to explore this question given their medicinal interest and apparently different substrate selectivities.

## Methods

### Peptide docking

The structure of SARS-CoV-2 PL^pro^ in which the active site Cys111 was covalently attached to the peptidomimetic inhibitor VIR251 was taken from the RCSB Protein Data Bank (PDB ID 6WX4, https://www.rcsb.org/structure/6WX4).13 Protein affinity maps covering the entire protein (center = (-0.785, -19.447, -32.879); box dimensions (Å) = (65.25, 50.25, 99.75)) were calculated with AutoGridFR (v. 1.2).^64^ Starting from an extended conformation and with the search space covering the entire protein, each oligopeptide was docked with AutoDock CrankPep (ADCP; in ADFRsuite v. 1.0),^36^ with 100 replicas and 3M energy evaluations per peptide amino acid, and the solutions outputted following clustering with a native contact cut-off of 0.8. The 100 outputted solutions were then filtered based on a comparison of the docked P4-P1 residues with VIR251. A solution was deemed to pass the filter if all four Cα atoms in P4-P1 were within 2 Å of the corresponding VIR251 Cα atoms. Protein-peptide complexes were visualized using PyMOL (open source, v. 2.3.0).^65^

### System setup

The PL^pro^ structure (PDB 6WX4) was prepared with Reduce (MolProbity, Duke University),^66^ with protonation states determined with H++ (Virginia Tech),^67^ PROPKA3 (PDB2PQR),^68^ and visual inspection. No flips were applied. All Asp/Glu residues were modeled as deprotonated; all Lys/Arg were protonated; all Cys residues were neutral and protonated, except the four Zn-coordinating residues (Cys189, 192, 224, 226), and Cys111 which is discussed below; His assignments are described in **Table S6**. For the active site Cys111 and His272 residues, both the neutral (“N”; His272 neutral, Nɛ-protonated) and zwitterionic (“Z”) states were considered. Using PyMOL (open source, v. 2.3.0),^65^ the protein N-terminal was capped with an acetyl group (ACE-EVRTI) and the C-terminal was capped with N-methyl (TTIKP-NME). The resultant charge of the protein, including Zn^2+^, was +1. Crystallographic waters were retained. In the peptides, the N-terminals were uncapped and the C-terminals NH_2_-capped, in accord with experimental conditions. For the comparative modeling of the nsp peptides, the residues in the original peptide were modified to match the peptide sequences of interest using the mutagenesis tool of PyMOL (open source, v. 2.3.0).^65^ For each residue, the least sterically clashing backbone-dependent rotamer was adopted.^69^

### Molecular dynamics (MD) simulations

MD simulations were conducted with GROMACS (versions 2019.2 and 2020.4),^37^ using the AMBER99SB-ILDN forcefield for amino acid residues,^38^ non-bonded parameters for Zn,^70^ and the TIP3P water model.^39^ The system was placed in a rhombic dodecahedral box (1 nm buffer), solvated, neutralized with sodium/chloride ions, and minimized until maximum force fell below 1000 kJ•mol^-1^ nm^-1^. By generating random velocities at 298.15 K, three replicas per system were initiated, then equilibrated with non-hydrogen atom restraints for 200 ps (1 fs step) under NVT conditions at 298.15 K, equilibrated for 200 ps (1 fs step) under NPT conditions at 298.15 K and 1.0 bar, before a 100 ns or 200 ns (2 fs step) production MD. Temperature was maintained at 298.15 K with a velocity-rescaling thermostat with a stochastic term (time constant 0.1 ps; separate coupling of protein and non-protein),^71^ and pressure at 1.0 bar with a Parrinello-Rahman barostat (time constant 2 ps).^72, 73^ Long-range electrostatics was treated with smooth Particle-mesh Ewald (1 nm cut-off).^74, 75^ Van der Waals interactions were treated with a 1 nm cut off. Analysis of production MD simulations was conducted using GROMACS tools (v 2019.2).^37^ To obtain representative structures of peptide-bound PL^pro^, clustering was performed with a 3 Å RMSD cut-off on the peptide backbone (when the focus is on peptide fold) or peptide non-hydrogen atoms (when the focus is on PL^pro^-peptide interactions, including sidechain interactions) using the gromos algorithm.^76^ A hydrogen bond (HB) was assigned if the donor-acceptor distance was below 3.5 Å and the hydrogen-donor-acceptor angle was below 30°. Peptide secondary structure assignments were conducted using DSSP (version 2.0.4).^77, 78^

### Quantum mechanics/molecular mechanics (QM/MM) calculations

To investigate the preferred state of the catalytic triad, proton transfers between Cys111 and His272, and between His272 and Asp286, were modeled with QM/MM umbrella sampling (US). For each of the apo or peptide **2**-bound PL^pro^ in the N- or Z-state, one representative structure was extracted from each of the three MD repeats (200 ns for apo PL^pro^; initial 100 ns for peptide **2**-bound PL^pro^; frames analyzed every ns) as the starting configuration, selecting the frame that had the lowest RMSD compared to the average PL^pro^(-peptide) structure in the simulation while satisfying all of the following distance criteria: (i) (for N-state) *d*(C111_HG - H272_ND1) or (for Z-state) *d*(C111_SG - H272_HD1) ≤ 2.5 Å; (ii) *d*(D286_ODX - H272_HE2) ≤ 2.5 Å (X = 1 or 2); and in the peptide-bound cases, (iii) *d*(C111_SG - peptide P1_C) ≤ 3.5 Å. A 6 Å solvation shell was retained in the structure. Using Amber18,^41^ each system was re-solvated with a 10 Å buffer region, neutralized, minimized, and equilibrated under NVT (10 ps) and NPT (20 ps) conditions to 298.15 K and 1.0 bar, using the MM forcefield restraining all protein and peptide atoms.

The QM region includes the sidechains of PL^pro^ Cys111, His272 and Asp286, and in the case of peptide **2**-bound PL^pro^, the substrate P2 (atoms C and O), P1 (all atoms), and P1′ (atoms except C and O), with a total of 25 or 44 atoms including link atoms for apo or peptide **2**-bound PL^pro^, respectively (**Figure S7**). The Cys111-His272 proton transfer was sampled (*k* = 100 kcal mol^-1^ Å^-2^) with the reaction coordinate (RC) = *d*(C111_SG - H) - *d*(H272_ND1 - H) between -1 (N-state) and 1 (Z-state) Å in 0.1 Å intervals. In each of the 21 windows, 12.5 ps (1 fs step) of DFTB3/MM-MD was performed under NPT conditions,^79^ with the first 2.5 ps discarded as equilibration. Similarly, the His272-Asp286 proton transfer was sampled with RC = *d*(H272_NE2 - H) - *d*(D286_ODX - H) (ODX being the less solvent-exposed carboxylate oxygen) from -1 (doubly protonated His272 and deprotonated Asp286) to 1 (neutral His272 and Asp286) Å in 0.1 Å intervals. In each window a 500 fs (1 fs step) DFT/MM-MD was performed at the PBE0-D3BJ/6-31G(d) level of theory,^80–83^ using Amber18/Gaussian16 (A.03).^41, 42^ Free energy profiles were obtained by the weighted histogram analysis method (WHAM).^84^

### Protein production

The PL^pro^ domain of the SARS-CoV-2 nsp3 (region E746-T1063) was produced using *E. coli* Lemo21(DE3) cells and purified as reported previously.^22^

### Oligopeptide synthesis

Oligopeptides **4**-**11** were prepared from the C- to N-termini by solid phase peptide synthesis (SPPS) using a Liberty Blue peptide synthesizer (CEM Microwave Technology Ltd.), analogous to the reported synthesis of the oligopeptides **1**-**3**.^22^ Rink Amide MBHA resin (AGTC Bioproducts Ltd.) was employed to obtain the oligopeptides as C-terminal amides following cleavage from the resin and purification by semi-preparative HPLC (Shimadzu UK Ltd.).

### Solid phase extraction mass spectrometry (SPE-MS)

SPE-MS assays were performed using a RapidFire RF 365 high-throughput sampling robot (Agilent) attached to an iFunnel Agilent 6550 accurate mass quadrupole time-of-flight (Q-TOF) mass spectrometer operated in the positive ionization mode equipped with a C4 SPE cartridge, as reported.^22^ Assay conditions: isolated recombinant SARS-CoV-2 PL^pro^ (0.2 µM), oligopeptides **4**-**11** (2.0 µM), and, if appropriate, the N-terminal acetylated C-terminal hydrolysis product peptide of the wt nsp2/3 peptide (**2**) (0.2 µM)^22^ in buffer (50 mM Tris, pH 8.0) at 20 ⁰C or 37 ⁰C.

## Data Availability

Detailed system setup and analysis of docking, MD, and QM/MM-US calculations are available in the Supporting Information. All PL^pro^-peptide models and MD simulation data (including input and trajectory files) are openly available on GitHub at: https://github.com/duartegroup/PLpro_MD.

## Author Contributions

HTHC, LB, CJS and FD conceptualized the study. HTHC carried out computational calculations and analyses. LB carried out oligopeptide synthesis, and SPE-MS experiments and analyses. PL and CS-D carried out PL^pro^ production with supervision from MAW. HTHC and LB wrote the original draft. All authors participated in reviewing of the manuscript. CJS and FD supervised the study.

## Acknowledgements

We acknowledge the philanthropic support of the donors to the University of Oxford’s COVID-19 Research Response Fund and King Abdulaziz University, Saudi Arabia, for funding. This research was funded in part by the Wellcome Trust (106244/Z/14/Z). For the purpose of open access, the authors have applied a CC BY public copyright license to any Author Accepted Manuscript version arising from this submission. We thank Cancer Research UK (C8717/A18245) and the Biotechnology and Biological Sciences Research Council (BB/J003018/1 and BB/R000344/1) for funding. HTHC thanks the Clarendon Fund, New College Oxford, and the EPSRC Centre for Doctoral Training in Synthesis for Biology and Medicine (EP/L015838/1) for a studentship, generously supported by AstraZeneca, Diamond Light Source, Defence Science and Technology Laboratory, Evotec, GlaxoSmithKline, Janssen, Novartis, Pfizer, Syngenta, Takeda, UCB and Vertex. This project made use of computer time on HPC granted via the UK High-End Computing Consortium for Biomolecular Simulation, HECBioSim (http://hecbiosim.ac.uk), supported by EPSRC (grant no. EP/R029407/1).

