## Supplementary material for "Studies on the selectivity of the SARS-CoV-2 papain-like protease reveal the importance of the P2′ proline of the viral polyprotein": SI

[c]Research Complex at Harwell, Harwell Science and Innovation Campus, Didcot, OX11 0FA, United Kingdom.

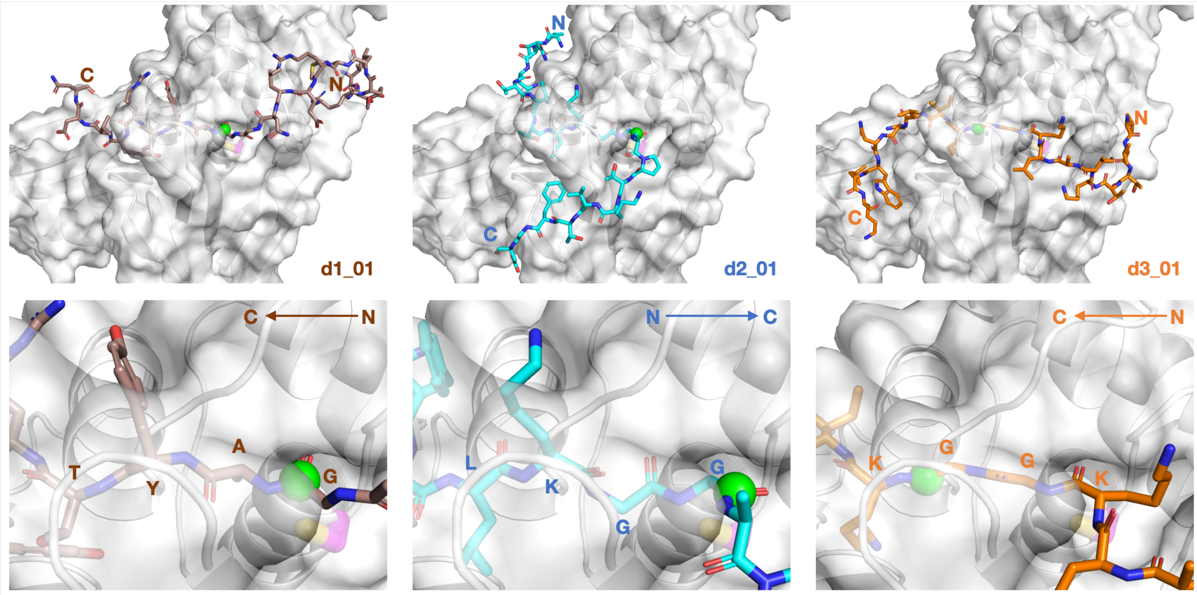

**Figure S1:** Top-ranked binding poses from AutoDock CrankPep (ADCP)[1] for the non-structural protein (nsp) peptides **1**, **2**, and **3**,[2] with the N- and C-terminals labeled. The P1 scissile amide carbonyl carbon is shown as a green sphere. The bottom row shows magnified views around the S4-S1 subsites, with each residue labeled and the N-/C-terminal peptide direction indicated.

**Table S1:** Deviations of P4-P1 Cα atom positions (compared to those in VIR251 in PDB 6WX4)[3] and root mean square deviation (RMSD) values for the 100 top-ranked solutions from ADCP[1] docking of peptides **1**, **2**, and **3**. Solutions that pass the filter (all four deviations within 2 Å) are in green, while reverse-bound solutions for peptide **3** that place Gly-Gly in S2-S1 are in orange.

| *Peptide 1* | | | | | |
|---|---|---|---|---|---|
| **Ranking** | **Deviation (Å)** | | | | **RMSD (Å)** |
| | *P4* | *P3* | *P2* | *P1* | |
| 1 | 17.719 | 13.064 | 7.576 | 1.417 | 11.662 |
| 2 | 13.666 | 8.103 | 7.947 | 9.937 | 10.177 |
| 3 | 18.191 | 12.249 | 5.673 | 0.914 | 11.335 |
| 4 | 17.483 | 13.424 | 7.936 | 1.724 | 11.745 |
| 5 | 6.922 | 0.564 | 0.830 | 0.933 | 3.528 |
| 6 | 5.533 | 0.709 | 0.507 | 0.679 | 2.821 |
| 7 | 31.863 | 29.764 | 30.688 | 30.875 | 30.807 |
| 8 | 32.268 | 25.838 | 25.227 | 19.290 | 26.064 |
| 9 | 17.040 | 10.391 | 4.744 | 2.624 | 10.340 |
| 10 | 25.450 | 22.260 | 19.441 | 14.266 | 20.765 |
| 11 | 17.323 | 11.897 | 5.589 | 0.513 | 10.876 |
| 12 | 17.102 | 17.723 | 14.568 | 10.983 | 15.325 |
| 13 | 16.175 | 9.664 | 4.350 | 3.021 | 9.786 |
| 14 | 28.948 | 26.041 | 23.593 | 21.042 | 25.078 |
| 15 | 27.717 | 25.926 | 25.155 | 22.993 | 25.504 |
| 16 | 1.399 | 1.056 | 0.932 | 0.820 | 1.074 |
| 17 | 18.706 | 20.327 | 15.567 | 15.243 | 17.591 |
| 18 | 21.105 | 19.892 | 19.762 | 18.792 | 19.905 |
| 19 | 29.355 | 30.862 | 30.857 | 31.482 | 30.649 |
| 20 | 6.181 | 0.631 | 0.771 | 0.980 | 3.169 |
| 21 | 31.736 | 27.538 | 22.098 | 20.509 | 25.857 |
| 22 | 31.833 | 27.764 | 22.741 | 21.052 | 26.194 |
| 23 | 34.470 | 32.255 | 32.393 | 31.838 | 32.755 |
| 24 | 27.746 | 25.320 | 21.586 | 20.650 | 23.996 |
| 25 | 30.216 | 30.142 | 26.237 | 24.589 | 27.904 |
| 26 | 20.123 | 18.924 | 13.763 | 15.361 | 17.237 |
| 27 | 31.402 | 26.795 | 24.208 | 21.814 | 26.296 |
| 28 | 29.973 | 27.894 | 24.003 | 25.343 | 26.902 |
| 29 | 29.966 | 27.525 | 24.618 | 25.643 | 27.015 |
| 30 | 29.860 | 32.634 | 30.199 | 29.803 | 30.646 |
| 31 | 15.447 | 9.754 | 13.325 | 11.967 | 12.792 |
| 32 | 32.234 | 25.686 | 24.249 | 18.890 | 25.708 |
| 33 | 19.313 | 18.175 | 16.090 | 11.954 | 16.622 |
| 34 | 31.060 | 26.630 | 23.550 | 21.756 | 25.990 |
| 35 | 30.071 | 27.136 | 24.316 | 23.089 | 26.292 |
| 36 | 29.439 | 30.119 | 26.938 | 23.384 | 27.597 |
| 37 | 28.470 | 23.456 | 20.084 | 14.413 | 22.203 |
| 38 | 10.893 | 4.367 | 3.239 | 8.895 | 7.539 |
| 39 | 34.381 | 31.994 | 32.021 | 31.557 | 32.507 |
| 40 | 22.689 | 23.255 | 25.439 | 27.057 | 24.672 |

| *Peptide 2* | | | | | |
|---|---|---|---|---|---|
| **Ranking** | **Deviation (Å)** | | | | **RMSD (Å)** |
| | *P4* | *P3* | *P2* | *P1* | |
| 1 | 0.555 | 0.638 | 0.616 | 0.445 | 0.568 |
| 2 | 0.546 | 0.646 | 0.614 | 0.383 | 0.557 |
| 3 | 13.315 | 8.287 | 3.650 | 4.068 | 8.304 |
| 4 | 18.696 | 18.812 | 14.792 | 13.679 | 16.653 |
| 5 | 23.861 | 20.525 | 16.843 | 12.695 | 18.944 |
| 6 | 0.625 | 0.544 | 0.682 | 0.756 | 0.656 |
| 7 | 12.635 | 8.459 | 8.616 | 12.230 | 10.666 |
| 8 | 37.973 | 33.416 | 31.510 | 27.160 | 32.746 |
| 9 | 1.239 | 1.065 | 0.387 | 0.476 | 0.872 |
| 10 | 29.630 | 26.500 | 23.873 | 20.509 | 25.351 |
| 11 | 13.272 | 11.405 | 10.346 | 9.681 | 11.258 |
| 12 | 21.599 | 18.559 | 18.474 | 15.576 | 18.674 |
| 13 | 28.438 | 23.705 | 20.353 | 17.767 | 22.916 |
| 14 | 4.552 | 2.358 | 2.622 | 2.189 | 3.080 |
| 15 | 17.537 | 11.482 | 9.744 | 8.085 | 12.244 |
| 16 | 24.056 | 20.174 | 16.827 | 17.105 | 19.757 |
| 17 | 30.482 | 30.402 | 30.337 | 31.269 | 30.625 |
| 18 | 17.712 | 17.224 | 11.983 | 9.072 | 14.459 |
| 19 | 27.797 | 21.113 | 16.537 | 12.024 | 20.227 |
| 20 | 30.334 | 29.271 | 25.621 | 20.723 | 26.753 |
| 21 | 38.124 | 33.812 | 32.604 | 27.433 | 33.212 |
| 22 | 11.333 | 10.431 | 9.675 | 9.663 | 10.298 |
| 23 | 20.064 | 14.913 | 14.285 | 11.854 | 15.569 |
| 24 | 12.906 | 11.650 | 5.525 | 1.711 | 9.162 |
| 25 | 25.053 | 22.424 | 20.733 | 15.158 | 21.155 |
| 26 | 23.032 | 19.377 | 15.844 | 11.336 | 17.927 |
| 27 | 41.061 | 38.721 | 38.606 | 34.456 | 38.285 |
| 28 | 29.117 | 28.875 | 25.548 | 19.656 | 26.080 |
| 29 | 34.970 | 27.981 | 25.388 | 18.620 | 27.373 |
| 30 | 36.073 | 29.573 | 28.459 | 24.032 | 29.847 |
| 31 | 21.790 | 17.449 | 15.413 | 15.124 | 17.646 |
| 32 | 17.676 | 17.419 | 15.769 | 11.733 | 15.829 |
| 33 | 29.421 | 25.649 | 22.888 | 18.467 | 24.435 |
| 34 | 18.540 | 18.569 | 15.021 | 13.975 | 16.654 |
| 35 | 37.315 | 31.198 | 27.664 | 21.703 | 30.008 |
| 36 | 19.309 | 19.874 | 14.822 | 13.717 | 17.144 |
| 37 | 33.199 | 27.787 | 25.667 | 20.789 | 27.227 |
| 38 | 0.745 | 0.526 | 0.701 | 1.004 | 0.763 |
| 39 | 13.785 | 8.636 | 4.172 | 3.511 | 8.578 |
| 40 | 28.610 | 25.173 | 22.788 | 19.112 | 24.170 |

| *Peptide 3* | | | | | |
|---|---|---|---|---|---|
| **Ranking** | **Deviation (Å)** | | | | **RMSD (Å)** |
| | *P4* | *P3* | *P2* | *P1* | |
| 1 | 11.898 | 7.809 | 1.303 | 6.506 | 7.851 |
| 2 | 23.181 | 16.380 | 10.563 | 4.065 | 15.279 |
| 3 | 21.375 | 17.295 | 14.571 | 11.297 | 16.552 |
| 4 | 20.953 | 15.725 | 19.831 | 18.326 | 18.811 |
| 5 | 29.626 | 27.569 | 22.286 | 20.224 | 25.216 |
| 6 | 16.900 | 17.037 | 16.321 | 17.637 | 16.980 |
| 7 | 15.259 | 8.595 | 3.408 | 4.180 | 9.162 |
| 8 | 34.033 | 32.487 | 30.714 | 26.474 | 31.056 |
| 9 | 27.685 | 26.231 | 23.336 | 22.612 | 25.052 |
| 10 | 40.175 | 35.169 | 31.132 | 25.347 | 33.402 |
| 11 | 27.886 | 26.856 | 27.162 | 26.899 | 27.204 |
| 12 | 15.469 | 9.031 | 3.766 | 3.816 | 9.349 |
| 13 | 17.990 | 16.932 | 13.953 | 12.753 | 15.554 |
| 14 | 20.601 | 20.131 | 19.251 | 20.858 | 20.219 |
| 15 | 23.907 | 17.900 | 18.867 | 20.651 | 20.459 |
| 16 | 15.972 | 10.648 | 4.278 | 2.913 | 9.941 |
| 17 | 18.113 | 18.376 | 15.694 | 13.355 | 16.510 |
| 18 | 28.422 | 23.714 | 23.635 | 21.246 | 24.393 |
| 19 | 24.405 | 21.608 | 18.356 | 15.256 | 20.200 |
| 20 | 15.195 | 8.625 | 3.605 | 3.970 | 9.138 |
| 21 | 29.275 | 27.440 | 22.585 | 20.852 | 25.273 |
| 22 | 27.867 | 27.591 | 23.014 | 22.175 | 25.294 |
| 23 | 23.167 | 19.542 | 17.246 | 15.462 | 19.073 |
| 24 | 22.629 | 18.852 | 17.926 | 13.329 | 18.483 |
| 25 | 44.617 | 41.349 | 39.960 | 36.192 | 40.642 |
| 26 | 44.700 | 41.208 | 39.550 | 35.504 | 40.376 |
| 27 | 22.662 | 16.161 | 10.385 | 4.213 | 15.003 |
| 28 | 23.907 | 18.682 | 15.821 | 13.244 | 18.346 |
| 29 | 22.391 | 15.797 | 13.637 | 11.300 | 16.314 |
| 30 | 32.843 | 30.974 | 32.785 | 32.223 | 32.215 |
| 31 | 22.878 | 21.267 | 20.096 | 21.467 | 21.450 |
| 32 | 24.833 | 22.145 | 19.383 | 14.554 | 20.583 |
| 33 | 23.664 | 23.184 | 27.114 | 28.236 | 25.641 |
| 34 | 22.422 | 22.828 | 23.759 | 26.057 | 23.808 |
| 35 | 24.498 | 25.178 | 27.726 | 29.359 | 26.762 |
| 36 | 22.776 | 16.403 | 10.496 | 3.984 | 15.115 |
| 37 | 13.372 | 10.599 | 8.571 | 8.791 | 10.511 |
| 38 | 15.813 | 14.702 | 13.782 | 14.830 | 14.799 |
| 39 | 35.034 | 35.680 | 34.739 | 32.458 | 34.499 |
| 40 | 35.552 | 33.192 | 32.097 | 29.156 | 32.580 |

| | | | | | | | | | | | | | | | | | | | |
|---|---|---|---|---|---|---|---|---|---|---|---|---|---|---|---|---|---|---|---|
| 41 | 21.999 | 20.491 | 16.194 | 14.350 | 18.520 | | 41 | 21.853 | 16.738 | 17.953 | 14.406 | 17.941 | | 41 | 17.951 | 18.080 | 16.320 | 13.311 | 16.527 |
| 42 | 29.654 | 29.093 | 24.277 | 23.912 | 26.865 | | 42 | 24.896 | 20.004 | 16.804 | 11.509 | 18.939 | | 42 | 29.979 | 25.653 | 23.150 | 20.742 | 25.115 |
| 43 | 29.046 | 25.650 | 22.535 | 22.643 | 25.110 | | 43 | 35.004 | 31.941 | 30.647 | 27.939 | 31.486 | | 43 | 15.998 | 9.970 | 4.367 | 2.993 | 9.790 |
| 44 | 17.333 | 12.783 | 7.424 | 1.329 | 11.410 | | 44 | 23.816 | 18.964 | 13.084 | 7.623 | 17.001 | | 44 | 19.676 | 17.881 | 15.871 | 14.505 | 17.096 |
| 45 | 29.782 | 30.492 | 27.242 | 24.646 | 28.135 | | 45 | 11.978 | 8.283 | 8.695 | 12.517 | 10.540 | | 45 | 33.535 | 34.767 | 33.957 | 30.967 | 33.337 |
| 46 | 2.977 | 2.419 | 3.897 | 4.738 | 3.618 | | 46 | 34.266 | 27.318 | 24.928 | 18.255 | 26.810 | | 46 | 18.557 | 19.095 | 17.362 | 16.749 | 17.965 |
| 47 | 23.776 | 17.510 | 14.650 | 7.907 | 16.949 | | 47 | 28.640 | 26.898 | 28.080 | 27.713 | 27.840 | | 47 | 20.068 | 18.848 | 15.834 | 14.661 | 17.490 |
| 48 | 18.357 | 12.796 | 6.210 | 1.131 | 11.625 | | 48 | 9.581 | 9.353 | 8.949 | 8.396 | 9.081 | | 48 | 9.365 | 3.661 | 9.214 | 15.616 | 10.367 |
| 49 | 28.459 | 25.941 | 22.612 | 22.700 | 25.047 | | 49 | 19.449 | 18.647 | 13.456 | 9.572 | 15.801 | | 49 | 18.545 | 17.061 | 14.079 | 12.946 | 15.818 |
| 50 | 28.902 | 27.319 | 27.084 | 26.634 | 27.498 | | 50 | 22.239 | 18.108 | 16.285 | 15.849 | 18.295 | | 50 | 28.853 | 30.215 | 29.093 | 26.412 | 28.677 |
| 51 | 16.950 | 12.619 | 7.377 | 1.610 | 11.220 | | 51 | 9.066 | 9.094 | 10.232 | 15.039 | 11.133 | | 51 | 16.569 | 9.669 | 10.577 | 9.529 | 11.945 |
| 52 | 18.827 | 18.476 | 15.425 | 15.600 | 17.155 | | 52 | 20.329 | 17.853 | 15.926 | 13.272 | 17.042 | | 52 | 15.162 | 8.471 | 3.408 | 4.337 | 9.111 |
| 53 | 22.520 | 20.156 | 14.798 | 11.219 | 17.736 | | 53 | 24.660 | 19.423 | 20.589 | 17.433 | 20.695 | | 53 | 14.336 | 10.410 | 5.476 | 2.397 | 9.349 |
| 54 | 17.635 | 12.795 | 7.484 | 2.020 | 11.563 | | 54 | 26.931 | 23.806 | 22.163 | 19.554 | 23.268 | | 54 | 22.998 | 19.905 | 15.490 | 10.235 | 17.817 |
| 55 | 6.025 | 1.049 | 1.360 | 0.606 | 3.147 | | 55 | 28.746 | 28.796 | 25.555 | 20.933 | 26.205 | | 55 | 19.235 | 19.814 | 16.979 | 12.397 | 17.353 |
| 56 | 34.977 | 34.766 | 32.347 | 32.463 | 33.661 | | 56 | 18.878 | 17.061 | 16.788 | 12.578 | 16.489 | | 56 | 11.885 | 7.879 | 1.406 | 6.353 | 7.837 |
| 57 | 23.732 | 23.631 | 23.334 | 23.154 | 23.464 | | 57 | 21.119 | 18.738 | 18.268 | 18.582 | 19.210 | | 57 | 20.482 | 15.989 | 11.219 | 7.434 | 14.631 |
| 58 | 19.883 | 18.503 | 15.127 | 15.560 | 17.383 | | 58 | 27.911 | 32.009 | 27.883 | 25.605 | 28.446 | | 58 | 10.578 | 4.250 | 9.663 | 13.476 | 10.062 |
| 59 | 20.828 | 19.422 | 15.715 | 11.400 | 17.234 | | 59 | 29.455 | 26.473 | 23.649 | 20.074 | 25.153 | | 59 | 24.914 | 22.227 | 19.383 | 15.904 | 20.877 |
| 60 | 24.589 | 23.006 | 24.288 | 24.204 | 24.029 | | 60 | 22.431 | 21.058 | 22.253 | 21.483 | 21.813 | | 60 | 22.795 | 15.683 | 9.992 | 3.958 | 14.841 |
| 61 | 21.450 | 18.219 | 16.360 | 13.440 | 17.609 | | 61 | 17.976 | 15.879 | 12.612 | 9.718 | 14.394 | | 61 | 22.172 | 22.355 | 24.973 | 26.153 | 23.974 |
| 62 | 19.482 | 21.221 | 15.798 | 14.975 | 18.053 | | 62 | 0.762 | 0.717 | 0.910 | 0.753 | 0.789 | | 62 | 24.748 | 21.858 | 22.862 | 20.350 | 22.511 |
| 63 | 28.894 | 28.875 | 27.866 | 26.015 | 27.937 | | 63 | 36.155 | 30.113 | 28.964 | 25.422 | 30.410 | | 63 | 12.526 | 7.426 | 8.893 | 10.875 | 10.117 |
| 64 | 22.148 | 16.510 | 14.849 | 15.612 | 17.517 | | 64 | 26.073 | 22.619 | 23.296 | 23.210 | 23.837 | | 64 | 14.472 | 11.588 | 9.870 | 8.401 | 11.310 |
| 65 | 30.362 | 33.225 | 30.485 | 27.293 | 30.414 | | 65 | 29.145 | 25.869 | 23.702 | 19.528 | 24.808 | | 65 | 18.018 | 17.833 | 12.184 | 10.912 | 15.085 |
| 66 | 30.776 | 30.913 | 26.655 | 24.429 | 28.329 | | 66 | 0.673 | 0.699 | 0.662 | 0.965 | 0.760 | | 66 | 18.800 | 18.891 | 15.754 | 12.679 | 16.728 |
| 67 | 10.358 | 10.700 | 16.707 | 21.025 | 15.354 | | 67 | 24.837 | 18.770 | 18.748 | 16.523 | 19.960 | | 67 | 25.175 | 20.676 | 17.202 | 13.105 | 19.551 |
| 68 | 22.208 | 21.067 | 18.147 | 15.014 | 19.312 | | 68 | 28.037 | 22.982 | 24.016 | 19.861 | 23.903 | | 68 | 20.348 | 15.872 | 16.241 | 13.253 | 16.624 |
| 69 | 29.541 | 25.938 | 22.445 | 23.849 | 25.583 | | 69 | 0.954 | 0.818 | 0.796 | 0.585 | 0.800 | | 69 | 16.008 | 11.465 | 11.262 | 13.937 | 13.311 |
| 70 | 28.591 | 24.011 | 21.080 | 15.032 | 22.717 | | 70 | 29.292 | 27.457 | 25.633 | 24.716 | 26.832 | | 70 | 23.274 | 19.565 | 17.454 | 15.895 | 19.247 |
| 71 | 22.100 | 20.028 | 19.769 | 19.011 | 20.260 | | 71 | 16.651 | 13.998 | 13.291 | 12.763 | 14.255 | | 71 | 21.379 | 18.088 | 16.459 | 13.842 | 17.655 |
| 72 | 18.897 | 14.300 | 9.538 | 10.896 | 13.886 | | 72 | 23.389 | 17.825 | 15.693 | 12.069 | 17.725 | | 72 | 13.018 | 7.663 | 6.783 | 9.846 | 9.633 |
| 73 | 30.024 | 27.437 | 23.294 | 22.186 | 25.928 | | 73 | 23.414 | 20.060 | 16.998 | 15.048 | 19.144 | | 73 | 22.374 | 17.668 | 12.173 | 13.009 | 16.809 |
| 74 | 16.973 | 13.134 | 7.715 | 1.772 | 11.437 | | 74 | 35.819 | 30.127 | 29.447 | 25.548 | 30.457 | | 74 | 22.923 | 19.600 | 15.574 | 10.362 | 17.745 |
| 75 | 21.811 | 17.673 | 14.451 | 12.799 | 17.035 | | 75 | 0.616 | 0.929 | 1.324 | 1.358 | 1.100 | | 75 | 15.969 | 10.744 | 4.330 | 2.964 | 9.974 |
| 76 | 22.697 | 16.972 | 16.652 | 17.556 | 18.633 | | 76 | 36.136 | 37.749 | 34.738 | 32.044 | 35.229 | | 76 | 22.043 | 20.931 | 17.130 | 13.422 | 18.692 |
| 77 | 14.722 | 11.995 | 11.241 | 13.165 | 12.848 | | 77 | 35.228 | 31.906 | 31.795 | 28.447 | 31.934 | | 77 | 29.216 | 25.742 | 23.515 | 18.497 | 24.552 |
| 78 | 20.211 | 19.284 | 16.563 | 15.319 | 17.954 | | 78 | 33.054 | 34.975 | 33.758 | 31.149 | 33.263 | | 78 | 11.896 | 8.710 | 13.649 | 15.934 | 12.821 |
| 79 | 7.662 | 8.211 | 13.077 | 18.129 | 12.508 | | 79 | 19.717 | 16.627 | 16.026 | 16.071 | 17.178 | | 79 | 21.040 | 17.156 | 14.913 | 15.772 | 17.379 |
| 80 | 23.075 | 23.584 | 26.542 | 27.221 | 25.170 | | 80 | 28.899 | 29.363 | 26.879 | 25.853 | 27.786 | | 80 | 12.987 | 9.846 | 7.451 | 8.250 | 9.864 |
| 81 | 2.517 | 2.559 | 3.597 | 4.657 | 3.447 | | 81 | 30.093 | 29.367 | 24.445 | 21.578 | 26.604 | | 81 | 24.373 | 21.063 | 19.039 | 19.258 | 21.042 |
| 82 | 23.282 | 23.631 | 22.898 | 25.139 | 23.753 | | 82 | 29.200 | 28.013 | 23.478 | 21.799 | 25.806 | | 82 | 21.434 | 14.656 | 16.151 | 15.620 | 17.169 |
| 83 | 22.426 | 23.086 | 25.967 | 26.888 | 24.664 | | 83 | 13.186 | 11.383 | 10.268 | 9.852 | 11.247 | | 83 | 14.957 | 8.277 | 3.525 | 4.132 | 8.969 |
| 84 | 0.869 | 1.315 | 1.760 | 2.971 | 1.898 | | 84 | 38.157 | 35.915 | 31.511 | 25.938 | 33.210 | | 84 | 40.267 | 35.113 | 31.168 | 25.455 | 33.443 |
| 85 | 33.064 | 32.368 | 27.822 | 25.192 | 29.789 | | 85 | 20.008 | 20.531 | 15.623 | 14.318 | 17.825 | | 85 | 21.326 | 16.886 | 15.093 | 15.389 | 17.353 |
| 86 | 15.788 | 10.397 | 13.368 | 12.356 | 13.122 | | 86 | 22.414 | 21.413 | 20.852 | 20.397 | 21.282 | | 86 | 31.943 | 30.820 | 28.342 | 24.297 | 28.999 |
| 87 | 18.188 | 12.302 | 5.782 | 0.741 | 11.359 | | 87 | 22.991 | 16.185 | 13.104 | 6.818 | 15.881 | | 87 | 12.227 | 8.545 | 10.873 | 10.704 | 10.669 |
| 88 | 26.414 | 25.839 | 21.916 | 20.254 | 23.748 | | 88 | 23.757 | 19.784 | 15.347 | 10.038 | 17.973 | | 88 | 12.386 | 7.134 | 8.384 | 13.237 | 10.605 |
| 89 | 22.957 | 18.111 | 14.596 | 12.203 | 17.443 | | 89 | 35.981 | 29.233 | 25.258 | 18.578 | 27.984 | | 89 | 34.346 | 35.043 | 33.712 | 32.345 | 33.876 |
| 90 | 29.125 | 26.231 | 23.740 | 21.325 | 25.272 | | 90 | 26.829 | 23.477 | 24.243 | 23.460 | 24.541 | | 90 | 12.972 | 10.037 | 10.174 | 9.871 | 10.839 |
| 91 | 18.260 | 12.371 | 5.664 | 0.985 | 11.397 | | 91 | 14.824 | 13.455 | 17.307 | 18.009 | 16.005 | | 91 | 24.566 | 21.802 | 18.858 | 15.277 | 20.419 |
| 92 | 18.579 | 17.262 | 14.294 | 13.134 | 15.969 | | 92 | 29.372 | 26.222 | 23.826 | 20.238 | 25.137 | | 92 | 21.103 | 17.107 | 15.082 | 13.656 | 16.970 |
| 93 | 10.793 | 4.126 | 3.746 | 9.602 | 7.742 | | 93 | 22.013 | 22.121 | 22.860 | 25.346 | 23.124 | | 93 | 27.402 | 24.602 | 21.427 | 16.428 | 22.831 |
| 94 | 19.196 | 17.789 | 15.619 | 11.840 | 16.348 | | 94 | 34.660 | 28.950 | 28.664 | 23.895 | 29.292 | | 94 | 13.994 | 14.006 | 13.201 | 14.261 | 13.871 |
| 95 | 29.126 | 30.097 | 26.909 | 23.291 | 27.481 | | 95 | 13.801 | 8.487 | 11.597 | 11.875 | 11.597 | | 95 | 23.152 | 17.972 | 16.728 | 12.874 | 18.059 |
| 96 | 22.880 | 17.966 | 12.117 | 12.871 | 17.020 | | 96 | 22.596 | 19.734 | 16.859 | 13.379 | 18.461 | | 96 | 13.646 | 10.899 | 10.554 | 8.872 | 11.125 |
| 97 | 39.615 | 36.685 | 32.698 | 28.006 | 34.528 | | 97 | 0.880 | 1.168 | 0.999 | 0.996 | 1.016 | | 97 | 16.995 | 17.135 | 16.648 | 17.737 | 17.133 |
| 98 | 35.822 | 36.519 | 34.517 | 32.731 | 34.927 | | 98 | 25.711 | 22.574 | 23.292 | 22.684 | 23.600 | | 98 | 21.506 | 16.417 | 18.341 | 14.722 | 17.924 |
| 99 | 27.975 | 26.127 | 24.801 | 22.370 | 25.400 | | 99 | 26.936 | 24.052 | 25.067 | 23.187 | 24.850 | | 99 | 23.718 | 21.634 | 22.663 | 21.199 | 22.325 |
| 100 | 18.301 | 12.586 | 6.158 | 1.055 | 11.537 | | 100 | 17.729 | 13.154 | 8.905 | 4.514 | 12.114 | | 100 | 18.771 | 17.789 | 15.972 | 14.502 | 16.839 |

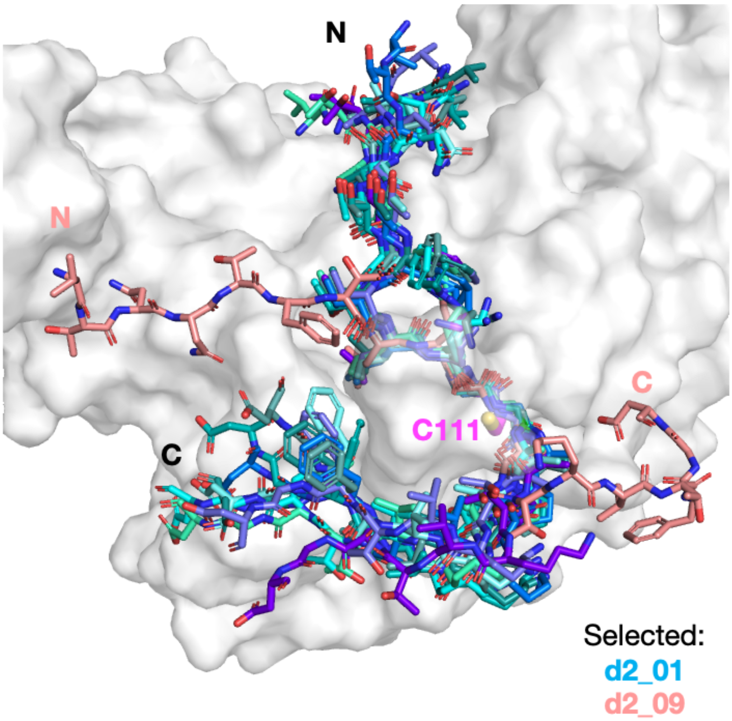

**Figure S2:** Docked conformations of peptide **2** that passed the applied filter, *i.e.*, the four Cα atoms of P4-P1 have to be within 2 Å of the corresponding Cα atoms of VIR251.[3] The poses include ranked solutions 01 (cyan), 02 (green-cyan), 06 (aquamarine), 09 (salmon), 38 (teal), 62 (deep teal), 66 (light teal), 69 (marine), 75 (slate), and 97 (purple blue). Except for d2_09, which adopts a unique extended conformation, d2_01 is representative of all the remaining poses.

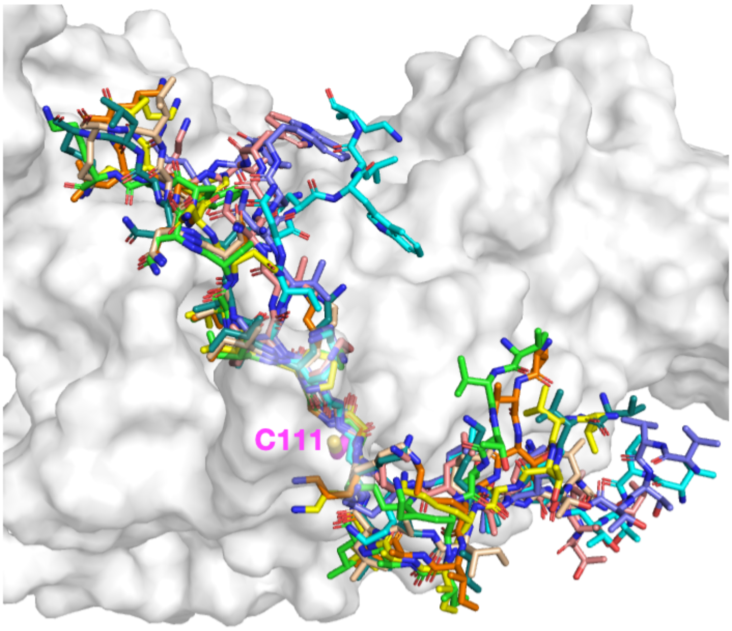

**Figure S3:** Docked reverse-bound poses of peptide **3**, which position the original P2-Gly residue in the S1 subsite, P1-Gly in the S2 subsite, P1′-Lys in the S3 subsite, and P2′-Ile in the S4 subsite. The poses include ranked solutions 07 (orange), 12 (green), 16 (cyan), 20 (yellow), 43 (salmon), 52 (teal), 75 (slate), and 83 (wheat). The highest ranked solution among them, d3_07, was selected as being representative and was used for subsequent modeling.

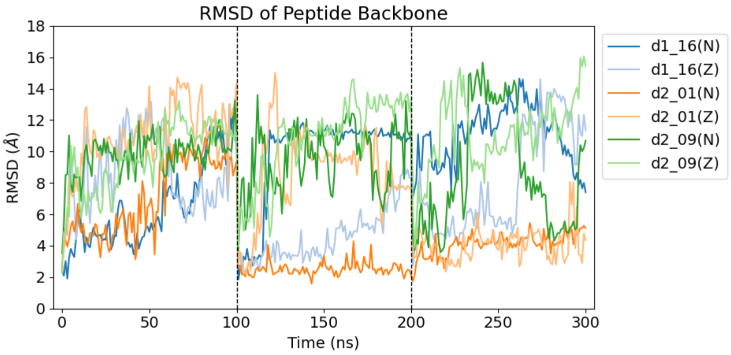

**Figure S4:** RMSD of peptide backbone atoms (N, Cα, C) in the combined 3 × 100 ns molecular dynamics (MD) trajectories (fitted based on PL[pro] backbone) initiated from the d1_16, d2_01, and d2_09 poses in either the neutral (N) or zwitterionic (Z) state of Cys111-His272, relative to their corresponding starting peptide poses.

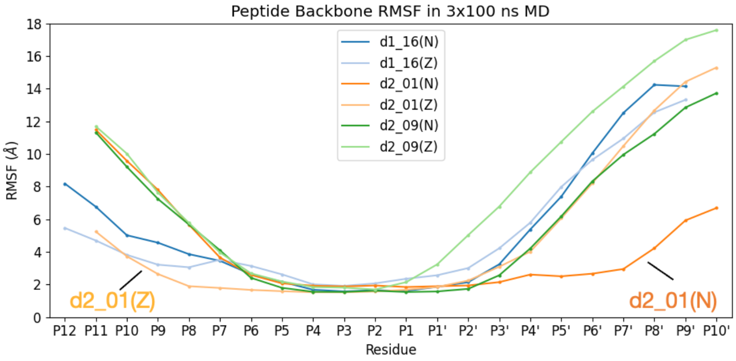

**Figure S5:** Per-residue root mean square fluctuation (RMSF) of peptide backbone atoms in the combined 3 × 100 ns MD (fitted based on PL[pro] backbone) initiated from the d1_16, d2_01, and d2_09 poses in either the N or Z state of Cys111-His272. Note that the C-terminal $NH_2$ group is treated as a separate residue.

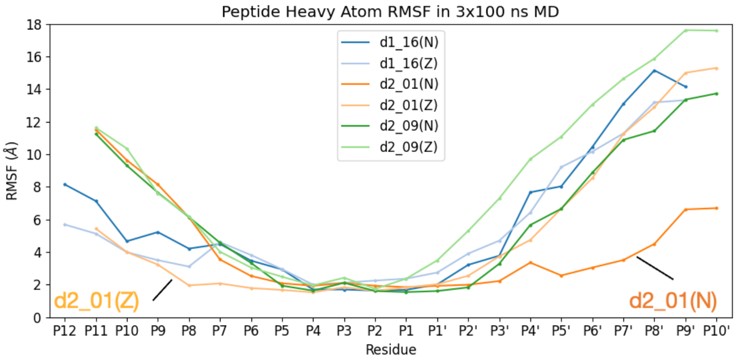

**Figure S6:** Per-residue RMSF of peptide non-hydrogen atoms in the combined 3 × 100 ns MD (fitted based on PL[pro] backbone) initiated from the d1_16, d2_01, and d2_09 poses in either the N or Z state of Cys111-His272.

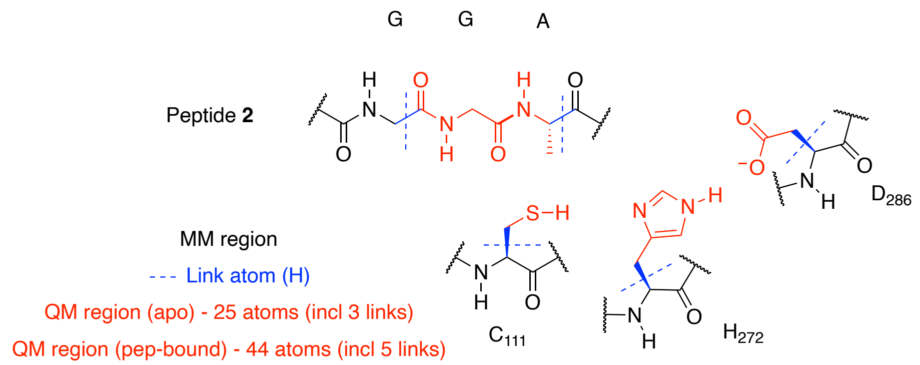

**Figure S7:** Definition of the quantum mechanical (QM) region in the quantum mechanics/molecular mechanics-umbrella sampling (QM/MM-US) calculations for proton transfer processes in the Cys111-His272-Asp286 catalytic triad.

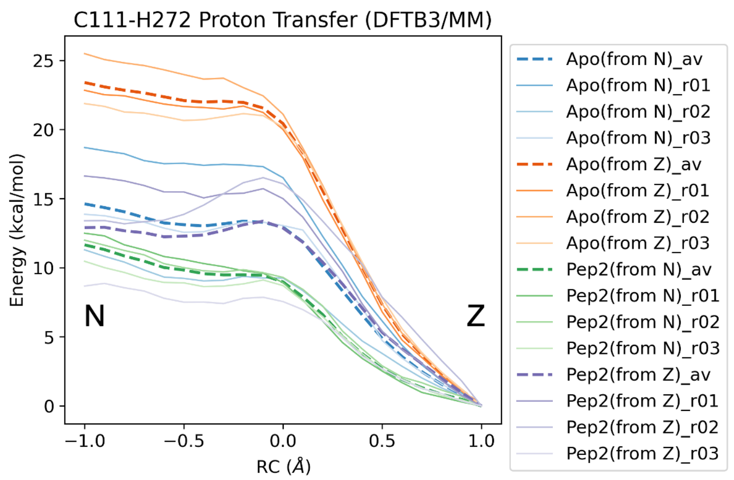

**Figure S8:** DFTB3/MM-US free energy profiles from the weighted histogram analysis method (WHAM)[4] for the proton transfer between Cys111 and His272, with reaction coordinate (RC) = -1 being the N state and +1 being the Z state. The energy profiles averaged from three replicas are shown as dotted lines.

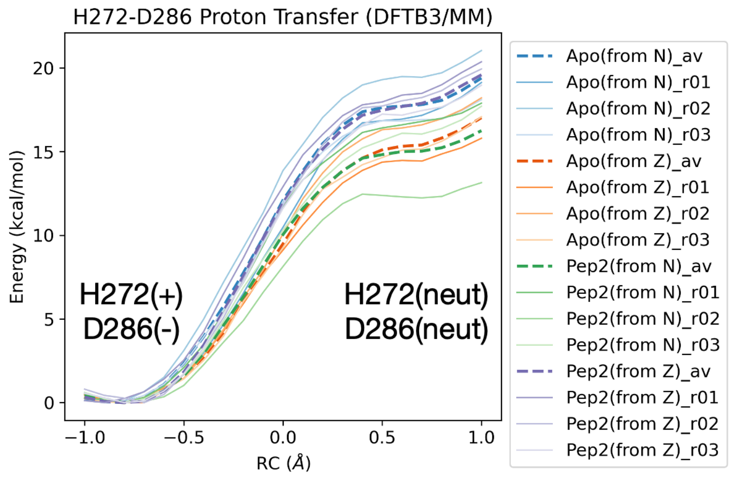

**Figure S9:** WHAM[4]-derived DFTB3/MM-US free energy profiles for the proton transfer between His272 and Asp286, between RC = -1 (doubly protonated His272 and deprotonated Asp286) and +1 (neutral His272 and neutral Asp286). The energy profiles averaged from three replicas are shown as dotted lines.

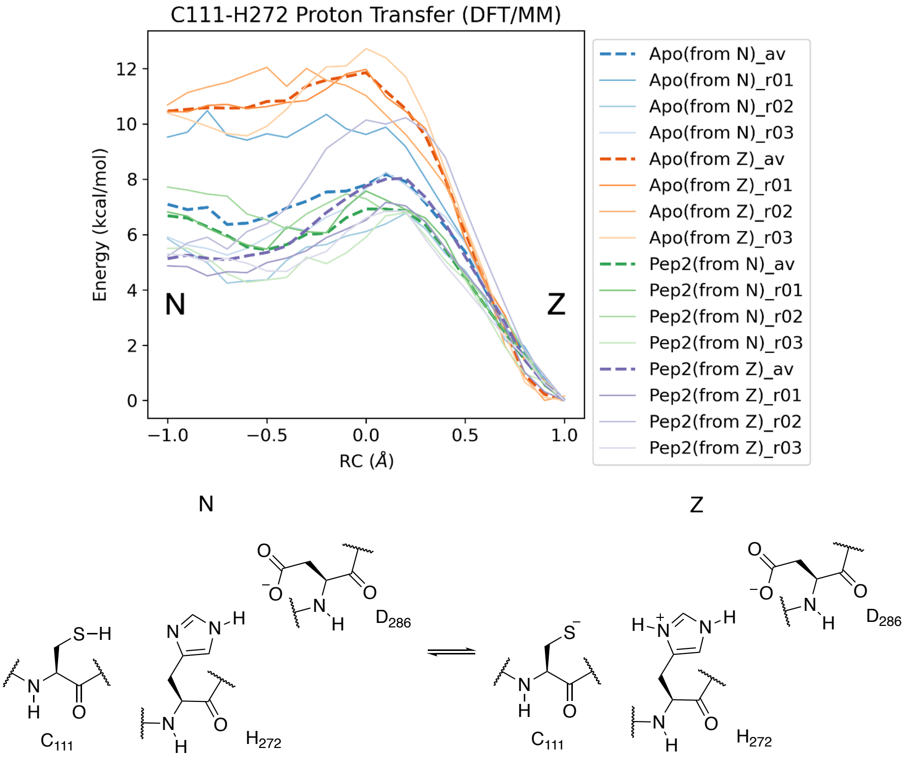

**Figure S10:** WHAM[4]-derived PBE0-D3BJ/6-31G(d)/MM-US free energy profiles for the proton transfer between Cys111 and His272, with RC = -1 being the N state and +1 being the Z state. The energy profiles averaged from three replicas are shown as dotted lines.

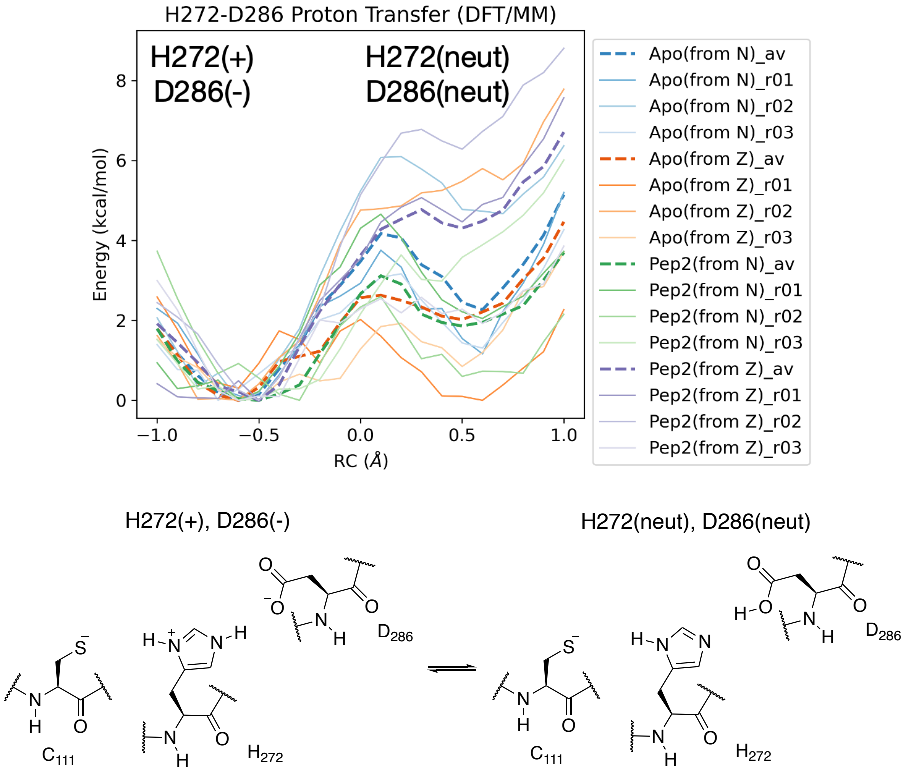

**Figure S11:** WHAM[4]-derived PBE0-D3BJ/6-31G(d)/MM-US free energy profiles for the proton transfer between His272 and Asp286, between RC = -1 (doubly protonated His272 and deprotonated Asp286) and +1 (neutral His272 and neutral Asp286). The energy profiles averaged from three replicas are shown as dotted lines.

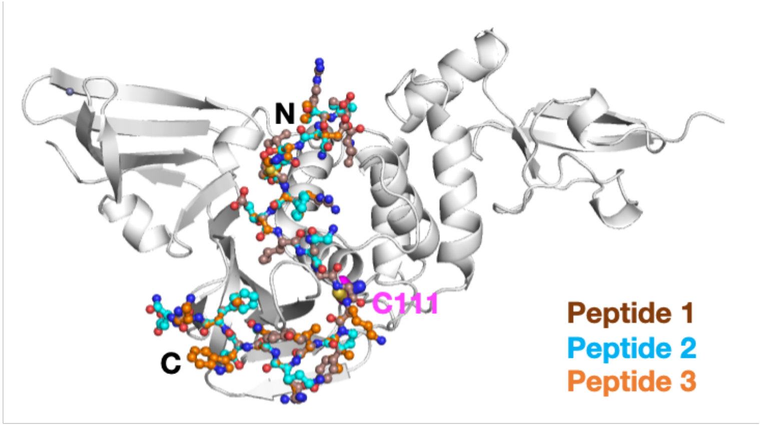

**Figure S12:** Starting conformations of PL$^{pro}$ complexed with each of the three nsp oligopeptides **1**, **2**, and **3**, based on the d2_01 conformation.

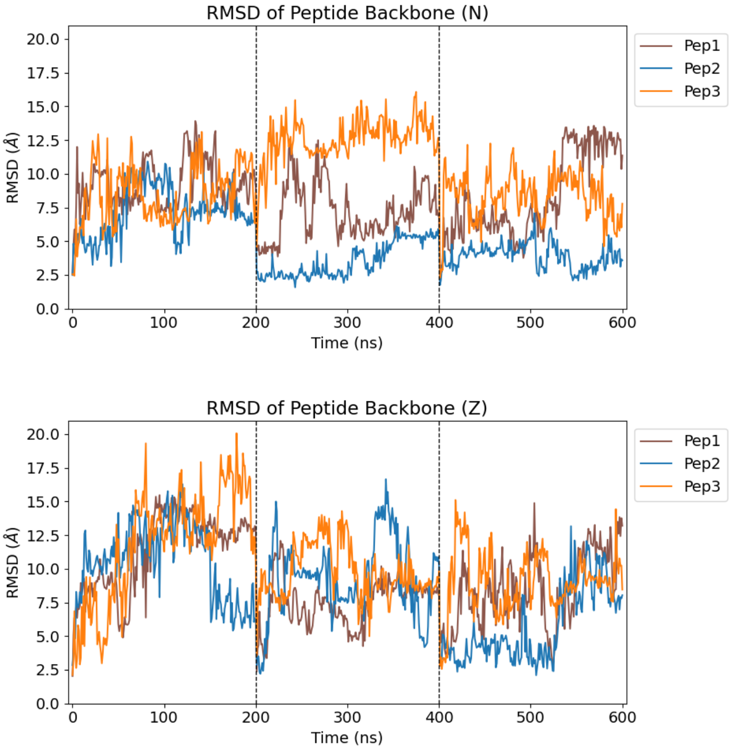

**Figure S13:** RMSD of peptide backbone atoms in the combined 3 × 200 ns MD trajectories (fitted based on PL$^{pro}$ backbone) for the three nsp peptides initiated from the d2_01 conformation, in the (top) N and (bottom) Z states of Cys111-His272, relative to their corresponding starting conformations.

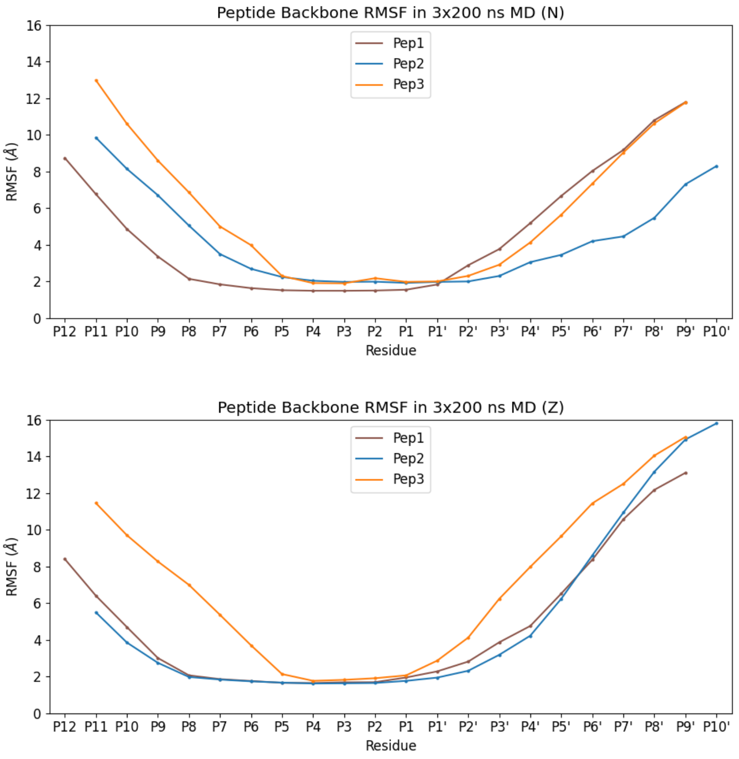

**Figure S14:** Per-residue RMSF of peptide backbone atoms in the combined 3 × 200 ns MD (fitted based on PL$^{pro}$ backbone) for the three nsp peptides initiated from the d2_01 conformation, in the (top) N and (bottom) Z states of Cys111-His272. Note that the C-terminal $NH_2$ group is treated as a separate residue.

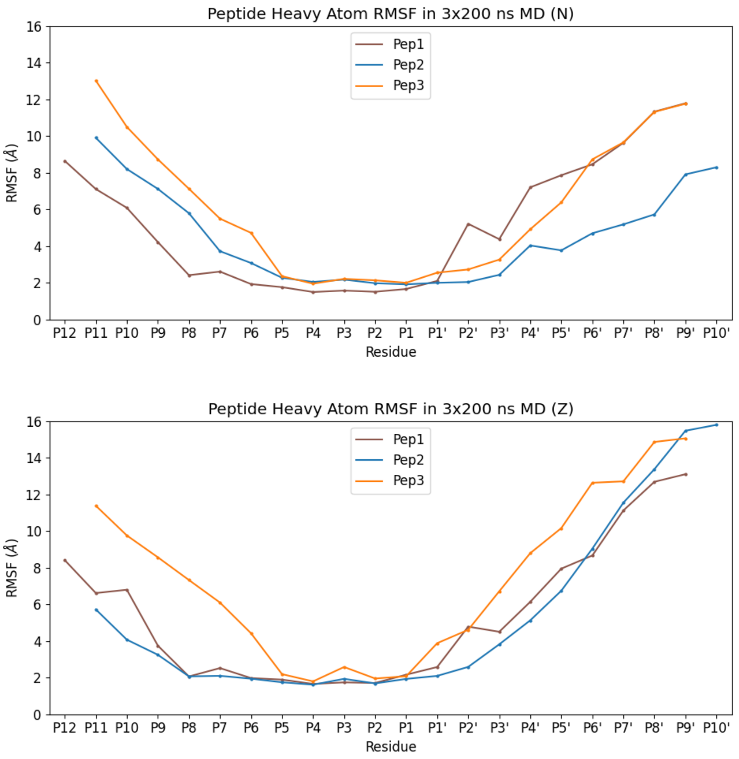

**Figure S15:** Per-residue RMSF of peptide non-hydrogen atoms in the combined 3 × 200 ns MD (fitted based on PL$^{pro}$ backbone) for the three nsp peptides initiated from the d2_01 conformation, in the (top) N and (bottom) Z states of Cys111-His272.

**Table S2:** PL$^{pro}$-peptide hydrogen bonds (HBs) of significance (occurrence ≥ 25% out of 600 frames analyzed every ns) observed over the combined 3 × 200 ns MD for each of the three nsp peptides **1**, **2**, and **3**, all initiated from the d2_01 conformation, in the (left) N and (right) Z states of Cys111-His272. HB occurrences involving equivalent carboxylate oxygens (in Asp and Glu sidechains) are combined, while salt bridges involving carboxylate oxygens and Arg sidechain guanidino group are combined and halved to avoid double counting. Peptide residues are italicized, while PL$^{pro}$ residues are not. Outside the core P5-P1′ region (see **Figure 3a**), N-terminal HBs are in blue and C-terminal HBs are in red.

| Donor | Acceptor | % | | Donor | Acceptor | % |
|---|---|---|---|---|---|---|
| Pep 1 (N) | P5-P1′=8-13 | | | Pep 1 (Z) | | |
| *GLY12N* | GLY271O | 96 | | *GLU8N* | ASP164OD | 97 |
| *GLY11N* | GLY163O | 95 | | *ASN10ND2* | TYR268O | 96 |
| *GLU8N* | ASP164OD | 94 | | *GLY11N* | GLY163O | 95 |
| GLY163N | *GLY11O* | 93 | | *LEU9N* | ASP164OD | 83 |
| *ASN10ND2* | TYR268O | 91 | | GLY271N | *ASN10O* | 80 |
| *LEU9N* | ASP164OD | 80 | | GLY163N | *GLY11O* | 80 |
| ARG166NH | *GLU8OE* | 80 | | ARG166NH | *GLU8OE* | 80 |
| *ASN10N* | TYR264OH | 74 | | *ASN10N* | TYR264OH | 79 |
| GLY271N | *ASN10O* | 74 | | *GLY12N* | GLY271O | 76 |
| TRP106NE1 | *GLY12O* | 59 | | *ALA13N* | GLY271O | 48 |
| *ALA13N* | GLY271O | 36 | | *THR15OG1* | ASP286OD | 35 |
| *ARG3NH* | GLU203OE | 32 | | *LEU5N* | TYR171OH | 28 |
| *ARG7NH* | GLU161OE | 25 | | *ARG3NH* | GLU203OE | 25 |
| Pep 2 (N) | P5-P1′=7-12 | | | Pep 2 (Z) | | |
| *GLY11N* | GLY271O | 95 | | *THR7N* | ASP164OD | 100 |
| TRP106NE1 | *GLY11O* | 95 | | *GLY10N* | GLY163O | 94 |
| GLY163N | *GLY10O* | 92 | | *LEU8N* | ASP164OD | 86 |
| *GLY10N* | GLY163O | 92 | | GLY271N | *LYS9O* | 80 |
| *THR7N* | ASP164OD | 70 | | GLY163N | *GLY10O* | 74 |
| GLY266N | *VAL16O* | 69 | | *LYS9N* | TYR264OH | 68 |
| THR265OG1 | *THR14O* | 68 | | *GLY11N* | GLY271O | 66 |
| *LEU8N* | ASP164OD | 64 | | *ASN4ND2* | GLN174OE1 | 58 |
| GLY271N | *LYS9O* | 56 | | *ASN4N* | TYR171OH | 45 |
| *LYS9N* | TYR264OH | 44 | | HIP272ND1 | *GLY11O* | 42 |
| GLN250N | *GLY19O* | 41 | | *ALA12N* | GLY271O | 39 |
| *ALA12N* | GLY271O | 36 | | ASN109ND2 | *GLY11O* | 35 |
| ARG166NH | *THR7OG1* | 33 | | GLN174NE2 | *ASN4OD1* | 33 |
| GLN250NE2 | *PHE18O* | 28 | | ARG166NH | *THR7OG1* | 29 |
| *ASN4ND2* | GLN174OE1 | 26 | | GLN174NE2 | *THR2O* | 27 |
| Pep 3 (N) | P5-P1′=7-12 | | | Pep 3 (Z) | | |
| TRP106NE1 | *GLY11O* | 96 | | *GLY10N* | GLY163O | 88 |
| GLY163N | *GLY10O* | 94 | | *GLY11N* | GLY271O | 72 |
| *GLY11N* | GLY271O | 94 | | GLY163N | *GLY10O* | 58 |
| *GLY10N* | GLY163O | 92 | | GLY271N | *LYS9O* | 38 |
| GLY271N | *LYS9O* | 60 | | *LYS12N* | GLY271O | 34 |
| *LYS9N* | TYR268O | 35 | | *LYS9N* | TYR264OH | 32 |
| *LEU8N* | ASP164OD | 28 | | *LYS9N* | TYR268O | 28 |
| *ALA7N* | ASP164OD | 28 | | | | |

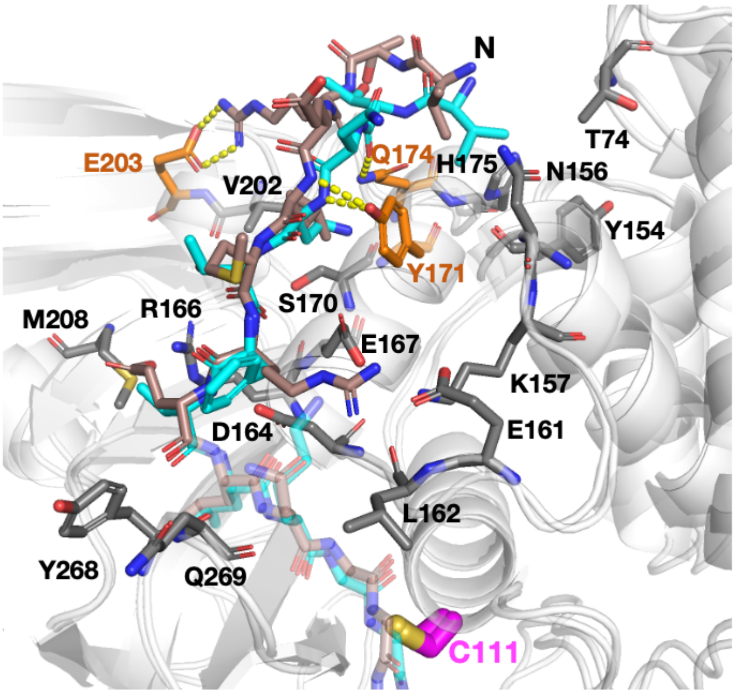

**Figure S16:** View of clustering-derived representative structures of Z-state PL$^{pro}$ (white cartoon) complexed with peptides **1** (brown) or **2** (cyan), with a focus on the interactions in the N-terminal region outside the consensus P4-P1 sequence. PL$^{pro}$ residues involved in HBs (yellow dotted lines; see **Table S2**; not all significant HBs are present in these snapshots) are shown as orange sticks. Other residues in close contact (within 4 Å) with the peptide residues N-terminal of P4 are in grey. The active site Cys111 is shown as magenta sticks. Clustering was performed using a 3 Å RMSD cut-off on the peptide non-hydrogen atoms using gmx cluster (gromos algorithm).[5,6] Out of 601 frames extracted every ns from the combined 3 × 200 ns MD (including *t* = 0), the number of (multimembered) clusters obtained for peptides **1**, **2**, and **3** are respectively 291 (81), 233 (76), and 361 (77).

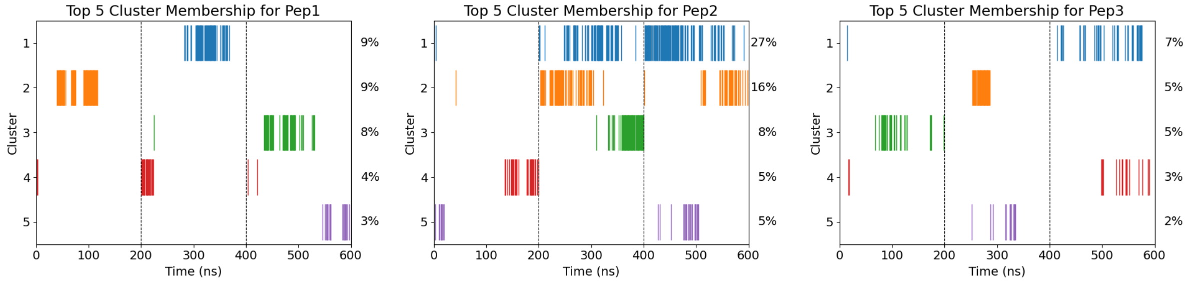

**Figure S17:** Cluster membership of frames extracted every ns from the combined 3 × 200 ns MD (fitted using PL$^{pro}$ backbone) of each of the three PL$^{pro}$-peptide (N state) complexes, with clustering performed using a 3 Å RMSD cut-off on the peptide backbone atoms (N, Cα, C) using gmx cluster (gromos algorithm).[5,6] Only the 5 clusters with the highest occupancy percentages are shown. Out of 601 frames, the number of (multimembered) clusters obtained for peptides **1**, **2**, and **3** are respectively 157 (73), 66 (41), and 201 (77).

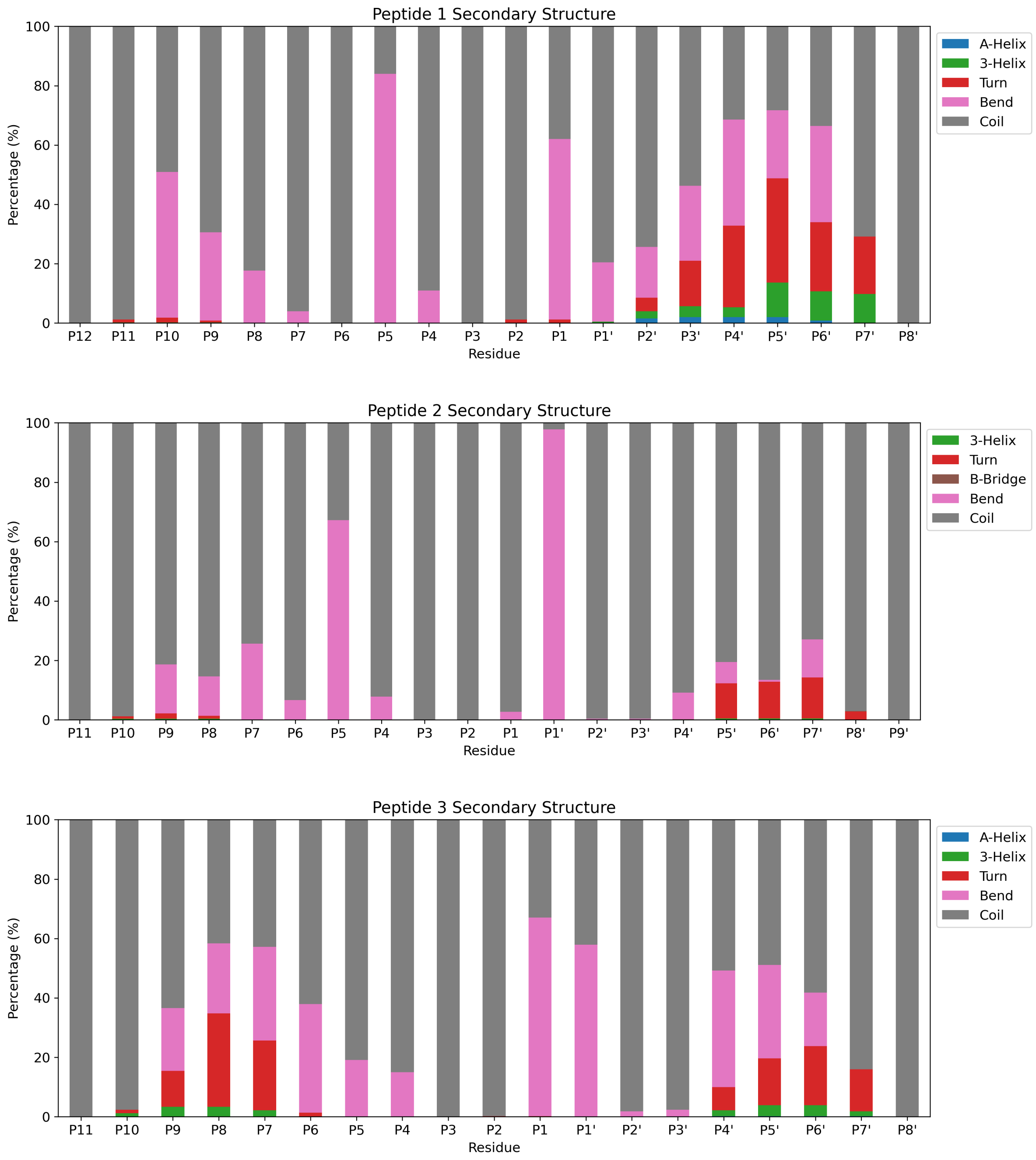

**Figure S18:** Proportion of secondary structure adopted by every peptide residue over the combined 3 × 200 ns MD of PL$^{pro}$-peptide (N state) complexes, assigned by DSSP (version 2.0.4).[7,8] Frames were analyzed every ns.

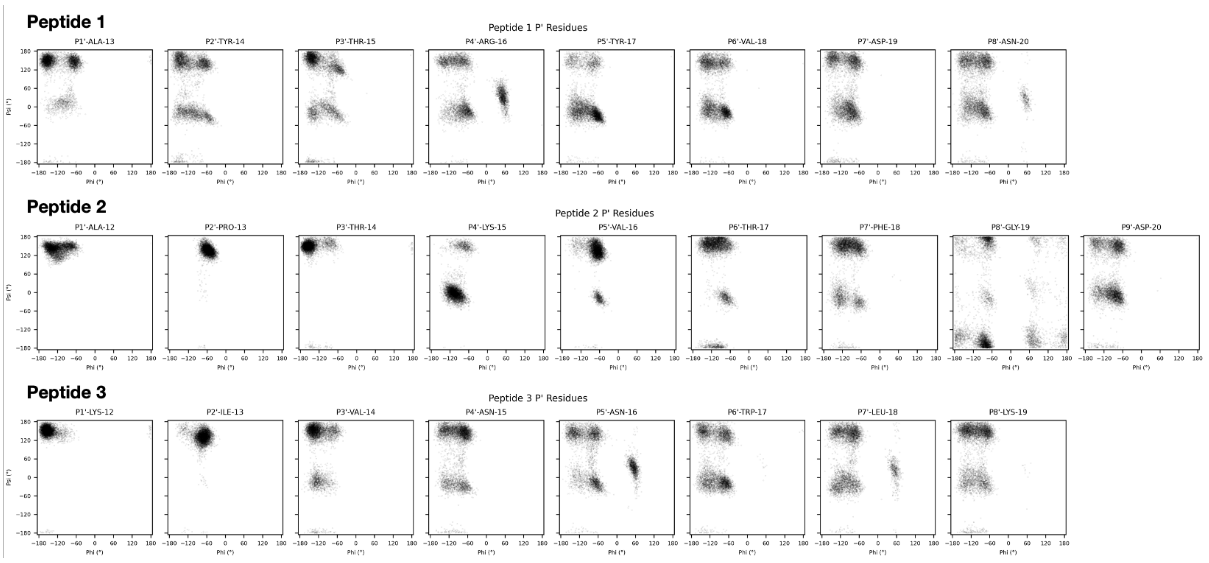

**Figure S19**: Ramachandran plots of every peptide P′ residue over the combined 3 × 200 ns MD of PL$^{pro}$-peptide (N state) complexes. Frames were analyzed every 0.1 ns.

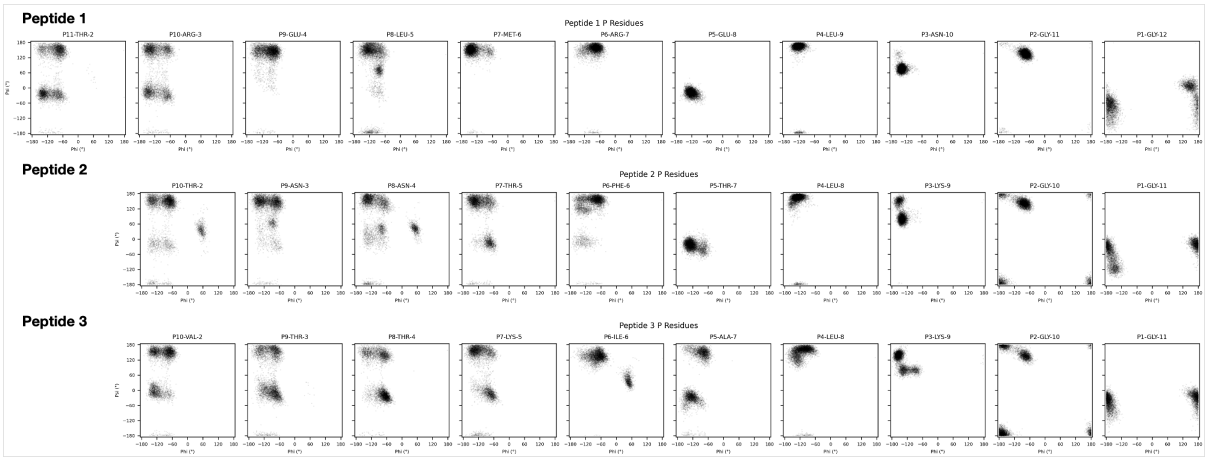

**Figure S20**: Ramachandran plots of every peptide P residue over the combined 3 × 200 ns MD of PL$^{pro}$-peptide (N state) complexes. Frames were analyzed every 0.1 ns.

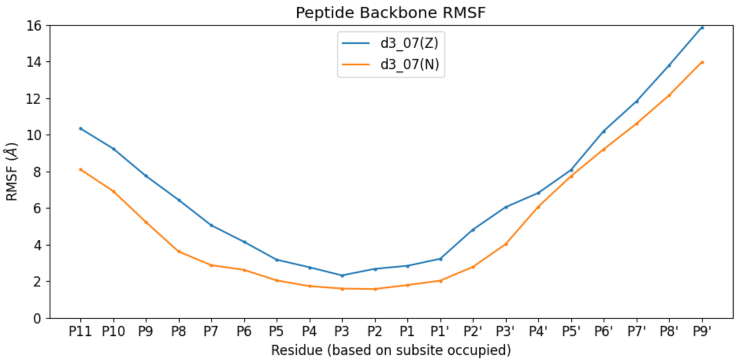

**Figure S21:** Per-residue RMSF of peptide backbone atoms in the combined 3 × 200 ns MD (fitted based on PL$^{pro}$ backbone) for peptide **3** initiated from the d3_07 conformation, in the N (orange) and Z (blue) states of Cys111-His272. Note that the C-terminal $NH_2$ cap is treated as a separate residue (P11).

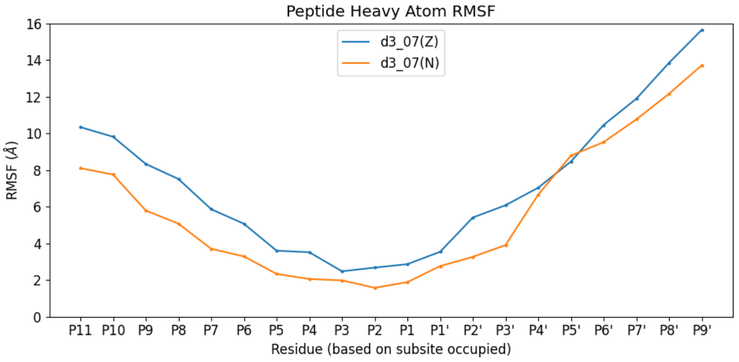

**Figure S22:** Per-residue RMSF of peptide non-hydrogen atoms in the combined 3 × 200 ns MD (fitted based on PL$^{pro}$ backbone) for peptide **3** initiated from the d3_07 conformation, in the N (orange) and Z (blue) states of Cys111-His272.

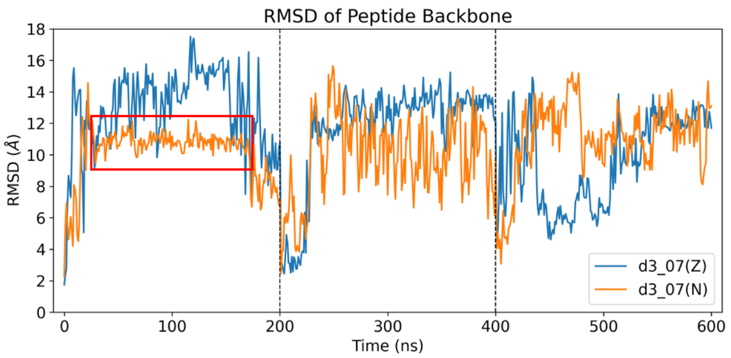

**Figure S23:** RMSD of peptide backbone atoms in the combined 3 × 200 ns MD trajectories (fitted based on PL$^{pro}$ backbone) for peptide **3** initiated from the d3_07 conformation, in the N (orange) and Z (blue) states of Cys111-His272, relative to the starting conformation. The period of stable binding in the N state is indicated by a red box.

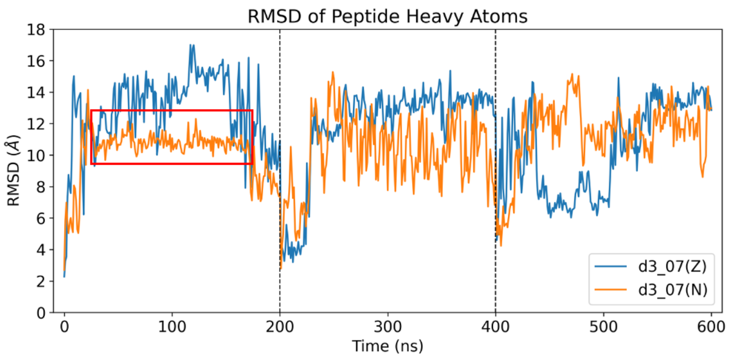

**Figure S24:** RMSD of peptide non-hydrogen atoms in the combined 3 × 200 ns MD trajectories (fitted based on PL$^{pro}$ backbone) for peptide **3** initiated from the d3_07 conformation, in the N (orange) and Z (blue) states of Cys111-His272, relative to the starting conformation. The period of stable binding in the N state is indicated by a red box.

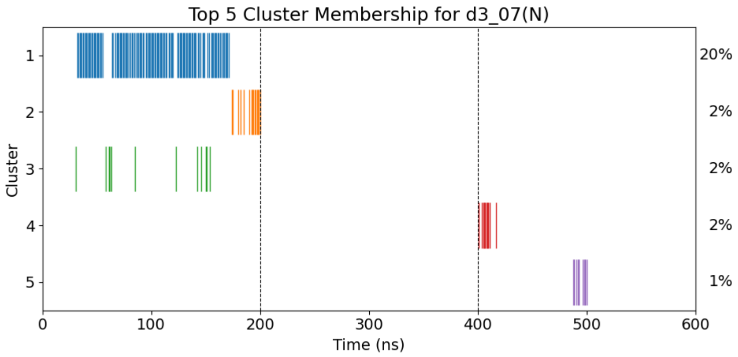

**Figure S25:** Cluster membership of frames extracted every ns from the combined 3 × 200 ns MD (fitted using PL$^{pro}$ backbone) of PL$^{pro}$ complexed with d3_07 (N state), with clustering performed using a 3 Å RMSD cut-off on the peptide non-hydrogen atoms using gmx cluster (gromos algorithm).[5,6] Only the 5 clusters with the highest occupancy percentages are shown. Cluster 1, which has the highest population, corresponds to the period of stable binding shown in **Figures S23-S24**. The complex structure at *t* = 159 ns is determined to be the representative middle structure for cluster 1, as defined by the smallest average RMSD to all other structures in the same cluster. Out of 601 frames, the number of (multimembered) clusters obtained is 344 (63).

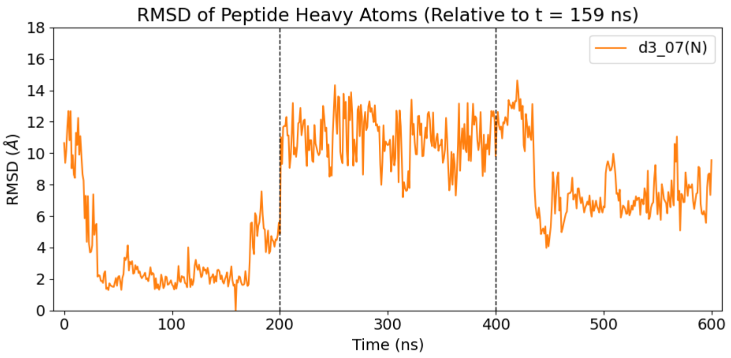

**Figure S26:** RMSD of peptide non-hydrogen atoms in the combined 3 × 200 ns MD trajectories (fitted based on PL$^{pro}$ backbone) for peptide **3** initiated from the d3_07 conformation in the N state, relative to the *t* = 159 ns frame which is representative of cluster 1 (see **Figure S25**).

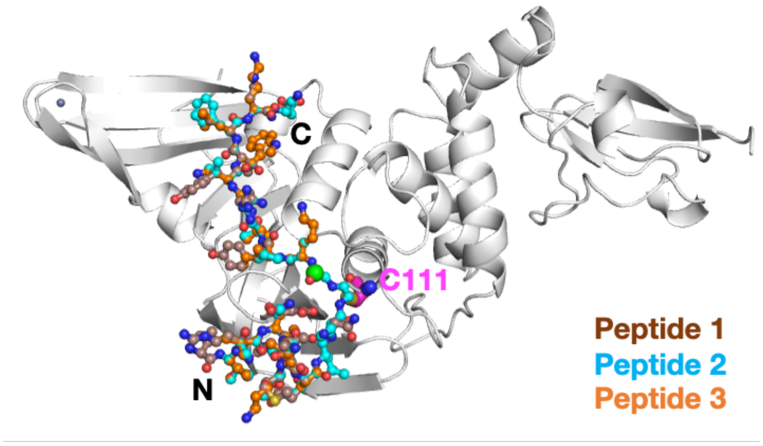

**Figure S27:** Starting conformations of PL$^{pro}$ complexed with the three nsp oligopeptides **1**, **2**, and **3** in the reverse active site binding mode.

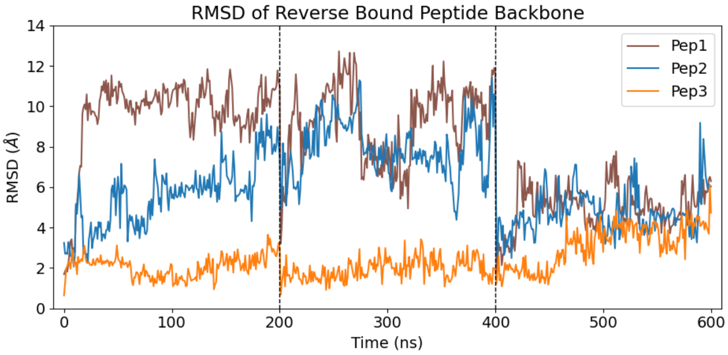

**Figure S28:** RMSD of peptide backbone atoms in the combined 3 × 200 ns MD trajectories (fitted based on PL$^{pro}$ backbone) for the three reverse-bound nsp peptides, in the Z state of Cys111-His272, relative to their corresponding starting conformations.

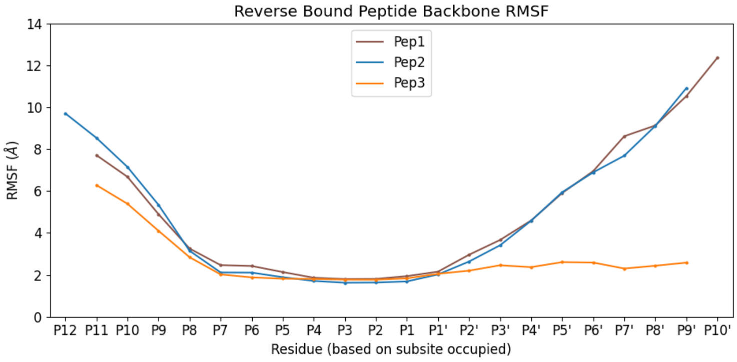

**Figure S29:** Per-residue RMSF of peptide backbone atoms in the combined 3 × 200 ns MD (fitted based on PL$^{pro}$ backbone) for the three reverse-bound nsp peptides, in the Z state of Cys111-His272. Note that the C-terminal $NH_2$ cap is treated as a separate residue (P12 for peptide **2**; P11 for peptides **1** and **3**).

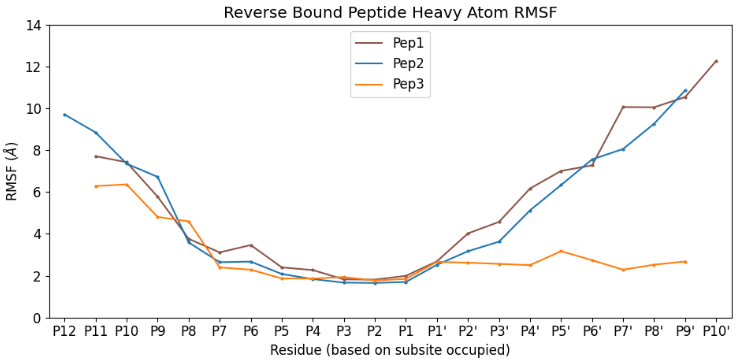

**Figure S30:** Per-residue RMSF of peptide non-hydrogen atoms in the combined 3 × 200 ns MD (fitted based on PL$^{pro}$ backbone) for the three reverse-bound nsp peptides, in the Z state of Cys111-His272.

**Table S3:** PL$^{pro}$-peptide HBs of significance (occurrence ≥ 25% out of 600 frames analyzed every ns) observed over the combined 3 × 200 ns MD for each of the three reverse-bound nsp peptides, in the Z state of Cys111-His272. HB occurrences involving equivalent carboxylate oxygens (in Asp and Glu sidechains) are combined, while salt bridges involving carboxylate oxygens and Arg sidechain guanidino group are combined and halved to avoid double counting. Peptide residues are italicized, while PL$^{pro}$ residues are not. HBs in the S and S′ subsites that are outside the S4-S1 subsites are in blue and red respectively.

| Donor | Acceptor | % | | Donor | Acceptor | % | | Donor | Acceptor | % |
|---|---|---|---|---|---|---|---|---|---|---|
| Pep 1 (Z) | Res 10-13 in S1-S4 | | | Pep 2 (Z) | Res 10-13 in S1-S4 | | | Pep 3 (Z) | Res 10-13 in S1-S4 | |
| *ALA12N* | GLY163O | 94 | | *ALA12N* | GLY163O | 85 | | ASN267ND2 | *VAL1O* | 95 |
| GLY271N | *GLY11O* | 87 | | ARG166NH | *LYS15O* | 69 | | ASN267ND2 | *THR3OG1* | 94 |
| TYR112N | *GLY10O* | 58 | | *THR14OG1* | ASP164OD | 68 | | *LYS12N* | GLY163O | 93 |
| *GLY11N* | GLY271O | 52 | | TYR112N | *GLY10O* | 51 | | GLY271N | *GLY11O* | 91 |
| *ARG6NH* | ASP286OD | 42 | | *LYS15NZ* | GLU167OE | 51 | | *THR3OG1* | GLN269OE1 | 81 |
| *TYR13N* | GLN269O | 36 | | *GLY10N* | GLY271O | 35 | | *ILE13N* | GLN269O | 71 |
| *ARG15NH* | GLU167OE | 35 | | ARG166NE | *THR14OG1* | 34 | | TYR112N | *GLY10O* | 66 |
| *THR14OG1* | ASP164OD | 32 | | TYR273OH | *ALA12O* | 33 | | TYR273OH | *LYS12O* | 48 |
| TYR273OH | *ALA12O* | 32 | | GLY271N | *GLY11O* | 33 | | ARG166NH | *ASN15O* | 46 |
| HIP272ND1 | *ASN9O* | 30 | | *GLY11N* | GLY271O | 33 | | *LYS12NZ* | ASP164OD | 44 |
| ARG166NE | *THR14OG1* | 26 | | ARG166NH | *ASP20OD* | 30 | | HIP272ND1 | *LYS9O* | 34 |
| | | | | | | | | ARG166NE | *ASN15O* | 33 |
| | | | | | | | | *GLY11N* | GLY271O | 30 |
| | | | | | | | | *LYS19NZ* | GLU203OE | 30 |
| | | | | | | | | *LYS12NZ* | GLU167OE | 29 |
| | | | | | | | | ASN109ND2 | *LYS9O* | 26 |

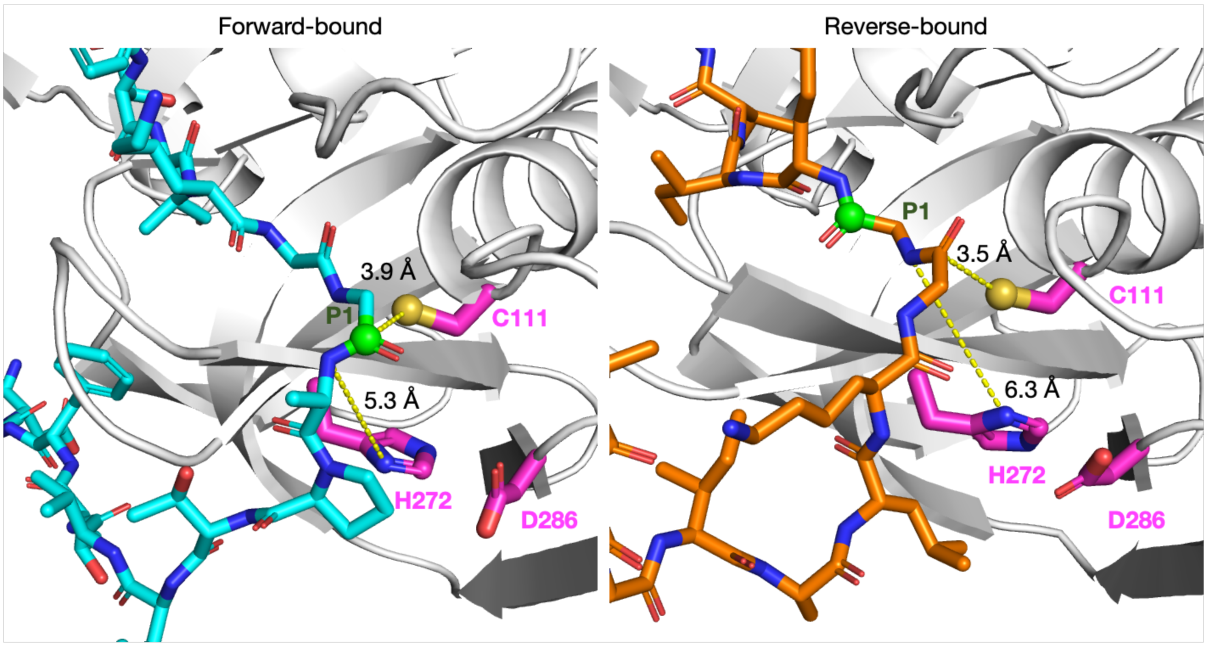

**Figure S31:** Comparison of the modeled forward and reverse oligopeptide binding modes near the active site, using the structures of PL$^{pro}$ complexed with forward-bound peptide **2** (cyan) and reverse-bound peptide **3** (orange) prior to MD for illustration. The catalytic triad residue sidechains are shown as magenta sticks and are labeled. The original P1 residue backbone carbonyl carbon is shown as a green sphere. Note that in the reverse binding mode, while Cys111 is positioned to attack the P2 carbonyl carbon, in the subsequent step the substrate amide nitrogen atom would be too far from His272 to be protonated and act as the leaving group. Relevant calculated distances (yellow dashed lines) are given.

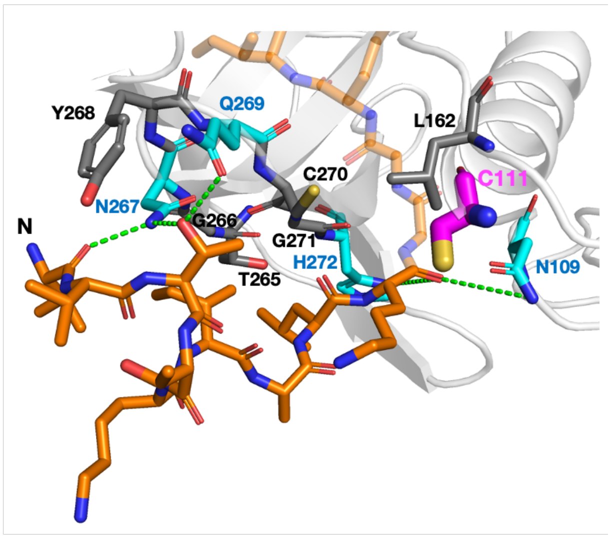

**Figure S32:** View of clustering-derived representative structure of Z-state PL$^{pro}$ (white cartoon) complexed with reverse-bound peptide **3** (orange), with focus on the interactions in the S′ subsites. PL$^{pro}$ residues involved in HBs (green dotted lines; see **Table S3**; not all significant HBs are present in these snapshots) are shown as cyan sticks. Other residues in close contact with the peptide N-terminal residues (within 4 Å) are shown in grey. The active site Cys111 is shown as magenta sticks. Clustering was performed using a 3 Å RMSD cut-off on the peptide non-hydrogen atoms using gmx cluster (gromos algorithm).[5,6] Out of 601 frames extracted every ns from the combined 3 × 200 ns MD, the number of (multimembered) clusters obtained is 29 (21).

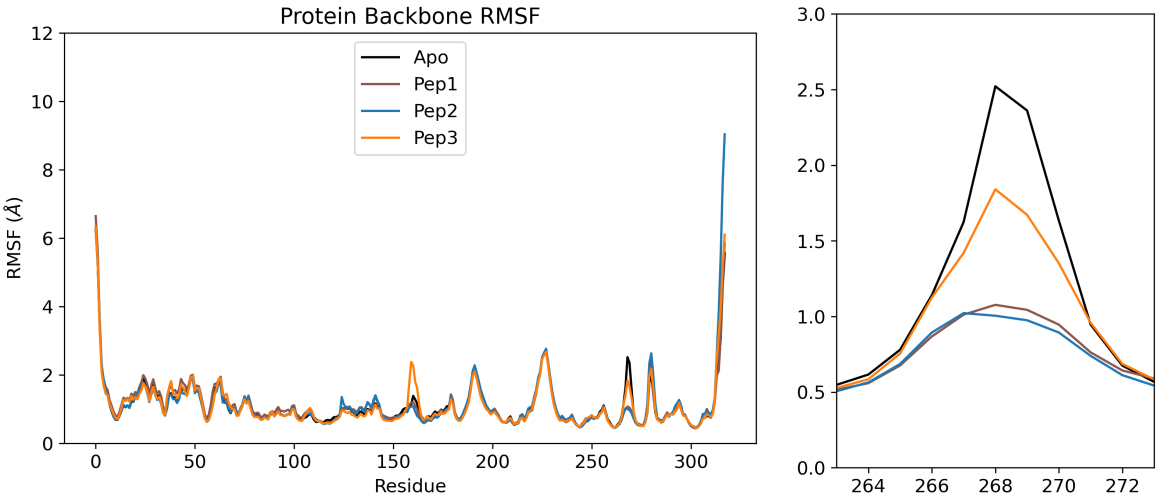

**Figure S33:** Per-residue RMSF of PL$^{pro}$ backbone atoms in the combined 3 × 200 ns MD (fitted based on PL$^{pro}$ backbone) for PL$^{pro}$ (Z state) when apo and when complexed with each of the three forward-bound nsp peptides, (left) in full and (right) focusing around the Tyr268-containing loop region.

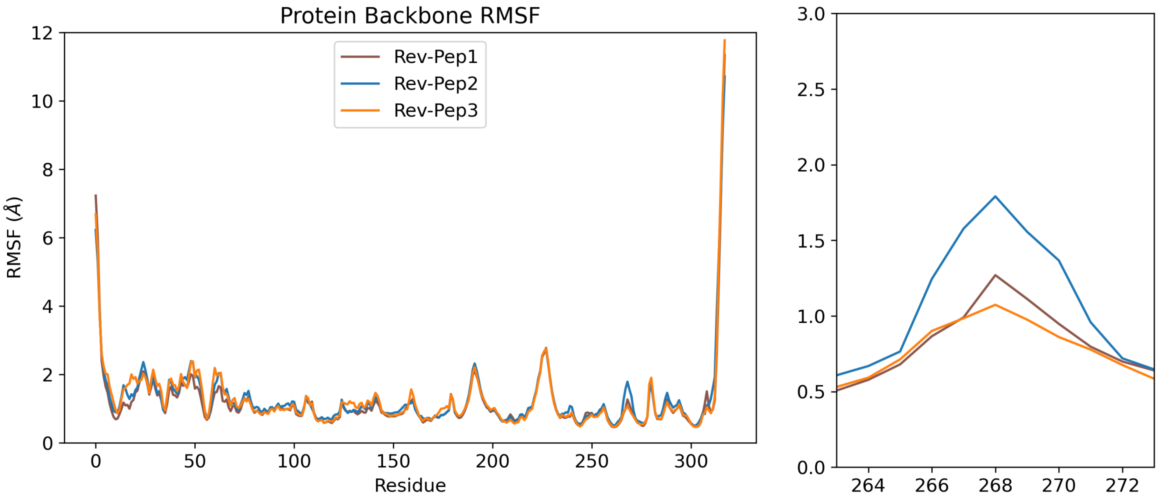

**Figure S34:** Per-residue RMSF of PL$^{pro}$ backbone atoms in the combined 3 × 200 ns MD (fitted based on PL$^{pro}$ backbone) for PL$^{pro}$ (Z state) when complexed with each of the three reverse-bound nsp peptides, (left) in full and (right) focusing around the Tyr268-containing loop region.

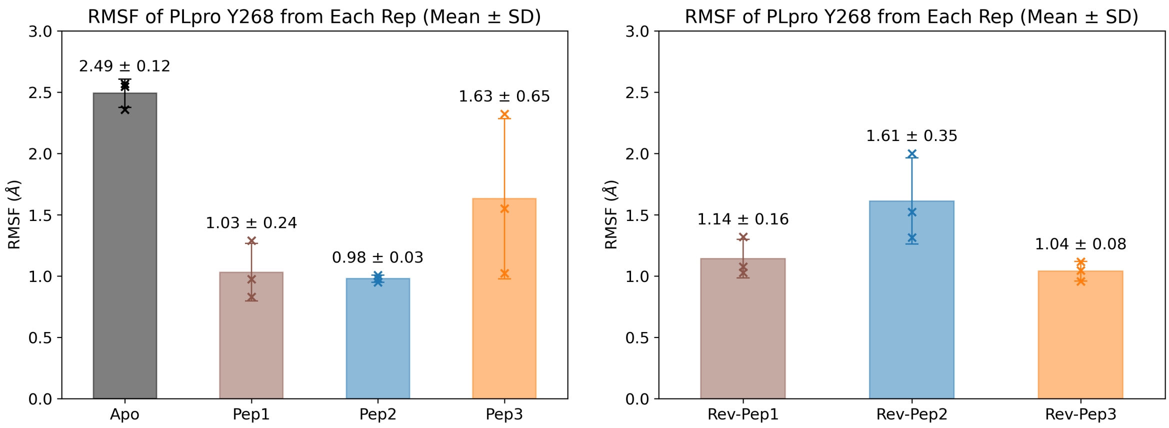

**Figure S35:** Per-residue RMSF of PL$^{pro}$ Tyr268 backbone atoms in each replica of 200 ns MD (fitted based on PL$^{pro}$ backbone) for PL$^{pro}$ (Z state) (left) when apo and when complexed with forward-bound peptides, and (right) when complexed with reverse-bound peptides. The bar height is the arithmetic mean RMSF, and the error bar corresponds to the standard deviation (SD) across the three replicas.

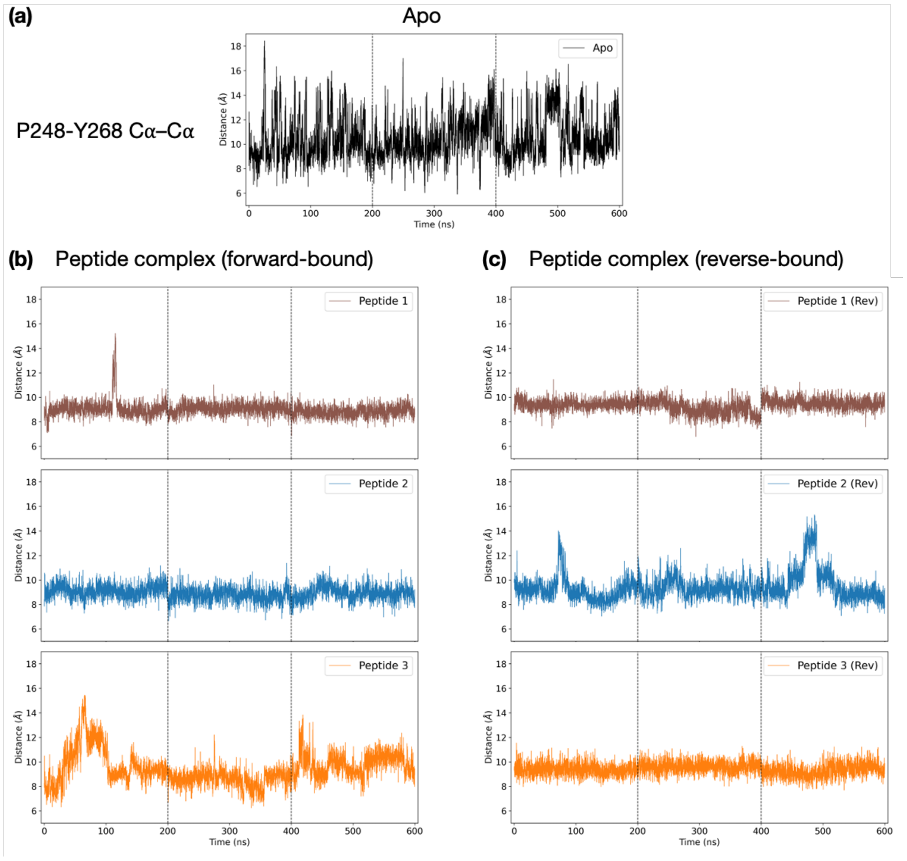

**Figure S36:** The evolution of the Pro248–Tyr268 Cα–Cα distance over combined 3 × 200 ns MD for PL$^{pro}$ (a) in the apo state, and when complexed with the nsp peptides in the (b) forward and (c) reverse modes.

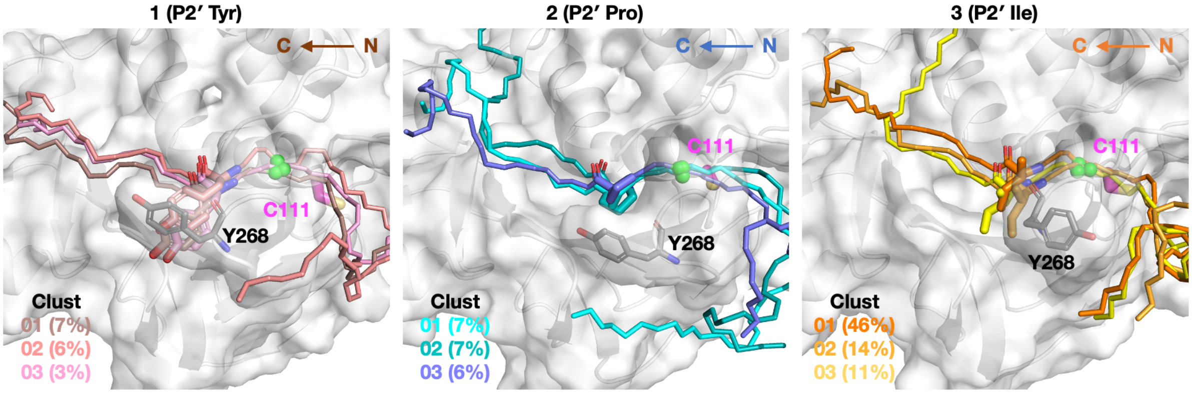

**Figure S37:** View of representative structures of the three most populated clusters of Z-state PL$^{pro}$ (white cartoon and surface) complexed with reverse-bound peptides **1**, **2**, and **3**, with a focus on the S4-occupying residue (which is originally the P2′ residue). For simplification, for the rest of the peptides only the backbone atoms are shown in a uniform color. Tyr268 is shown as grey sticks, Cys111 as magenta sticks, and the original P1-Gly amide carbonyl carbon atoms as green spheres. Clustering was performed using a 3 Å RMSD cut-off on the peptide non-hydrogen atoms using gmx cluster (gromos algorithm).[5,6] Out of 601 frames extracted every ns from the combined 3 × 200 ns MD, the number of (multimembered) clusters obtained for peptides **1**, **2**, and **3** are, respectively, 226 (83), 213 (68), and 29 (21).

**Table S4:** Deviations of P4-P1 Cα atom positions (compared to the VIR251 in PDB 6WX4)[3] and RMSD values for the 100 highest ranked solutions from ADCP[1] docking of peptides **8**, **9**, and **10**, derived from the human proteins IRF3, PROS1, and ULK1, respectively. Solutions that pass the filter are in green. Other solutions which have Gly-Gly in S2-S1 are in orange.

| *Peptide 8* | IRF3 | | | | | *Peptide 9* | PROS1 | | | | | *Peptide 10* | ULK1 | | | | |
|---|---|---|---|---|---|---|---|---|---|---|---|---|---|---|---|---|---|
| **Ranking** | **Deviation (Å)** | | | | **RMSD (Å)** | **Ranking** | **Deviation (Å)** | | | | **RMSD (Å)** | **Ranking** | **Deviation (Å)** | | | | **RMSD (Å)** |
| | *P4* | *P3* | *P2* | *P1* | | | *P4* | *P3* | *P2* | *P1* | | | *P4* | *P3* | *P2* | *P1* | |
| 1 | 20.691 | 14.988 | 11.329 | 12.316 | 15.271 | 1 | 21.076 | 19.259 | 17.103 | 16.880 | 18.658 | 1 | 12.663 | 7.514 | 3.548 | 4.217 | 7.861 |
| 2 | 26.166 | 27.233 | 23.456 | 21.644 | 24.723 | 2 | 18.251 | 12.861 | 12.110 | 10.890 | 13.818 | 2 | 26.802 | 29.800 | 29.013 | 26.499 | 28.064 |
| 3 | 30.575 | 32.051 | 27.490 | 22.512 | 28.393 | 3 | 11.785 | 7.754 | 1.270 | 6.462 | 7.784 | 3 | 30.919 | 31.341 | 28.685 | 25.911 | 29.294 |
| 4 | 9.673 | 6.512 | 6.460 | 7.313 | 7.602 | 4 | 31.728 | 28.276 | 23.500 | 19.529 | 26.172 | 4 | 11.646 | 7.326 | 0.722 | 6.814 | 7.685 |
| 5 | 17.590 | 12.066 | 9.581 | 5.071 | 11.964 | 5 | 30.648 | 29.071 | 23.579 | 20.387 | 26.249 | 5 | 25.403 | 27.730 | 22.492 | 22.203 | 24.562 |
| 6 | 20.291 | 16.738 | 14.997 | 15.492 | 17.006 | 6 | 0.981 | 0.741 | 0.702 | 0.694 | 0.788 | 6 | 12.550 | 8.660 | 3.674 | 4.020 | 8.096 |
| 7 | 30.030 | 31.418 | 27.298 | 22.797 | 28.079 | 7 | 18.226 | 17.470 | 17.154 | 17.163 | 17.509 | 7 | 45.458 | 41.300 | 38.057 | 33.415 | 39.803 |
| 8 | 11.887 | 11.666 | 13.353 | 12.782 | 12.441 | 8 | 34.235 | 30.474 | 30.091 | 28.322 | 30.855 | 8 | 11.376 | 6.749 | 0.269 | 7.174 | 7.525 |
| 9 | 10.673 | 10.953 | 9.500 | 11.421 | 10.661 | 9 | 17.485 | 17.538 | 20.884 | 19.271 | 18.847 | 9 | 6.453 | 0.641 | 0.536 | 0.555 | 3.265 |
| 10 | 21.082 | 18.648 | 17.556 | 17.096 | 18.659 | 10 | 28.704 | 27.161 | 24.419 | 23.441 | 26.017 | 10 | 20.371 | 17.614 | 14.836 | 11.094 | 16.343 |
| 11 | 19.702 | 19.001 | 14.174 | 15.563 | 17.265 | 11 | 12.565 | 8.398 | 2.960 | 4.667 | 8.046 | 11 | 11.715 | 6.787 | 0.492 | 6.859 | 7.592 |
| 12 | 11.265 | 7.065 | 0.615 | 6.908 | 7.499 | 12 | 33.369 | 31.689 | 29.804 | 26.067 | 30.354 | 12 | 12.975 | 8.051 | 3.630 | 4.065 | 8.107 |
| 13 | 12.320 | 11.779 | 12.456 | 12.425 | 12.248 | 13 | 14.189 | 15.573 | 11.775 | 10.268 | 13.114 | 13 | 27.052 | 26.956 | 24.305 | 20.514 | 24.850 |
| 14 | 18.997 | 17.115 | 15.134 | 10.545 | 15.764 | 14 | 17.633 | 15.702 | 13.248 | 9.508 | 14.347 | 14 | 21.116 | 18.839 | 13.351 | 12.353 | 16.820 |
| 15 | 17.135 | 17.100 | 15.180 | 10.824 | 15.277 | 15 | 21.699 | 15.364 | 17.451 | 17.363 | 18.117 | 15 | 31.001 | 26.026 | 24.437 | 19.253 | 25.526 |
| 16 | 19.973 | 16.753 | 15.648 | 16.511 | 17.299 | 16 | 37.978 | 32.768 | 31.810 | 27.811 | 32.792 | 16 | 30.274 | 25.702 | 24.355 | 19.348 | 25.222 |
| 17 | 13.107 | 11.471 | 10.480 | 12.410 | 11.908 | 17 | 15.371 | 12.243 | 14.213 | 10.322 | 13.179 | 17 | 20.039 | 19.637 | 14.544 | 10.271 | 16.615 |
| 18 | 22.687 | 19.700 | 15.029 | 10.059 | 17.535 | 18 | 21.918 | 19.298 | 16.352 | 13.399 | 18.026 | 18 | 5.990 | 0.314 | 0.204 | 0.326 | 3.006 |
| 19 | 20.821 | 18.225 | 16.819 | 16.613 | 18.197 | 19 | 29.433 | 29.209 | 23.557 | 22.084 | 26.278 | 19 | 31.486 | 25.165 | 24.560 | 19.813 | 25.595 |
| 20 | 17.576 | 11.120 | 9.732 | 5.296 | 11.783 | 20 | 40.358 | 35.592 | 33.287 | 26.988 | 34.395 | 20 | 19.005 | 18.689 | 15.009 | 16.053 | 17.273 |
| 21 | 12.096 | 11.960 | 12.863 | 12.971 | 12.481 | 21 | 1.135 | 0.429 | 0.547 | 0.825 | 0.783 | 21 | 14.141 | 13.780 | 13.020 | 13.568 | 13.633 |
| 22 | 19.244 | 15.705 | 10.521 | 12.630 | 14.893 | 22 | 14.277 | 9.035 | 3.562 | 4.178 | 8.883 | 22 | 12.411 | 7.737 | 3.733 | 4.061 | 7.815 |
| 23 | 17.168 | 15.298 | 17.201 | 15.271 | 16.262 | 23 | 39.317 | 34.383 | 32.064 | 28.067 | 33.705 | 23 | 6.370 | 0.480 | 0.407 | 0.334 | 3.205 |
| 24 | 14.089 | 13.870 | 12.312 | 14.472 | 13.710 | 24 | 30.028 | 23.453 | 20.499 | 12.864 | 22.569 | 24 | 13.952 | 8.293 | 3.821 | 3.846 | 8.556 |
| 25 | 19.352 | 17.359 | 14.816 | 12.363 | 16.188 | 25 | 12.449 | 8.372 | 3.093 | 4.471 | 7.979 | 25 | 20.832 | 18.536 | 14.731 | 11.662 | 16.812 |
| 26 | 12.480 | 11.524 | 11.101 | 13.938 | 12.309 | 26 | 21.488 | 18.817 | 14.730 | 9.978 | 16.825 | 26 | 30.233 | 27.101 | 24.454 | 19.728 | 25.669 |
| 27 | 17.630 | 11.413 | 9.271 | 5.143 | 11.763 | 27 | 22.502 | 20.023 | 16.919 | 15.477 | 18.928 | 27 | 22.096 | 19.487 | 14.444 | 15.792 | 18.207 |
| 28 | 20.641 | 22.132 | 26.152 | 28.800 | 24.644 | 28 | 26.184 | 25.970 | 29.012 | 29.035 | 27.590 | 28 | 13.144 | 15.459 | 15.164 | 15.139 | 14.755 |
| 29 | 25.775 | 25.880 | 30.175 | 32.643 | 28.767 | 29 | 30.986 | 26.733 | 23.596 | 20.537 | 25.755 | 29 | 38.179 | 32.881 | 30.668 | 25.908 | 32.213 |
| 30 | 20.205 | 16.639 | 14.992 | 15.520 | 16.961 | 30 | 31.806 | 28.913 | 25.012 | 19.249 | 26.663 | 30 | 30.727 | 29.031 | 30.082 | 30.484 | 30.088 |
| 31 | 24.233 | 22.299 | 26.627 | 28.817 | 25.612 | 31 | 21.046 | 17.326 | 12.489 | 8.450 | 15.576 | 31 | 32.089 | 27.556 | 27.318 | 22.498 | 27.575 |
| 32 | 23.044 | 18.123 | 16.737 | 12.799 | 18.052 | 32 | 19.615 | 14.246 | 12.180 | 9.108 | 14.309 | 32 | 24.393 | 19.035 | 18.574 | 13.644 | 19.291 |
| 33 | 19.841 | 18.325 | 14.756 | 15.791 | 17.296 | 33 | 19.701 | 14.034 | 13.578 | 8.600 | 14.521 | 33 | 20.053 | 16.283 | 15.602 | 10.838 | 16.033 |
| 34 | 29.485 | 27.341 | 20.815 | 19.940 | 24.737 | 34 | 20.643 | 18.835 | 16.820 | 16.580 | 18.294 | 34 | 31.005 | 29.692 | 30.207 | 30.504 | 30.356 |
| 35 | 22.886 | 19.103 | 18.205 | 18.458 | 19.753 | 35 | 17.205 | 14.089 | 13.719 | 10.832 | 14.142 | 35 | 38.870 | 34.856 | 33.079 | 27.674 | 33.859 |
| 36 | 26.523 | 27.270 | 24.076 | 21.748 | 24.999 | 36 | 13.374 | 8.961 | 3.363 | 4.314 | 8.501 | 36 | 14.647 | 9.749 | 11.869 | 13.805 | 12.659 |
| 37 | 26.606 | 22.046 | 17.865 | 14.534 | 20.762 | 37 | 22.831 | 16.179 | 15.177 | 9.894 | 16.668 | 37 | 19.449 | 18.656 | 13.305 | 9.808 | 15.808 |
| 38 | 7.044 | 6.165 | 9.760 | 10.641 | 8.604 | 38 | 17.916 | 15.767 | 11.780 | 8.513 | 13.972 | 38 | 13.992 | 13.712 | 15.522 | 16.683 | 15.025 |
| 39 | 23.949 | 20.824 | 17.191 | 15.842 | 19.708 | 39 | 14.400 | 15.122 | 15.782 | 15.866 | 15.304 | 39 | 38.435 | 36.025 | 32.879 | 26.802 | 33.817 |
| 40 | 14.874 | 12.558 | 13.756 | 16.505 | 14.496 | 40 | 16.689 | 17.024 | 17.090 | 17.777 | 17.150 | 40 | 40.739 | 41.688 | 39.097 | 34.607 | 39.127 |
| 41 | 19.069 | 18.892 | 14.883 | 11.338 | 16.360 | 41 | 19.004 | 16.033 | 14.334 | 10.930 | 15.355 | 41 | 15.809 | 10.564 | 11.097 | 9.719 | 12.032 |
| 42 | 12.268 | 6.742 | 0.644 | 6.856 | 7.800 | 42 | 40.782 | 35.525 | 32.894 | 28.495 | 34.710 | 42 | 39.401 | 35.295 | 30.683 | 26.048 | 33.234 |
| 43 | 13.340 | 16.318 | 17.258 | 21.746 | 17.428 | 43 | 22.383 | 20.613 | 16.273 | 13.634 | 18.551 | 43 | 28.203 | 25.487 | 24.805 | 25.407 | 26.009 |
| 44 | 29.522 | 26.607 | 20.674 | 20.107 | 24.552 | 44 | 17.027 | 12.239 | 9.223 | 7.938 | 12.122 | 44 | 5.488 | 0.778 | 0.376 | 0.405 | 2.785 |
| 45 | 11.696 | 11.449 | 13.140 | 12.376 | 12.183 | 45 | 17.695 | 13.742 | 11.542 | 11.610 | 13.874 | 45 | 30.508 | 29.834 | 29.577 | 29.551 | 29.870 |
| 46 | 30.424 | 27.337 | 24.964 | 17.480 | 25.503 | 46 | 26.188 | 21.323 | 22.136 | 21.173 | 22.797 | 46 | 30.065 | 24.869 | 23.670 | 18.878 | 24.693 |
| 47 | 18.992 | 19.500 | 15.180 | 16.004 | 17.518 | 47 | 16.113 | 12.168 | 14.415 | 10.869 | 13.543 | 47 | 12.991 | 17.066 | 22.337 | 28.731 | 21.121 |
| 48 | 19.040 | 17.955 | 14.451 | 12.287 | 16.161 | 48 | 27.159 | 26.221 | 24.256 | 20.438 | 24.654 | 48 | 20.242 | 21.853 | 23.716 | 24.397 | 22.610 |
| 49 | 12.246 | 6.726 | 0.692 | 6.839 | 7.785 | 49 | 30.480 | 30.550 | 25.052 | 24.015 | 27.689 | 49 | 21.073 | 18.474 | 15.922 | 11.910 | 17.181 |
| 50 | 21.462 | 17.141 | 13.699 | 13.579 | 16.782 | 50 | 14.814 | 12.838 | 14.369 | 12.699 | 13.711 | 50 | 17.967 | 13.439 | 11.692 | 7.416 | 13.182 |
| 51 | 27.908 | 28.925 | 25.734 | 24.096 | 26.732 | 51 | 27.023 | 26.952 | 24.537 | 20.713 | 24.939 | 51 | 37.171 | 32.765 | 31.211 | 26.662 | 32.172 |
| 52 | 25.663 | 22.495 | 24.807 | 24.688 | 24.441 | 52 | 39.351 | 32.574 | 30.144 | 27.115 | 32.609 | 52 | 12.889 | 8.350 | 3.605 | 4.194 | 8.161 |
| 53 | 29.671 | 27.910 | 23.621 | 18.261 | 25.252 | 53 | 8.913 | 2.015 | 7.871 | 13.392 | 9.011 | 53 | 18.799 | 16.341 | 20.513 | 20.978 | 19.244 |
| 54 | 18.734 | 17.404 | 20.405 | 21.843 | 19.668 | 54 | 27.557 | 26.375 | 24.011 | 19.885 | 24.632 | 54 | 35.488 | 28.976 | 24.772 | 20.153 | 27.923 |
| 55 | 18.370 | 12.512 | 9.910 | 5.155 | 12.438 | 55 | 26.560 | 26.679 | 29.708 | 30.054 | 28.298 | 55 | 13.577 | 12.056 | 11.402 | 13.414 | 12.645 |
| 56 | 27.075 | 26.458 | 23.556 | 19.451 | 24.322 | 56 | 19.402 | 13.226 | 14.128 | 9.822 | 14.555 | 56 | 27.469 | 29.756 | 26.799 | 25.072 | 27.326 |
| 57 | 21.372 | 18.220 | 13.268 | 14.190 | 17.074 | 57 | 37.838 | 32.868 | 33.611 | 29.328 | 33.548 | 57 | 7.145 | 0.376 | 0.309 | 0.420 | 3.587 |
| 58 | 27.416 | 22.950 | 21.197 | 16.348 | 22.332 | 58 | 0.814 | 0.227 | 0.354 | 0.693 | 0.575 | 58 | 28.243 | 24.220 | 24.823 | 20.538 | 24.608 |
| 59 | 11.046 | 7.784 | 7.255 | 9.063 | 8.908 | 59 | 13.419 | 13.331 | 13.502 | 13.993 | 13.564 | 59 | 18.359 | 14.162 | 16.956 | 17.214 | 16.744 |
| 60 | 18.556 | 11.710 | 9.933 | 5.081 | 12.308 | 60 | 15.034 | 10.638 | 10.345 | 8.994 | 11.479 | 60 | 24.948 | 21.134 | 20.163 | 16.328 | 20.870 |
| 61 | 30.378 | 32.123 | 27.672 | 23.238 | 28.550 | 61 | 28.684 | 26.185 | 22.582 | 22.923 | 25.218 | 61 | 13.157 | 10.947 | 12.003 | 11.043 | 11.821 |
| 62 | 29.465 | 25.386 | 20.854 | 13.210 | 23.033 | 62 | 32.653 | 29.050 | 23.693 | 18.989 | 26.609 | 62 | 15.153 | 11.757 | 10.722 | 12.871 | 12.732 |
| 63 | 18.813 | 14.783 | 14.520 | 15.039 | 15.886 | 63 | 21.482 | 18.976 | 17.415 | 17.097 | 18.822 | 63 | 30.944 | 30.191 | 27.836 | 29.789 | 29.712 |
| 64 | 19.235 | 17.356 | 20.577 | 20.606 | 19.489 | 64 | 21.759 | 17.088 | 13.132 | 13.918 | 16.820 | 64 | 31.708 | 27.685 | 23.867 | 16.933 | 25.633 |
| 65 | 16.665 | 11.720 | 11.730 | 8.710 | 12.535 | 65 | 22.324 | 22.765 | 18.469 | 14.002 | 19.709 | 65 | 12.798 | 13.036 | 11.485 | 9.454 | 11.779 |
| 66 | 28.216 | 26.191 | 20.952 | 18.130 | 23.716 | 66 | 42.391 | 38.717 | 37.043 | 33.314 | 38.007 | 66 | 33.496 | 35.322 | 33.082 | 30.927 | 33.243 |
| 67 | 10.723 | 7.143 | 7.167 | 7.800 | 8.340 | 67 | 18.926 | 21.189 | 15.219 | 12.485 | 17.282 | 67 | 44.035 | 43.143 | 39.384 | 34.631 | 40.469 |
| 68 | 13.189 | 7.837 | 8.228 | 9.332 | 9.876 | 68 | 12.576 | 14.023 | 15.133 | 14.911 | 14.196 | 68 | 6.268 | 0.348 | 0.677 | 1.028 | 3.199 |
| 69 | 29.453 | 27.523 | 29.247 | 28.515 | 28.695 | 69 | 32.491 | 29.851 | 25.878 | 19.487 | 27.369 | 69 | 45.462 | 41.439 | 37.403 | 33.058 | 39.610 |
| 70 | 21.829 | 21.695 | 23.691 | 26.053 | 23.384 | 70 | 32.214 | 27.082 | 26.563 | 22.656 | 27.341 | 70 | 24.552 | 27.003 | 22.592 | 21.379 | 23.976 |
| 71 | 12.551 | 12.164 | 13.986 | 12.562 | 12.835 | 71 | 11.665 | 8.976 | 12.068 | 12.049 | 11.263 | 71 | 26.778 | 25.074 | 24.809 | 21.665 | 24.651 |
| 72 | 19.477 | 16.851 | 20.195 | 21.865 | 19.680 | 72 | 21.107 | 19.615 | 18.674 | 13.991 | 18.539 | 72 | 5.391 | 0.830 | 0.368 | 0.686 | 2.755 |
| 73 | 17.329 | 12.440 | 10.789 | 10.510 | 13.056 | 73 | 30.662 | 28.185 | 29.179 | 27.507 | 28.907 | 73 | 30.960 | 29.574 | 30.164 | 30.027 | 30.185 |
| 74 | 16.904 | 16.583 | 15.971 | 16.218 | 16.423 | 74 | 31.275 | 29.465 | 26.133 | 20.521 | 27.159 | 74 | 30.289 | 29.590 | 29.719 | 28.374 | 29.501 |
| 75 | 34.709 | 36.127 | 34.818 | 33.596 | 34.824 | 75 | 21.169 | 19.693 | 24.833 | 22.877 | 22.226 | 75 | 12.836 | 11.994 | 14.555 | 14.711 | 13.572 |
| 76 | 33.779 | 36.609 | 35.487 | 30.978 | 34.279 | 76 | 31.331 | 28.389 | 23.830 | 19.075 | 26.074 | 76 | 28.680 | 28.638 | 22.228 | 19.308 | 25.048 |
| 77 | 15.351 | 11.047 | 10.719 | 10.344 | 12.037 | 77 | 21.110 | 14.497 | 14.503 | 10.332 | 15.596 | 77 | 11.266 | 6.657 | 0.383 | 7.314 | 7.498 |
| 78 | 11.793 | 10.642 | 10.062 | 13.661 | 11.621 | 78 | 30.036 | 25.670 | 21.575 | 21.574 | 24.960 | 78 | 25.757 | 21.168 | 18.450 | 14.354 | 20.359 |
| 79 | 26.638 | 28.374 | 22.533 | 21.123 | 24.842 | 79 | 36.648 | 35.685 | 33.309 | 28.223 | 33.625 | 79 | 12.338 | 8.272 | 3.351 | 4.398 | 7.925 |
| 80 | 10.620 | 10.878 | 10.094 | 11.854 | 10.880 | 80 | 13.411 | 8.909 | 3.122 | 4.651 | 8.524 | 80 | 20.202 | 16.316 | 15.598 | 11.222 | 16.152 |
| 81 | 13.312 | 13.470 | 10.736 | 14.199 | 12.995 | 81 | 28.468 | 26.260 | 23.207 | 22.892 | 25.311 | 81 | 30.268 | 30.852 | 29.007 | 24.920 | 28.855 |

| | | | | | | | | | | | | | | | | | | | |
|---|---|---|---|---|---|---|---|---|---|---|---|---|---|---|---|---|---|---|---|
| 82 | 34.840 | 34.727 | 30.609 | 29.112 | 32.420 | | 82 | 38.657 | 33.586 | 33.706 | 29.521 | 34.022 | | 82 | 63.529 | 57.479 | 53.332 | 47.791 | 55.830 |
| 83 | 23.462 | 21.572 | 15.908 | 12.209 | 18.828 | | 83 | 27.877 | 26.274 | 24.054 | 19.675 | 24.663 | | 83 | 31.400 | 26.767 | 26.052 | 21.216 | 26.605 |
| 84 | 19.064 | 14.861 | 14.321 | 14.746 | 15.865 | | 84 | 16.768 | 17.975 | 17.085 | 17.719 | 17.393 | | 84 | 4.044 | 3.996 | 10.493 | 16.966 | 10.372 |
| 85 | 28.433 | 27.112 | 20.999 | 19.472 | 24.308 | | 85 | 14.685 | 12.910 | 14.588 | 12.465 | 13.698 | | 85 | 14.347 | 11.485 | 11.522 | 12.470 | 12.510 |
| 86 | 11.606 | 7.171 | 0.733 | 6.645 | 7.597 | | 86 | 30.389 | 26.999 | 25.251 | 22.592 | 26.459 | | 86 | 5.613 | 0.951 | 0.891 | 0.924 | 2.918 |
| 87 | 15.770 | 10.738 | 10.010 | 7.412 | 11.392 | | 87 | 18.461 | 15.648 | 13.876 | 15.514 | 15.960 | | 87 | 22.803 | 20.597 | 14.633 | 12.274 | 18.090 |
| 88 | 12.523 | 13.524 | 11.707 | 15.679 | 13.441 | | 88 | 14.463 | 16.910 | 13.376 | 13.006 | 14.519 | | 88 | 14.643 | 12.631 | 12.550 | 10.327 | 12.630 |
| 89 | 45.239 | 45.557 | 48.074 | 46.112 | 46.259 | | 89 | 38.390 | 32.932 | 31.981 | 27.958 | 33.026 | | 89 | 24.102 | 19.276 | 18.604 | 14.628 | 19.446 |
| 90 | 22.863 | 18.192 | 16.886 | 12.580 | 18.007 | | 90 | 23.658 | 22.885 | 21.198 | 19.149 | 21.791 | | 90 | 33.137 | 27.723 | 25.252 | 21.145 | 27.163 |
| 91 | 20.126 | 16.732 | 17.682 | 14.132 | 17.301 | | 91 | 36.404 | 30.811 | 28.744 | 24.192 | 30.356 | | 91 | 22.105 | 23.275 | 21.697 | 19.275 | 21.637 |
| 92 | 24.487 | 25.176 | 27.819 | 30.362 | 27.061 | | 92 | 12.172 | 7.650 | 1.364 | 6.525 | 7.923 | | 92 | 11.144 | 6.626 | 0.681 | 7.370 | 7.465 |
| 93 | 17.701 | 11.093 | 9.375 | 5.372 | 11.760 | | 93 | 18.423 | 13.003 | 12.023 | 10.899 | 13.891 | | 93 | 13.714 | 12.225 | 12.306 | 8.841 | 11.907 |
| 94 | 30.556 | 29.966 | 32.072 | 32.570 | 31.309 | | 94 | 29.586 | 29.605 | 31.566 | 31.064 | 30.468 | | 94 | 11.985 | 11.423 | 11.155 | 10.385 | 11.252 |
| 95 | 20.159 | 20.271 | 15.490 | 16.705 | 18.278 | | 95 | 24.271 | 24.022 | 23.634 | 24.265 | 24.049 | | 95 | 16.431 | 14.010 | 17.500 | 18.366 | 16.657 |
| 96 | 17.734 | 15.158 | 17.133 | 15.121 | 16.328 | | 96 | 17.338 | 10.679 | 9.691 | 8.998 | 12.140 | | 96 | 11.805 | 7.187 | 0.603 | 6.736 | 7.693 |
| 97 | 27.092 | 24.839 | 22.227 | 20.378 | 23.771 | | 97 | 1.332 | 0.508 | 0.178 | 0.382 | 0.743 | | 97 | 19.636 | 12.804 | 11.247 | 7.828 | 13.577 |
| 98 | 19.454 | 20.097 | 18.269 | 19.247 | 19.278 | | 98 | 27.880 | 22.269 | 17.417 | 10.810 | 20.576 | | 98 | 32.158 | 30.987 | 31.349 | 31.800 | 31.576 |
| 99 | 32.660 | 31.484 | 32.540 | 31.449 | 32.038 | | 99 | 34.652 | 32.575 | 31.146 | 27.991 | 31.684 | | 99 | 23.225 | 24.207 | 22.031 | 20.372 | 22.504 |
| 100 | 12.424 | 12.057 | 12.723 | 10.933 | 12.054 | | 100 | 40.157 | 33.291 | 28.952 | 26.947 | 32.731 | | 100 | 12.447 | 7.368 | 3.541 | 4.324 | 7.753 |

**Table S5:** Deviations of P4-P1 Cα atom positions (compared to the VIR251 in PDB 6WX4)[3] and RMSD values for the 100 highest ranked solutions from ADCP[1] docking of peptide **11** derived from the human protein ATG7. None of the solutions passed the filter or had a Gly-Gly in S2-S1. Solutions where the peptide passes through S2-S1 are in yellow.

| *Peptide 11* | ATG7 | | | | |
|---|---|---|---|---|---|
| **Ranking** | **Deviation(Å)** | | | | **RMSD(Å)** |
| | *P4* | *P3* | *P2* | *P1* | |
| 1 | 24.146 | 18.466 | 14.408 | 8.272 | 17.321 |
| 2 | 22.661 | 23.382 | 21.177 | 15.339 | 20.880 |
| 3 | 27.039 | 30.261 | 27.297 | 26.712 | 27.863 |
| 4 | 21.213 | 21.798 | 20.288 | 15.273 | 19.812 |
| 5 | 28.858 | 26.516 | 23.809 | 24.188 | 25.922 |
| 6 | 25.972 | 25.956 | 25.364 | 22.815 | 25.060 |
| 7 | 26.087 | 26.573 | 23.289 | 21.794 | 24.515 |
| 8 | 29.679 | 31.882 | 29.182 | 27.231 | 29.540 |
| 9 | 29.074 | 24.228 | 18.254 | 12.004 | 21.850 |
| 10 | 26.997 | 21.082 | 21.437 | 18.325 | 22.185 |
| 11 | 43.782 | 37.931 | 34.472 | 27.697 | 36.439 |
| 12 | 27.916 | 28.063 | 25.421 | 21.954 | 25.957 |
| 13 | 23.682 | 23.583 | 21.601 | 15.514 | 21.356 |
| 14 | 34.681 | 34.114 | 32.082 | 30.547 | 32.897 |
| 15 | 23.121 | 21.828 | 19.609 | 12.732 | 19.734 |
| 16 | 13.513 | 7.986 | 10.109 | 9.339 | 10.438 |
| 17 | 39.861 | 38.628 | 40.778 | 41.826 | 40.291 |
| 18 | 20.335 | 17.590 | 11.596 | 6.524 | 15.000 |
| 19 | 26.799 | 22.570 | 23.739 | 18.063 | 23.008 |
| 20 | 9.246 | 3.158 | 3.166 | 7.696 | 6.417 |
| 21 | 17.448 | 22.418 | 26.469 | 29.720 | 24.448 |
| 22 | 24.168 | 24.618 | 25.501 | 23.925 | 24.560 |
| 23 | 23.340 | 17.549 | 13.661 | 8.147 | 16.626 |
| 24 | 29.742 | 32.834 | 30.845 | 26.788 | 30.132 |
| 25 | 16.359 | 11.543 | 9.929 | 8.919 | 12.031 |
| 26 | 21.760 | 20.369 | 14.298 | 9.299 | 17.170 |
| 27 | 25.744 | 25.200 | 21.251 | 19.254 | 23.022 |
| 28 | 35.985 | 34.227 | 31.426 | 25.205 | 31.974 |
| 29 | 25.468 | 20.450 | 17.549 | 11.722 | 19.444 |
| 30 | 18.149 | 17.654 | 15.596 | 11.627 | 15.965 |
| 31 | 23.469 | 22.271 | 21.732 | 15.577 | 20.987 |
| 32 | 32.757 | 28.374 | 23.792 | 22.195 | 27.096 |
| 33 | 12.383 | 12.076 | 9.240 | 7.307 | 10.464 |
| 34 | 23.294 | 21.501 | 18.033 | 14.224 | 19.573 |
| 35 | 26.815 | 22.678 | 20.954 | 15.843 | 21.929 |
| 36 | 21.106 | 20.956 | 15.086 | 13.041 | 17.904 |
| 37 | 17.799 | 13.363 | 7.549 | 4.815 | 11.995 |
| 38 | 11.963 | 8.214 | 12.195 | 15.410 | 12.214 |
| 39 | 24.486 | 24.894 | 25.494 | 22.736 | 24.424 |
| 40 | 35.218 | 35.235 | 33.692 | 27.884 | 33.145 |
| 41 | 16.111 | 16.971 | 12.972 | 11.562 | 14.573 |
| 42 | 19.689 | 13.857 | 10.849 | 9.782 | 14.080 |
| 43 | 25.753 | 20.952 | 15.913 | 14.413 | 19.769 |
| 44 | 29.451 | 28.255 | 23.957 | 21.827 | 26.058 |
| 45 | 27.441 | 24.773 | 25.094 | 22.394 | 24.990 |
| 46 | 36.855 | 35.511 | 33.371 | 34.126 | 34.991 |
| 47 | 23.834 | 24.400 | 24.154 | 22.173 | 23.656 |
| 48 | 8.088 | 4.003 | 4.028 | 7.231 | 6.123 |
| 49 | 32.119 | 27.936 | 25.673 | 20.879 | 26.959 |
| 50 | 34.957 | 30.435 | 27.744 | 21.984 | 29.161 |
| 51 | 27.168 | 30.062 | 27.429 | 25.690 | 27.632 |
| 52 | 16.750 | 15.689 | 10.854 | 8.123 | 13.328 |
| 53 | 16.082 | 16.522 | 17.695 | 19.788 | 17.580 |
| 54 | 29.480 | 28.185 | 22.300 | 20.323 | 25.366 |
| 55 | 27.406 | 21.161 | 20.718 | 17.509 | 21.992 |
| 56 | 20.743 | 18.213 | 14.725 | 13.384 | 17.014 |
| 57 | 28.452 | 25.228 | 23.794 | 22.963 | 25.196 |
| 58 | 44.517 | 38.651 | 34.920 | 29.808 | 37.362 |
| 59 | 43.926 | 38.541 | 35.424 | 28.655 | 37.050 |
| 60 | 28.481 | 31.029 | 26.524 | 26.383 | 28.167 |
| 61 | 28.568 | 25.866 | 24.765 | 19.437 | 24.881 |
| 62 | 26.914 | 20.908 | 21.625 | 17.729 | 22.042 |
| 63 | 26.867 | 27.681 | 27.292 | 30.578 | 28.142 |
| 64 | 30.946 | 33.258 | 29.226 | 27.193 | 30.238 |
| 65 | 44.062 | 37.789 | 36.976 | 32.794 | 38.119 |
| 66 | 20.063 | 14.382 | 11.671 | 10.054 | 14.549 |
| 67 | 35.626 | 32.205 | 28.949 | 23.406 | 30.382 |
| 68 | 3.947 | 8.819 | 13.124 | 14.870 | 11.031 |
| 69 | 15.958 | 15.415 | 10.769 | 7.879 | 12.945 |
| 70 | 22.895 | 16.141 | 12.849 | 8.445 | 15.978 |
| 71 | 25.323 | 19.264 | 18.567 | 12.420 | 19.438 |
| 72 | 17.197 | 22.859 | 26.480 | 29.292 | 24.380 |
| 73 | 32.720 | 30.270 | 24.796 | 22.179 | 27.810 |
| 74 | 28.921 | 26.225 | 23.383 | 23.665 | 25.646 |
| 75 | 19.772 | 18.516 | 19.601 | 15.553 | 18.438 |
| 76 | 31.739 | 25.947 | 22.864 | 18.456 | 25.219 |
| 77 | 17.466 | 22.216 | 27.582 | 32.940 | 25.712 |
| 78 | 18.950 | 19.698 | 14.874 | 13.083 | 16.878 |
| 79 | 44.167 | 38.874 | 35.751 | 30.657 | 37.683 |
| 80 | 31.887 | 29.713 | 26.523 | 25.003 | 28.409 |
| 81 | 26.758 | 30.275 | 27.049 | 26.545 | 27.699 |

| | | | | | |
|---|---|---|---|---|---|
| 82 | 33.434 | 30.317 | 25.931 | 18.718 | 27.657 |
| 83 | 21.738 | 20.458 | 21.000 | 17.991 | 20.345 |
| 84 | 27.989 | 27.519 | 22.246 | 20.844 | 24.849 |
| 85 | 26.843 | 22.182 | 19.922 | 14.634 | 21.352 |
| 86 | 23.842 | 21.479 | 18.132 | 13.534 | 19.632 |
| 87 | 24.162 | 22.162 | 16.957 | 13.548 | 19.660 |
| 88 | 19.540 | 13.959 | 11.139 | 9.690 | 14.095 |
| 89 | 18.141 | 14.707 | 10.166 | 5.392 | 13.018 |
| 90 | 17.780 | 22.471 | 25.685 | 28.218 | 23.859 |
| 91 | 27.179 | 25.918 | 26.832 | 24.900 | 26.222 |
| 92 | 23.165 | 17.632 | 13.196 | 9.198 | 16.630 |
| 93 | 19.612 | 16.053 | 11.628 | 6.053 | 14.267 |
| 94 | 16.920 | 22.975 | 26.466 | 28.589 | 24.145 |
| 95 | 3.878 | 4.159 | 8.390 | 12.619 | 8.093 |
| 96 | 11.413 | 12.372 | 14.710 | 11.489 | 12.567 |
| 97 | 22.907 | 17.155 | 13.205 | 8.290 | 16.295 |
| 98 | 25.420 | 26.473 | 23.376 | 21.716 | 24.316 |
| 99 | 27.590 | 27.642 | 24.082 | 23.276 | 25.725 |
| 100 | 27.255 | 23.527 | 23.373 | 17.906 | 23.256 |

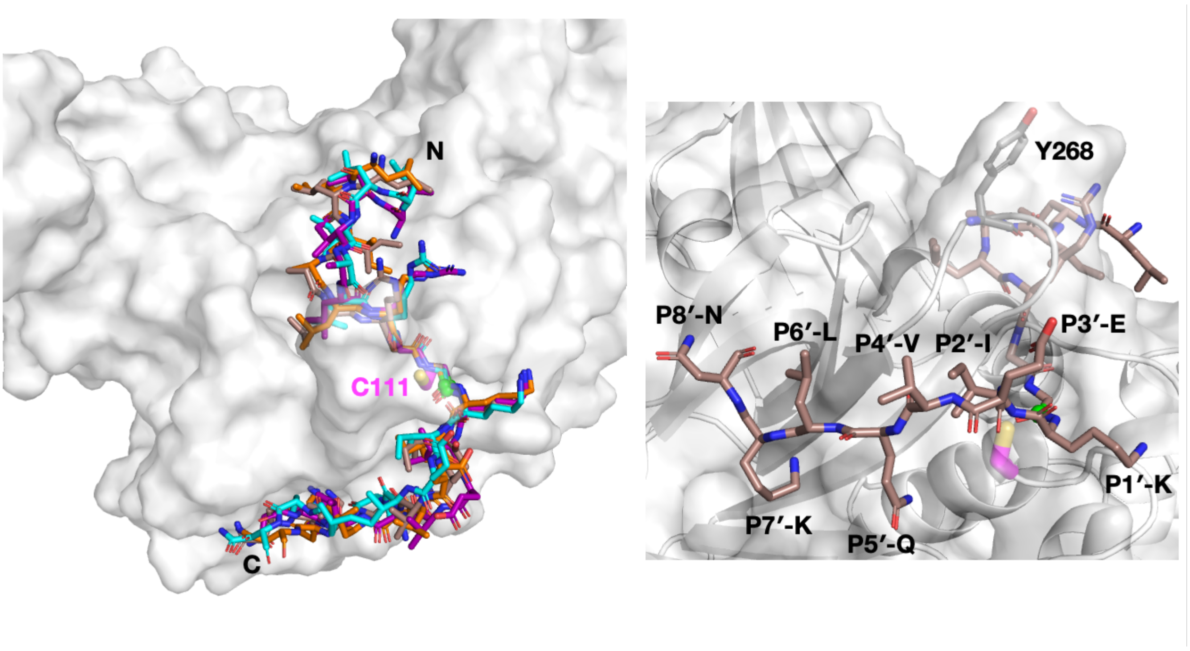

**Figure S38:** Docked conformations of PROS1-derived peptide **9** that passed the applied filter, *i.e.*, the four Cα atoms of P4-P1 have to be within 2 Å of the corresponding Cα atoms of VIR251[3]. The poses include ranked solutions 06 (brown), 21 (cyan), 58 (orange), and 97 (purple), with the general positioning of the terminals labeled. The P1 amide carbonyl carbon is shown as a green sphere. A magnified view of the P′ residues in solution 06 is shown, along with Tyr268 (grey sticks) in the BL2 loop.

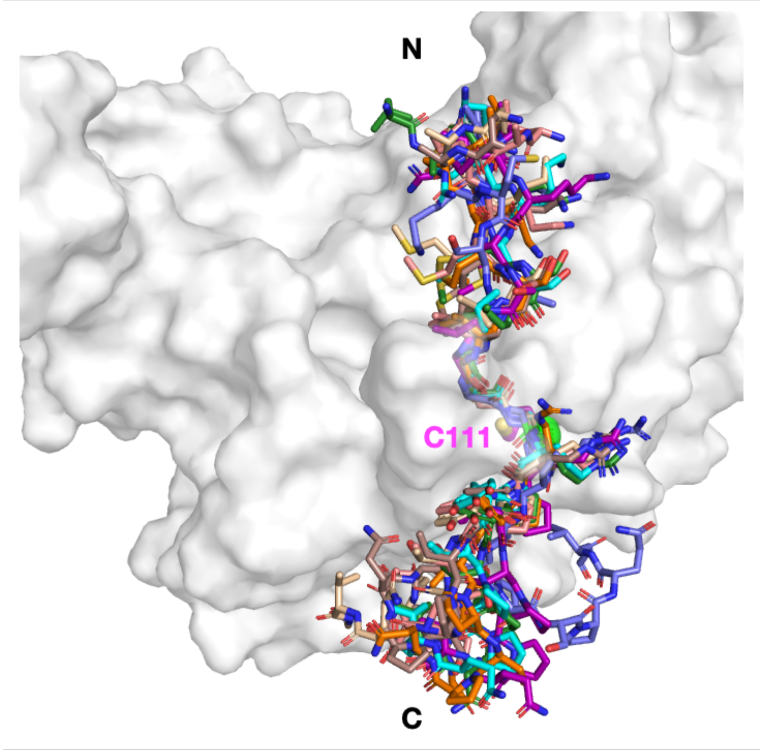

**Figure S39:** Docked conformations of ULK1-derived peptide **10** where P3-P1 Cα deviations were within 2 Å, but not P4. The poses include ranked solutions 09 (brown), 18 (cyan), 23 (orange), 44 (purple), 57 (dark green), 68 (slate), 72 (wheat), and 86 (salmon), with the general positioning of the terminals labeled. The P1 amide carbonyl carbon is shown as a green sphere.

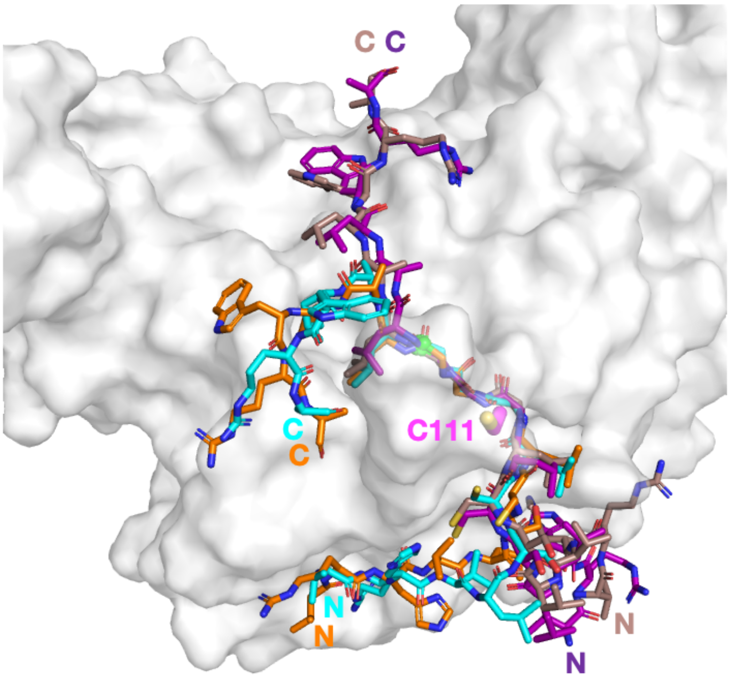

**Figure S40:** Docked conformations of IRF3-derived peptide **8** that are bound in reverse with the P2-Gly in S2 and P3-Gly in S1. The poses include ranked solutions 12 (brown), 42 (cyan), 49 (orange), and 86 (purple), with individual N- and C-terminals labeled. The P1 amide carbonyl carbon is shown as a green sphere.

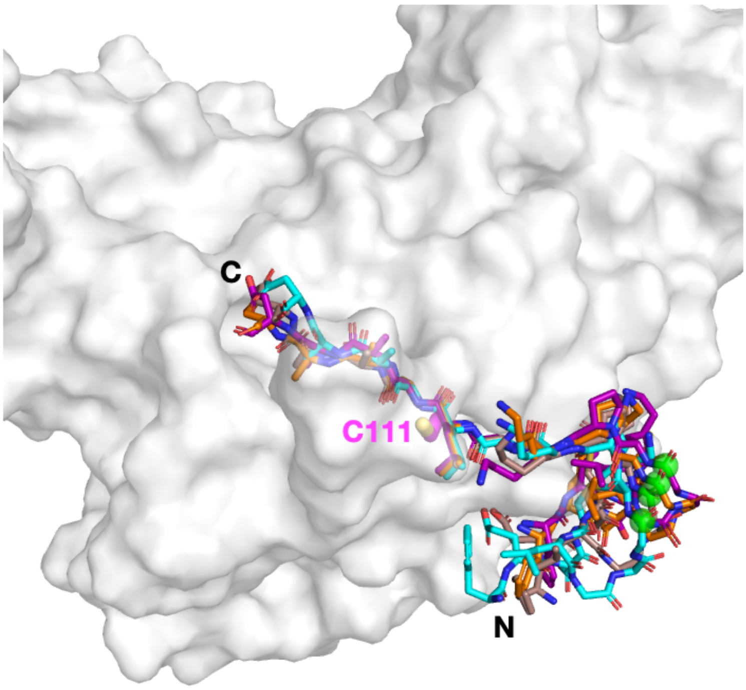

**Figure S41:** Docked conformations of ATG7-derived peptide **11** that are bound in reverse with the P5′-P8′ LAAA residues in the S1-S4 subsites, respectively. The poses include ranked solutions 02 (brown), 4 (cyan), 13 (orange), and 31 (purple), with the general positioning of the terminals labeled. The P1 amide carbonyl carbon is shown as a green sphere.

**Table S6**: Assignment of $PL^{pro}$ histidine protonation states from the structure of PDB 6WX4.[3] Neutral $N^{\delta}$-protonated, neutral $N^{\varepsilon}$-protonated, and doubly protonated histidines are named HID, HIE, and HIP respectively.

| HIS | Assignment | Reason |
|---|---|---|
| 17 | HIP | HE2 hydrogen bond (HB) donor to the Glu67 carboxylate (N-O 2.8 Å). Solvent exposed on ND1 side. p$K_a$ calculations (H++ and PROPKA)[9,10] consistently suggest double protonation at pH 7, with negatively charged residues Asp12 and Glu67 nearby, but no positively charged residues nearby. |
| 47 | HIE | ND1 HB acceptor for Ser49 sidechain OH (N-O 3.0 Å) or Ser49 backbone NH (N-N 3.1 Å). |
| 50 | HIE | ND1 HB acceptor for Tyr27 sidechain OH (N-O 2.9 Å) or His47 backbone NH (N-N 3.2 Å). |
| 73 | HIE | No clear HB partners nearby, assuming HIE. |
| 89 | HID | NE2 solvent exposed. ND1/HD1 HB donors to Ser85 backbone O (N-O 3.4 Å). |
| 175 | HID | ND1/HD1 HB donors to Tyr171 backbone O (N-O 3.0 Å). |
| 255 | HIE | NE2/HE2 HB donors to Lys279 backbone O (N-O 2.6 Å). |
| 272 | HIE/HIP | Part of the catalytic triad. HE2 HB donor to Asp286 (N-O 3.0 Å). ND1 side solvent exposed; protonation state likely depends on the charge state of Cys111.<br>Subsequent QM/MM-US calculations (**Figures S7-S11**) suggested HIP is favored. |
| 275 | HID | NE2 HB acceptor for Gln122 NE2 (N-N 2.8 Å), which in turn is a HB donor to backbone O of Leu118 (N-O 3.0 Å) and HB acceptor, via its OE1, for T277 sidechain OH (O-O 2.6 Å). |
